# Discovery and Engineering of a New BvCas12a Nuclease for Mammalian Genome Editing and Nucleic Acid Detection

**DOI:** 10.1101/2024.08.07.606956

**Authors:** Xuechun SHEN, Lei HUANG, Dan WANG, Chuan LIU, Ke CHEN, Baitao LI, Chen QI, Yuanqiang ZOU, Liang XIAO, Yuan JIANG, Hongxia LAN, Yue ZHENG

## Abstract

Cas12a is an RNA-guided endonuclease that has emerged as a powerful gene-editing tool. We have identified a novel Cas protein, BvCas12a, from *Butyricimonas virosa* with a 5’-TYTN protospacer adjacent motif (PAM). BvCas12a exhibits double-stranded DNA cleavage activity *in vitro* and genome editing activity in eukaryotic cells. Though the editing efficiency of BvCas12a is marginally lower than that of AsCas12a, the editing specificity of BvCas12a in eukaryotic cells is comparable or superior to that of AsCas12a. Moreover, BvCas12a exhibits substantial collateral activity and can detect HPV DNA effectively and accurately in conjunction with isothermal amplification, highlighting its potential in nucleic acid diagnostics. Furthermore, we have engineered BvCas12a to create two variants, BvCas12a-R (N549R, T606P) and BvCas12a-RVR (N549R, K555V, C559R, T606P). These variants recognize expanded 5’-YYN and 5’-YN PAM *in vitro*, respectively. Additionally, they exhibit higher editing activity than wild-type BvCas12a and recognize 5’-YYN PAM *in vivo* at all sites detected. In conclusion, we have identified a novel BvCas12a protein with high specificity and engineered two variants with broader PAM compatibility and improved genome editing efficiency. These findings offer a potent gene editing tool for application in scientific research, gene therapy, and nucleic acid diagnostics.

## Introduction

The CRISPR/Cas system, an acronym for Clustered Regularly Interspaced Short Palindromic Repeats and CRISPR-associated proteins, represents an adaptive immune mechanism prevalent among archaea and bacteria.^1^ Under the guidance of crRNA (CRISPR RNA) or sgRNA (single guide RNA), the Cas protein can specifically cleave the target sequences, causing a DSB (DNA double-strand break), with activity contingent on the presence of an appropriate PAM (Protospacer Adjacent Motif). In eukaryotic cells, DSBs can be repaired through either HDR (Homology-Directed Repair)^2^ or NHEJ (Non-Homologous End Joining)^3^ pathways, potentially resulting in gene knock-down^4^ or knock-in^5^ events during the repair process. The CRISPR/Cas system has evolved into a simple, efficient, accurate, and low-toxicity genome editing tool, and it can be easily used to edit targeting DNA sequences of interest by designing and employing specific gRNAs.^6^

In 2015, Zetsche *et al*. discovered a type V CRISPR gene editing system, with an effector protein guided by a single crRNA, known as CRISPR-Cpf1 (CRISPR from *Prevotella* and *Francisella* 1) or CRISPR-Cas12a.^7^ Compared to the well-characterized Cas9,^8^ Cas12a offers several benefits: 1) It only requires a single, shorter crRNA to target DNA. 2) Unlike Cas9, which recognizes a G-rich PAM at the 3′ end of the protospacer, Cas12a identifies a T-rich PAM at the 5′ end, broadening the scope of targetable sequences for Cas proteins. 3) CRISPR-Cas12a demonstrates *cis*- and *trans*-cleavage activities on ssDNA (single-stranded DNA), facilitating its use in nucleic acid diagnostics. These notable benefits have quickly established Cas12a as a preferred genome editing tool, leading to systematic research and successful application in gene therapy,^9–11^ genome editing,^12,13^ and *in vitro* molecular diagnostics.^14–16^

Despite the significant potential of Cas12a nucleases in genome editing, major obstacles still hinder their widespread application. First, most Cas12a proteins are restricted to recognizing T-rich PAM sequences, leading to target-dependent editing inefficiencies and constraining their utility in genome editing. Second, the potential safety concerns due to the unavoidable off-target effects must be addressed. Third, the patent protection of Cas12a proteins restricts their applications in commercial areas. Consequently, it’s critical to discover and characterize new Cas12a proteins or to modify existing ones to fulfill future research and commercial demands. The main strategies for enhancing the CRISPR/Cas system’s efficiency and precision include crRNA and protein engineering. Optimizing the structure of crRNA can increase its binding affinity to nucleases, thereby improving targeting activity and editing efficiency. Park *et al.*^17^ found that by extending the crRNA’s 5’ end, the gene editing efficiency of Cas12a is enhanced in both *in vitro* and *in vivo* settings. More precise transcription of crRNA and improved editing efficiency can be attained by appending tRNA^18^ or the self-cleaving ribozyme HDV^19^ to the 3’ end of crRNA. Su Bin Moon *et al.*^20^ observed that the 3’-terminal U-rich crRNA can significantly boost the indel efficiency of AsCas12a to a level nearly comparable with that of SpCas9. Engineered variants of the Cas12a protein have been developed through directed evolution, and rational design, broadening the scope of its applicability or improving its specificity and efficiency. Liyang Zhang *et al*.^21^ constructed a random amino acid mutation library through low-fidelity PCR, characterized each mutant’s identity and frequency via NGS sequencing, and screened for AsCas12a mutants under the selective pressure of antibacterial toxins. Consequently, they isolated variants of AsCas12a capable of recognizing the 5’TTTT PAM sequence. Feng Zhang *et al*.^22^ utilized the crystal structure of the AsCas12a-crRNA-target DNA complex to strategically mutate specific residues, thereby successfully identifying mutants with an expanded range of PAM recognition.

Here, we have functionally characterized a novel type V-A CRISPR-Cas12a protein, BvCas12a, derived from *Butyricimonas virosa*. BvCas12a recognizes 5’-TYTN PAM and efficiently cleaves dsDNA within a temperature range of 16–50°C. Additionally, we evaluated the gene editing activity and specificity of BvCas12a *in vivo*. We found that although BvCas12a has a slightly lower editing efficiency, its specificity in editing is on par with or possibly surpasses that of AsCas12a. Furthermore, we observed collateral cleavage activity of BvCas12a and demonstrated its potential in HPV (human papillomavirus) detection. Finally, we engineered two variants of BvCas12a with more relaxed PAM recognition and enhanced genome editing activity.

## Results

### 1. Characterization of BvCas12a from *Butyricimonas virosa*

A novel RNA-guided endonuclease BvCas12a, which is an ortholog of the Cas12a from the species *Butyricimonas virosa*, was discovered in CNGB (China National GeneBank) database.^23^ The BvCas12a protein consists of 1245 amino acids and shares 45.7% protein identity with LbCas12a and 33.1% with AsCas12a. BvCas12a possesses a complete RuvC-like endonuclease domain with characteristic active sites consistent with AsCas12a^24^ as predicted by AlphaFold2 (Figure S1A). So, we hypothesize that BvCas12a may serve as a functional RNA-guided endonuclease. Considering the evolutionary lineage of Cas12a family proteins and the conservation of direct repeat sequences, we predicted that the CRISPR-BvCas12a system includes a mature-crRNA with 19 nucleotides (nt) of the direct repeat (Figure S1B, C and D). To test the activity of BvCas12a with the predicted mature-crRNA, the *in vitro* dsDNA cleavage assay was performed using a dsDNA library containing 7 bp random sequences followed by a target sequence. The result demonstrated that the BvCas12a with a crRNA specific for the target sequence could cleave the dsDNA library *in vitro* (Figure S1E). To characterize the PAM sequences recognized by BvCas12a, we previously utilized our DocMF (DNB-based on-chip motif finding) platform,^25^ which showed that BvCas12a recognizes 5’-TYTN PAM efficiently *in vitro*, similar but more flexible to the 5’-TTTN PAM recognized by AsCas12a and LbCas12a.

**Figure S1.**
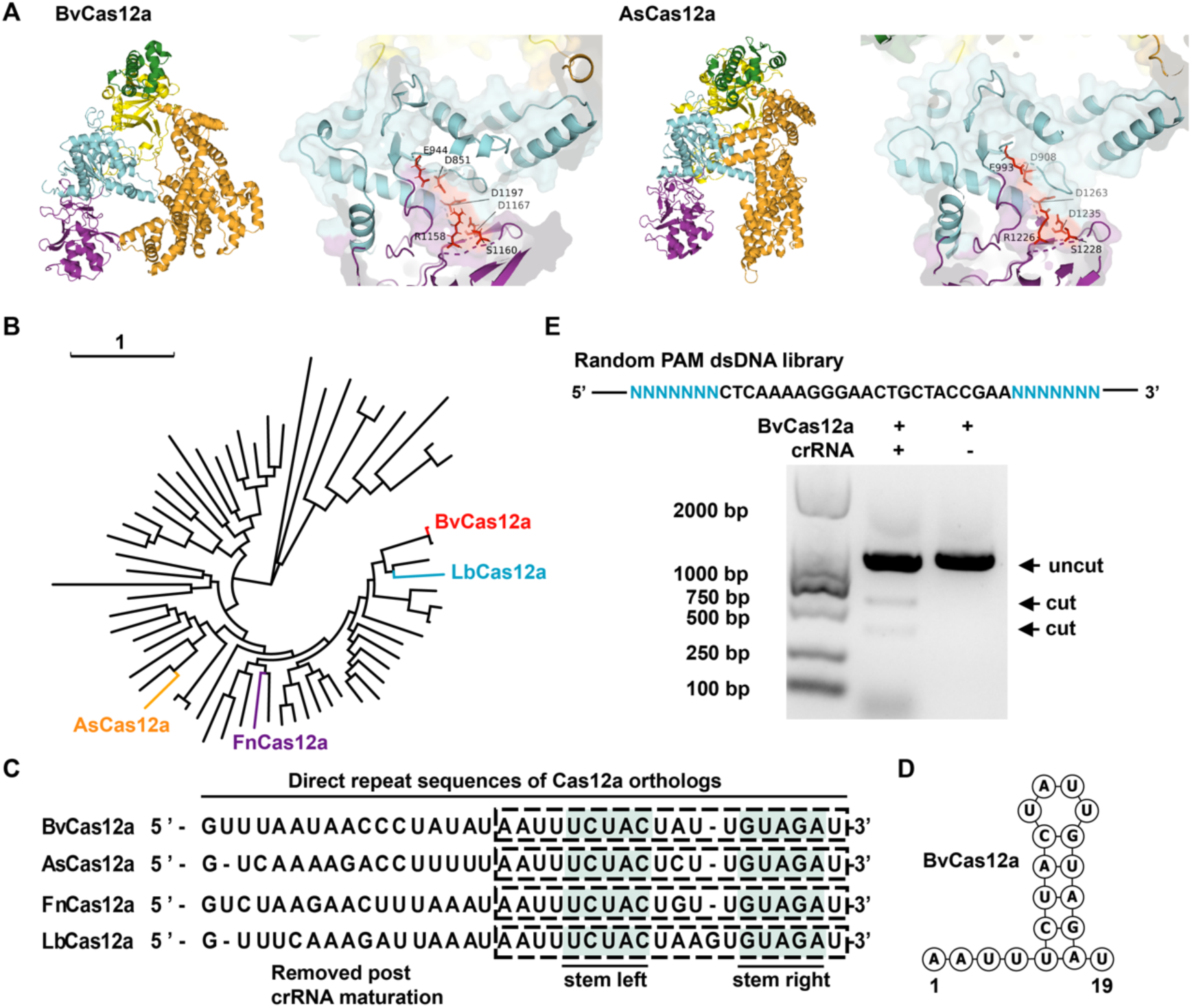
Characterization of BvCas12a and Its dsDNA Cleavage Activity *In Vitro*. (A) The predicted structure and active sites of BvCas12a, as inferred by AlphaFold2, are compared to those of AsCas12a (PDB ID: 5B43J. (B) A phylogenetic tree of type V-A effector proteins, with BvCas12a, AsCas12a, FnCas12a, and LbCas12a highlighted on the branches in red, orange, purple, and blue, respectively. (C) Alignment of the repeat sequences in the BvCas12a, AsCas12a, FnCas12a, and LbCas12a systems. The dashed boxes highlight the mature crRNA repeats, while the shaded areas illustrate the stem-loop structures. (D) A schematic diagram of the mature crRNA repeat structure of BvCas12a. (E) An *in vitro* cleavage assay utilizing a random-PAM dsDNA library. The linear dsDNA PAM library, containing a target site bordered by seven random nucleotides at both the 3’ and 5’ ends, was treated with BvCas12a in the presence of the corresponding crRNA. The agarose gel image displays the electrophoretic separation of cleavage products derived from the random PAM library following a 30-minute incubation with the BvCas12a effector complex

Considering the influence of spacer length on the architecture of the RNP complex and the editing efficiency of Cas12a,^26^ we explored the necessary guide sequence length for BvCas12a-mediated DNA cleavage. An *in vitro* cleavage assay on supercoiled plasmids, targeting the ampicillin coding region with the BvCas12a crRNA, revealed that a minimum of 16 nt in the guide sequence is essential for significant DNA cleavage, with an optimal guide sequence length of at least 20 nt (Figure 1A). Previous research has observed that the 3’-terminal U-rich tail of crRNA can enhance the activity of AsCas12a both *in vitro* and *in vivo*.^27^ Our result also confirmed that BvCas12a with the 3’-terminal U4AU4 tail of crRNA has optimal dsDNA cleavage efficiency *in vitro* (Figure 1A). Further, the optimal temperature range for enzyme activity is crucial for evaluating the suitability of Cas12a nucleases for various applications.^28^ We conducted a BvCas12a activity assay at divergent temperatures using a linear DNA fragment containing the HPV16 fragment as the substrate. The result showed that the substrate was cleaved by BvCas12a efficiently within a temperature spectrum of 16–50°C, and the peak activity was at approximately 46°C (Figure 1B). This suggests that BvCas12a is adaptable for genome editing in various organisms, such as *Xenopus* (20-24°C), *Drosophila* (20-25°C)*, Saccharomyces* (25-28°C), animals, and plants.

**Figure 1.**
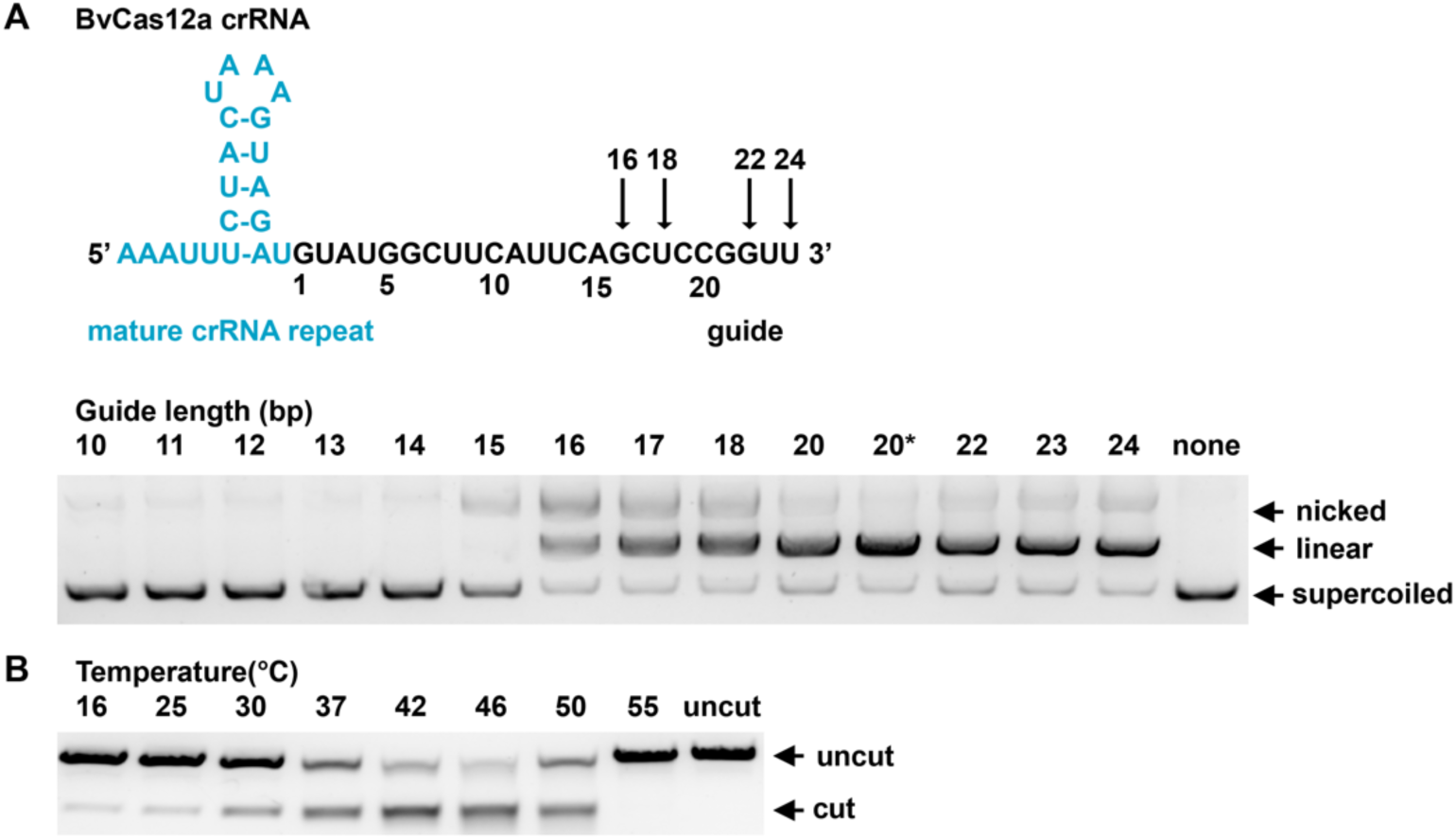
Characterization of the Optimal Guide Length and Temperature for BvCas12a Activity*In Vitro*. (A) Effect of guide length on BvCas12a cleavage activity *in vitro*. BvCas12a and crRNAs with varying guide lengths were utilized in a supercoiled plasmid cleavage assay. The asterisk indicates a crRNAfeaturing a 3’ U4AU4 extension at its 3’ end.^27^ (B) DNA cleavage activity of BvCas12a at different temperatures *in vitro*. BvCas12a and its crRNA with a 23 bp guide were employed in a linear DNA fragment cleavage assay.

### 2. BvCas12a facilitates genome editing in mammalian cells

Previous results have demonstrated that BvCas12a possesses RNA-guided endonuclease activity for dsDNA cleavage *in vitro*. To explore the potential of BvCas12a as a genome-editing tool in human cells, we performed genome editing experiments in HEK293T cells using a mammalian expression plasmid containing a human-codon-optimized BvCas12a and crRNA expression cassette regulated by the RNA polymerase Ⅲ U6 promoter (Figure S2A). Initial detection of endogenous indels was accomplished using the T7E1 (T7 endonuclease 1) assay. The results revealed that BvCas12a can cause indels in three target sites within the *AAVS1* gene, demonstrating that BvCas12a enables genome editing in mammalian cells (Figure S2A).

Although we have previously assessed the optimal guide sequence length for BvCas12a-mediated DNA cleavage *in vitro*, further experiments are required to ascertain the ideal crRNA guide sequence length for effective genome editing *in vivo*. Prior studies have demonstrated that self-cleaving ribozymes can enhance the processing of crRNA transcripts to yield the exact guide molecule, thereby improving the gene-editing efficiency of the CRISPR-Cas12a system.^22^ Accordingly, we constructed a mammalian expression plasmid of BvCas12a by inserting an HDV ribozyme at the 3’ end of the crRNA (Figure S2B). Subsequent experiments were performed to determine the optimal guide length of crRNA for *in vivo* genome editing. According to the amplicon-seq results,^29^ BvCas12a showed high editing efficiency with a spacer length of 20∼24 nt *in vivo*, remaining relatively stable at approximately 60% indel rate at the A2 target site of the AAVS1 gene detected. However, the efficiency decreased dramatically when the spacer length was 18 nt or less, around 6% indel rate at the same target site with an 18 nt spacer, and undetectable with a 16 nt spacer (Figure 2A). Consequently, the 23 bp spacer was selected for further experimental work.

**Figure 2.**
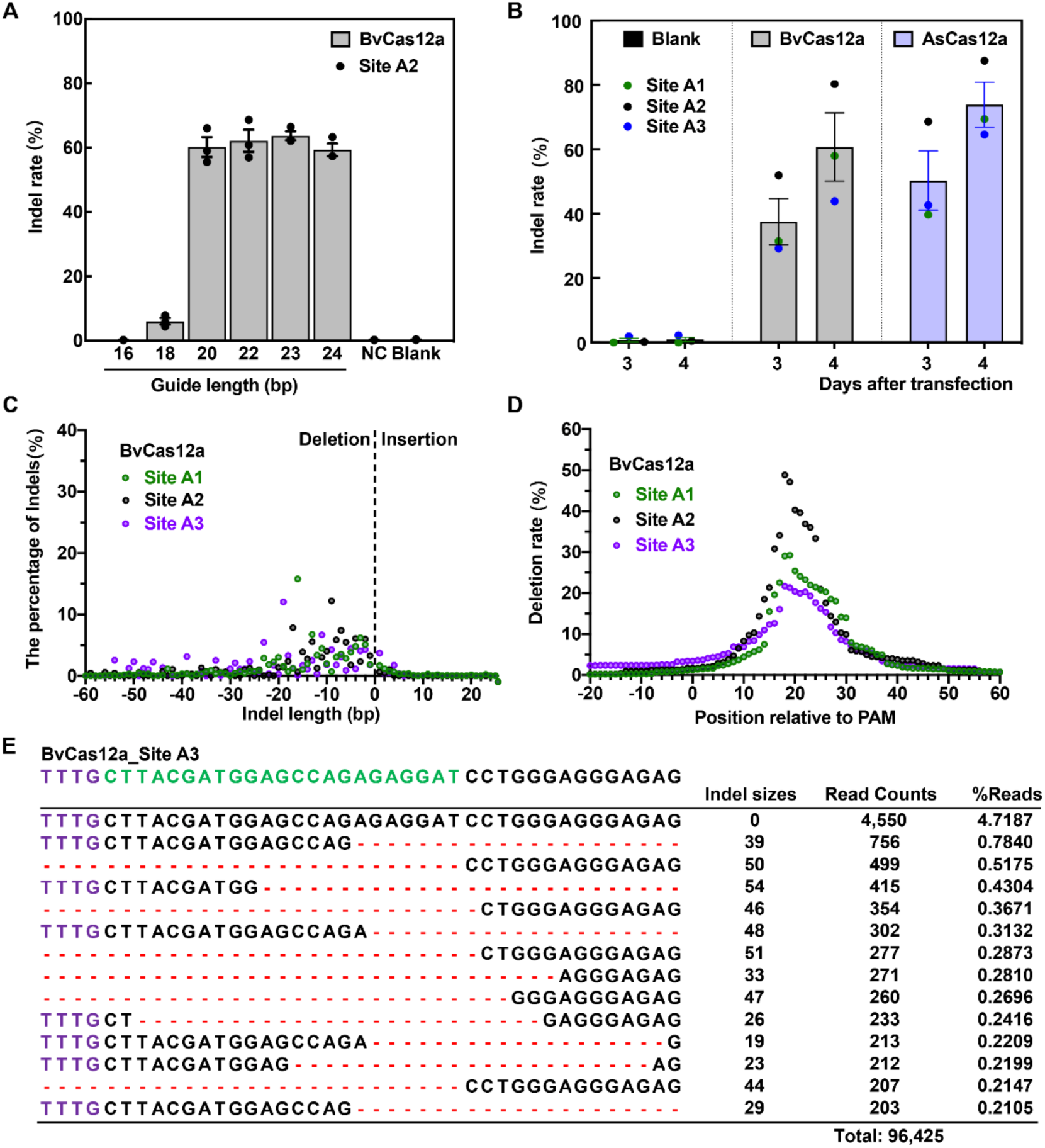
Characterization of BvCas12a Activity in Mammalian Cells. (A) Impact of crRNA guide length on indel efficiency induced by BvCas12a *in vivo*. HEK293T cells were transfected with plasmids encoding BvCas12a and crRNAs of various guide lengths. The indel rates were evaluated via targeted amplicon sequencing, mean values are presented with the standard error of the mean (SEM), based on n=3 independent experiments. (B) Editing efficiency comparison between BvCas12a and AsCas12a *in vivo*. The indel rates were evaluated via targeted amplicon sequencing. Mean indel rates with corresponding SEMs were obtained from BvCas12a and AsCas12a-mediated genome editing events at three AAVS1 target sites in HEK293T cells, three to four days post-transfection. (C) Indel length distribution in genome editing mediated by BvCas12a. The data represent the percentage of aligned reads that exhibit an insertion or deletion at the specified length of three AAVS1 target sites. (D) Deletion rate at each nucleotide position during genome editing with BvCas12a. The data illustrate the rate of aligned reads with a deletion at the indicated nucleotide positions in three AAVS1 target sites. In the TTTN PAM sequence, ’N’ denotes position 0. (E) The typical genome editing (deletion) of BvCas12a at target site A3. The PAM and target sequences of A3 site were indicated in purple and green, respectively.

**Figure S2.**
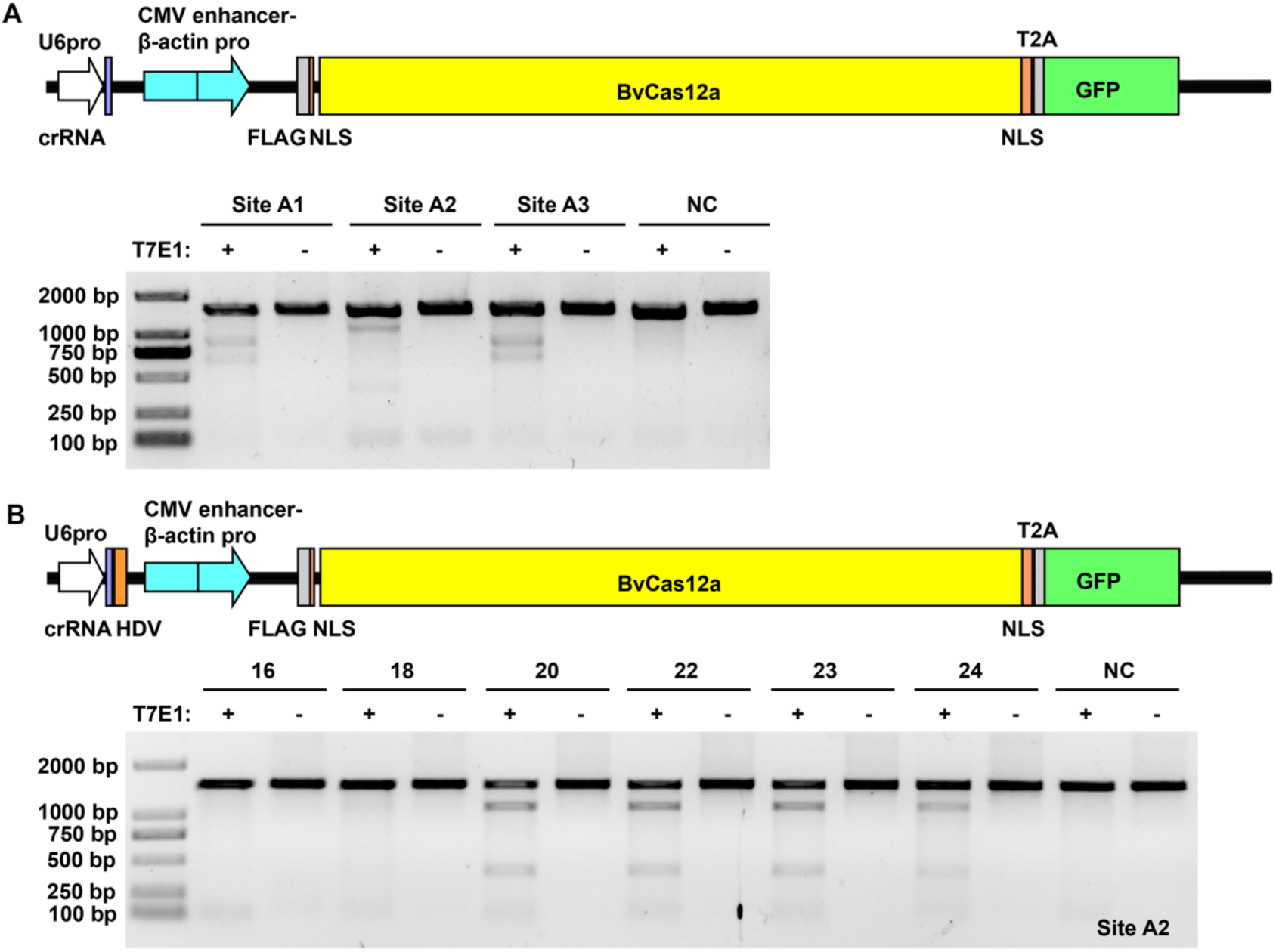
Characterization of BvCas12a Genome Editing by T7 Endonuclease I (T7E1) Assay. (A) T7E1 assay of BvCasI 2a-mediated gene editing in HEK293T cells. The upper panel presents a schematic map of the BvCasI 2a plasmid utilized for human genome editing, while the lower panel displays the T7E1 assay results at three AAVS1 sites, each with a 5’ TTTN PAM sequence. (B) Evaluation of the optimal guide RNA length for BvCasI 2a-mediated editing in HEK293T cells. The upper panel illustrates a schematic map of the modified BvCasI 2a plasmid, which includes an HDV ribozyme for precise gRNA production, and the lower panel shows the T7E1 assay outcomes for genome editing samples with guide RNAs ranging from 16 bp to 24 bp at the AAVS1 site A2.

To systematically assess the genome editing capabilities of BvCas12a, we performed a genome editing experiment *in vivo*, comparing its efficiency to AsCas12a. The next-generation sequencing (NGS) results showed that BvCas12a exhibits substantial, albeit slightly lower, editing efficiency than AsCas12a, with 38% indels at three days post-transfection, rising to 61% by day four (Figure 2B). BvCas12a mainly caused deletions, with most ranging from 1 to 60 base pairs, concentrated 16-18 base pairs downstream from the PAM (Figure 2C to 2E). Although BvCas12a’s editing activity is slightly lower, its pattern mirrors AsCas12a’s (Figure S3). In addition to deletions, BvCas12a induced a small percentage of insertions, typically less than 1% (Figure 2C). This indicates that while BvCas12a favors deletions, it can also cause insertion at a much lower frequency.

**Figure S3.**
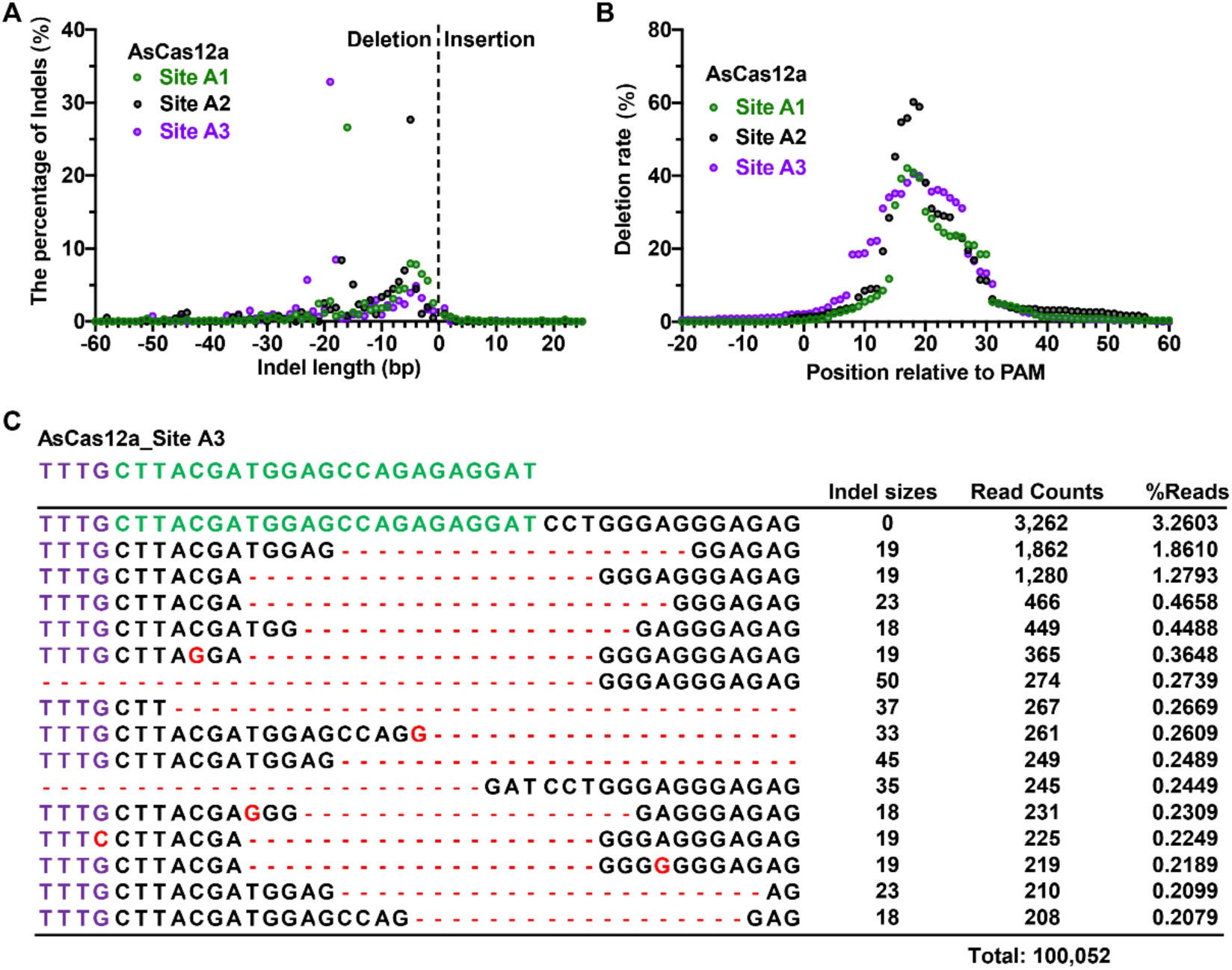
Characterization of AsCas12a genome editing in Mammalian Cells. (A) Indel length distribution in genome editing mediated by AsCas12a. The data represent the percentage of aligned reads that exhibit an insertion or deletion at the specified length of three AAVS1 target sites. (B) Deletion rate at each nucleotide position during genome editing with AsCas12a. The data illustrate the rate of aligned reads with a deletion at the indicated nucleotide positions in three AAVS1 target sites. In the TTTN PAM sequence, ’N’ denotes position 0. (C) The typical genome editing (deletion) of AsCas12a at target site A3. The PAM and target sequences of A3 site were indicated in purple and green, respectively.

To evaluate the genome editing specificity of BvCas12a, the on-target and off-target indel frequencies were investigated for three targets of the AAVS1 gene. Initially, potential off-target sites were predicted through in-silico analysis using the tool Cas-OFFinder,^30^ which considered up to five base mismatches and required the presence of a 5’-TTTN PAM sequence. This led to the identification of nineteen off-target sites for the A1 and A3 targets, and sixteen for the A2 target (Figure S4B, S4D, and S4F). Subsequent PCR amplification of on-target and off-target regions allowed for the estimation of indel frequencies at each site through targeted amplicon sequencing. The NGS amplicon library was prepared at all target sites and the predicted off-target sites. The sequencing results from the NGS amplicon library indicated indel frequencies of 19.4%, 31.6%, and 87.9% at the A1, A2, and A3 target sites, respectively, for BvCas12a (Figure S4A, S4C, and S4E). These efficiencies closely match those achieved by AsCas12a, with corresponding indel frequencies of 27.1%, 46.5%, and 95.1% (Figure S4A, S4C, and S4E). In terms of off-target activity, both BvCas12a and AsCas12a demonstrated indel rates below 1% at all identified sites, except for A1-OFF8, A1-OFF16, and A2-OFF1, which were attributed to background indels (Figure S4A, S4C, and S4E). Moreover, the detected off-target indel frequencies of all off-target sites for BvCas12a and AsCas12a exhibited no statistically significant differences when compared to control samples, with the sole exception being at the A3-OFF8 site (Figure S4A, S4C, and S4E). Notably, the AsCas12a protein showed a significant off-target effect at the A3-OFF8 site, with an indel frequency of 0.90%, a phenomenon not observed with BvCas12a and control (Figure 3A). Further, the detailed editing of AsCas12a at the A3-OFF8 off-target site was analyzed and the results showed that AsCas12a predominately caused small deletion (less than 20 bp) in the sites between +7 to +35 downstream of PAM, indicating that it’s a true off-target site forAsCas12a (Figure 3B and 3C). Based on these findings, we predict that the editing specificity of BvCas12a in eukaryotic cells is comparable to or even better than that of AsCas12a.

**Figure 3.**
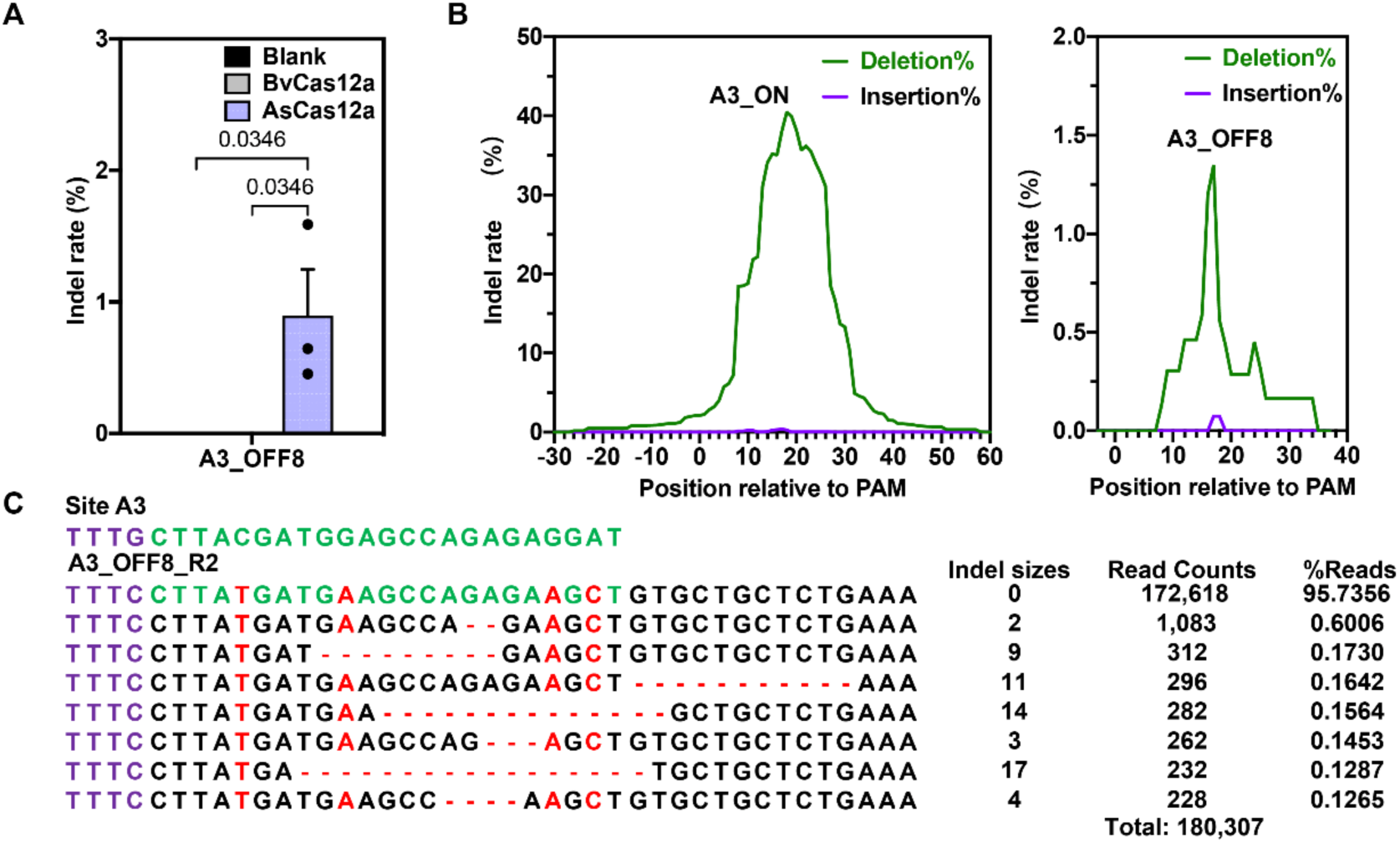
The Genome Editing Specificity Comparison of BvCas12a with AsCas12a in Mammalian Cells. (A) The genome editing specificity comparison of BvCas12a with AsCas12a. The AsCas12a protein showed a significant off-target effect at the A3-OFF8 site. Indel frequencies at predicted off-target sites were evaluated via targeted amplicon sequencing. The mean values with standard error of the mean (SEM) from three biological replicates are depicted. A two-way ANOVA followed by Tukey’s multiple comparison test was used to assess statistical significance of all detected sites, and sites with significant differences are shown. (B) The typical indel rates distribution of genome editing at each nucleotide position of A3 site and its off-target site A3_OFF8 with AsCas12a. In the TTTN PAM sequence, ’N’ denotes position 0. (C) The typical off-target editing (deletion) of AsCas12a at off-target site A3_OFF8. The PAM sequences were indicated in purple

**Figure S4.**
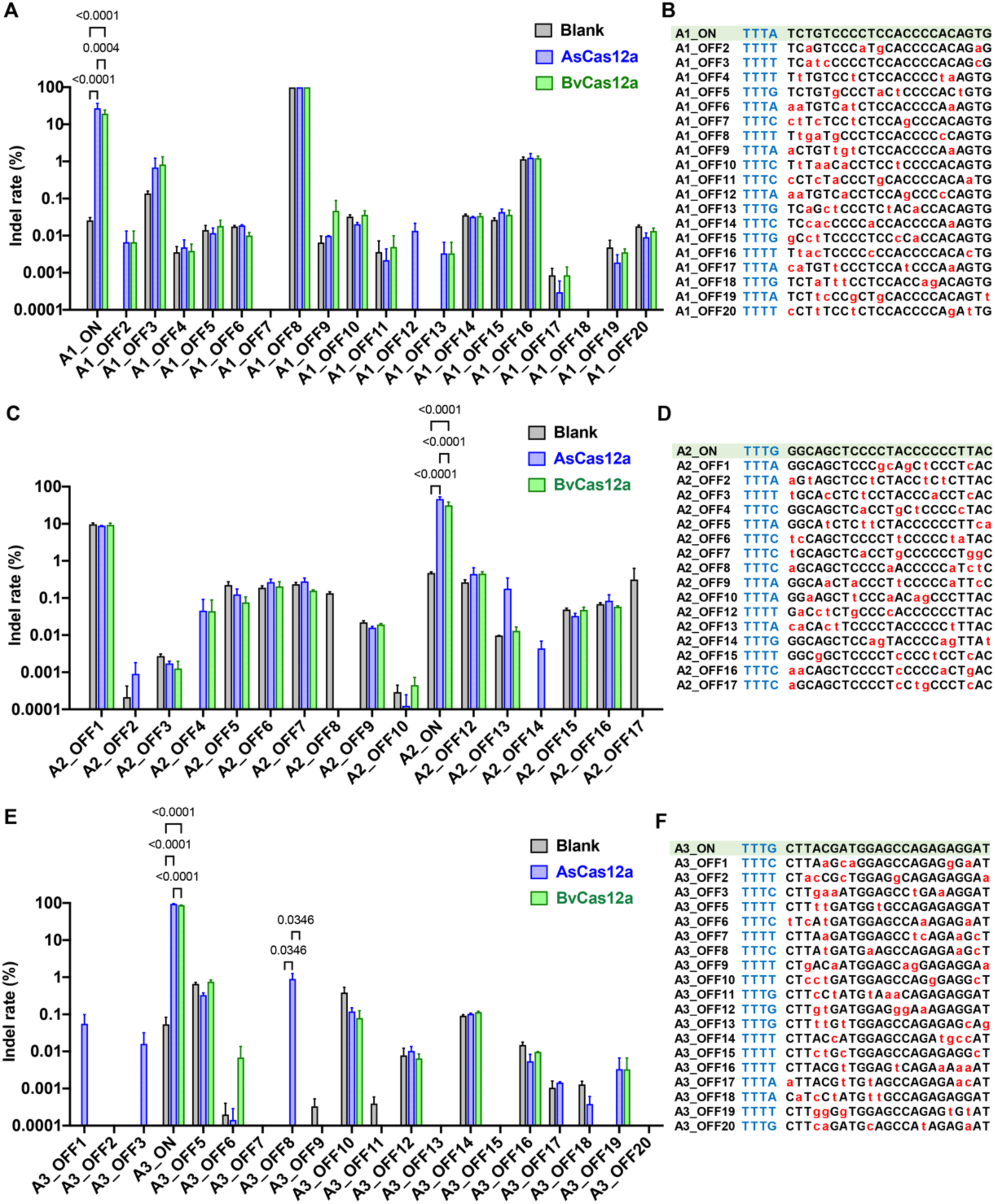
Comparison of Off-target Mutations induced by BvCas12a and AsCas12a. (A, C, and E) The on-target (ON) and off-target (OFF) indel rates (%) of BvCas12a and AsCas12a at three AAVS1 target sites, namely A1 site (A), A2 site (C), and A3 site (E), are presented on a logarithmic scale. Indel frequencies at predicted off-target sites were evaluated via targeted amplicon sequencing. The mean values with standard error of the mean (SEM) from three biological replicates are depicted. A two-way AN OVA followed by Tukey’s multiple comparison test was used to assess statistical significance. (B, D, and F) The off-target sites predicted for the three AAVS1 target sequences. The off-target sites of target sequences A1, A2, and A3 were predicted using Cas-OFFinder, allowing no more than five base mismatches and including a PAM sequence of 5’-TTTN. The protospacer-adjacent motif (PAM) sequences and mismatched bases between the target and off-target sequences are highlighted in blue and red, respectively.

### 3. BvCas12a has *trans*-cleavage activity on ssDNA

Previous research has shown that collateral activity is common among Cas12a proteins.^31^ In our study, we examined the collateral activity of BvCas12a to assess its potential for molecular diagnostic applications. The early detection and differentiation of human papillomaviruses (HPVs), particularly high-risk strains like HPV16 and HPV18, is crucial for the prevention and management of cervical cancer. To evaluate BvCas12a’s capacity to specifically recognize and distinguish HPV dsDNA viruses, particularly HPV16 and HPV18, we initially crafted two crRNA specifically targeting these viruses based on previously published studies.^14^ Subsequently, we incubated the BvCas12a-crRNA complex with target DNA fragments from HPV16 or HPV18 and ssDNA-FQ (fluorescein-quencher) probes at 37℃ and monitored the fluorescence changes every five minutes. The results showed that only the reactions with crRNAs matching their respective target generated a fluorescence signal, confirming that BvCas12a possesses collateral activity and is capable of accurately identifying HPV16 and HPV18 using specific crRNA sequences (Figure 4A).

**Figure 4.**
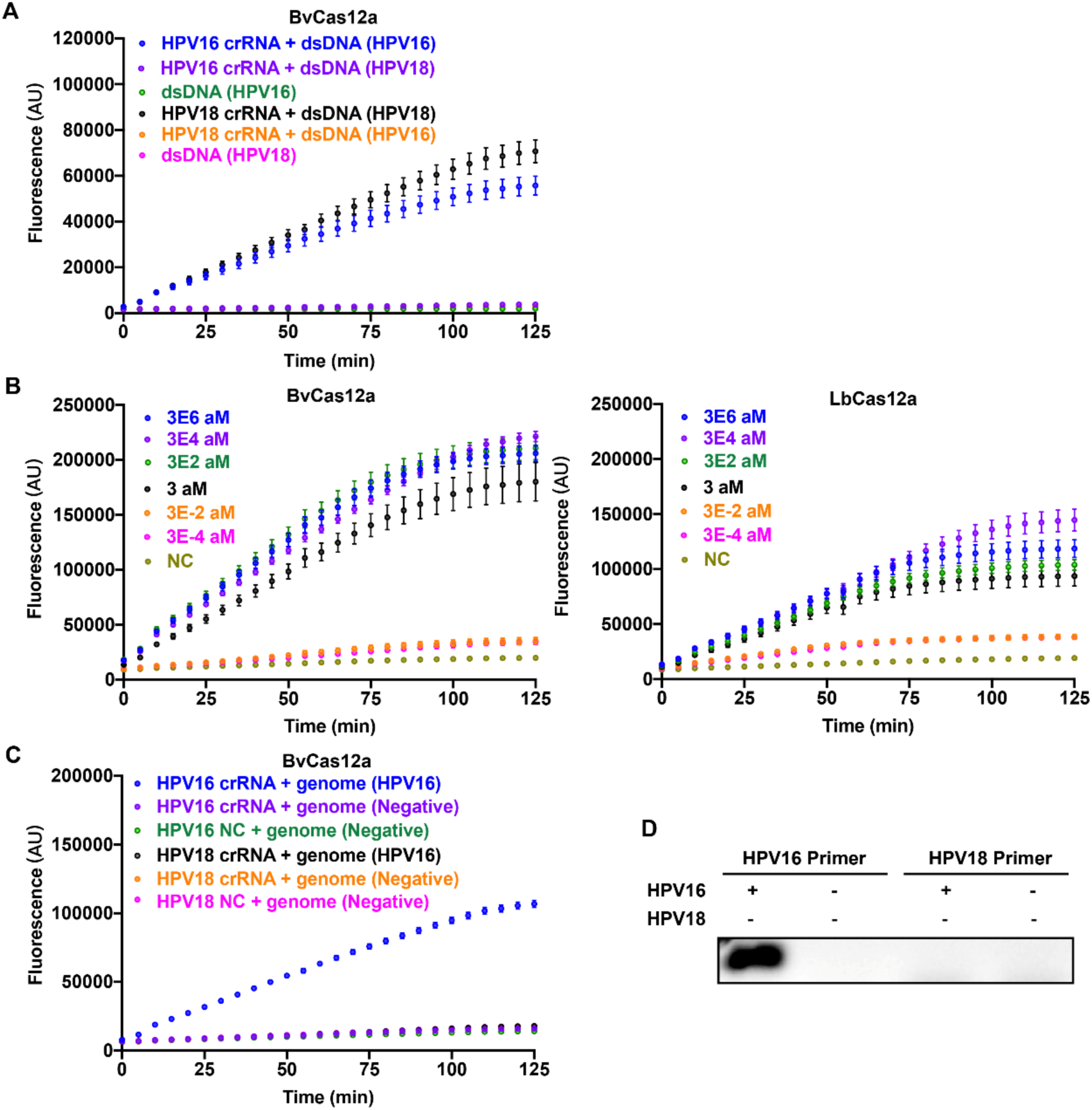
BvCas12a Demonstrates Collateral Cleavage Activity and Facilitates the Differentiation of HPV Types 16 and 18. (A) Fluorescence curves for BvCas12a-mediated ssDNAtrans-cleavage activated by PCR fragments of HPV16 or HPV18. with or without the specified crRNA. Error bars represent the mean ± SEM based on n=3 replicates. (B) BvCas12a demonstrates detection sensitivity comparable to LbCas12a, achieving attomolar detection limits in conjunction with isothermal amplification RPA. A range of diluted HPV16 plasmids were used as samples for detection. Error bars represent the mean ± SEM based on n=3 replicates. (C) Fluorescence curves for BvCas12a-mediated genomic detection of HPV16 in combination with RPA. Error bars represent the mean ± SEM based on n=3 replicates. (D) PCR vadiates the presence of HPV16 in genomic samples utilized in Figure 4C. The PCR amplification was performed with primers specific to HPV16 and HPV18.

Furthermore, we combined isothermal amplification via recombinase polymerase amplification (RPA)^32^ with BvCas12a to enhance nucleic acid detection sensitivity. We began by determining the detection limit using a series of diluted HPV16 plasmids and the assay was conducted following the previously established DETECTR protocol.^14^ The fluorescence curve of the assay confirmed that the sensitivity of the BvCas12a combined with RPA achieved attomolar levels, on par with that of LbCas12a (Figure 4B). Subsequently, we assessed BvCas12a’s proficiency in specifically detecting HPV16-positive human genomic samples. The fluorescence curve showed that BvCas12a only combined with HPV16-specific crRNA and HPV16 positive genomic sample elicited a significant fluorescence response, while the control lacking either the HPV16 target or specific crRNA exhibited merely background fluorescence (Figure 4C). Additionally, PCR analysis of the detected samples corroborated the results of the DETECTR assay mediated by BvCas12a (Figure 4D). Collectively, these findings suggest that BvCas12a has the potential to be developed for molecular diagnostics, offering high sensitivity and specificity.

### 4. BvCas12a variants present relaxed PAM and better activity

To enhance the editing capabilities and overcome the PAM restrictions of BvCas12a, we have engineered mutants of BvCas12a. These modifications focused on sites predicted to influence substrate binding and PAM recognition, guided by insights from the structures and engineered variants of AsCas12a^22^ and LbCas12a^33^, which have demonstrated altered PAM preferences. As a result, we generated two BvCas12a variants: BvCas12a-R (N549R, T606P) and BvCas12a-RVR (N549R, K555V, C559R, T606P).

The BvCas12a variants successfully cleaved the dsDNA library within a 7 bp random 5 ’-PAM, demonstrating that they retain the activity to cleave dsDNA *in vitro* (Figure S5A). Furthermore, our analysis revealed that BvCas12a-R recognized 5’YYN PAM, whereas BvCas12a-RVR recognized 5’YN PAM, in contrast to the 5’TYTN PAM recognized by the wild-type BvCas12a *in vitro* using the DocMF platform (Figure 5A). This indicates that the PAM preferences for these variants are more flexible *in vitro*. We also conducted *in vitro* cleavage assays on linear DNA fragments with identical target sequences but variable PAM sequences. The results confirmed that both BvCas12a-R and BvCas12a-RVR variants, particularly BvCas12a-RVR, can cleave dsDNA with 5’ AACA, AATA, and ACAA PAM targets, capabilities not observed with the wild-type BvCas12a *in vitro* (Figure S5B). This further highlights the broader PAM compatibility of the variants. Activity assays in HEK293T cells demonstrated that BvCas12a-R and BvCas12a-RVR exhibited enhanced activity for 5’ YYYN PAM targets detected, with a notable increase for non-T-rich PAM targets (Figure 5B to 5E). These findings support the conclusion that the BvCas12a-R and BvCas12a-RVR variants have higher gene editing activity and less stringent PAM requirements.

**Figure 5.**
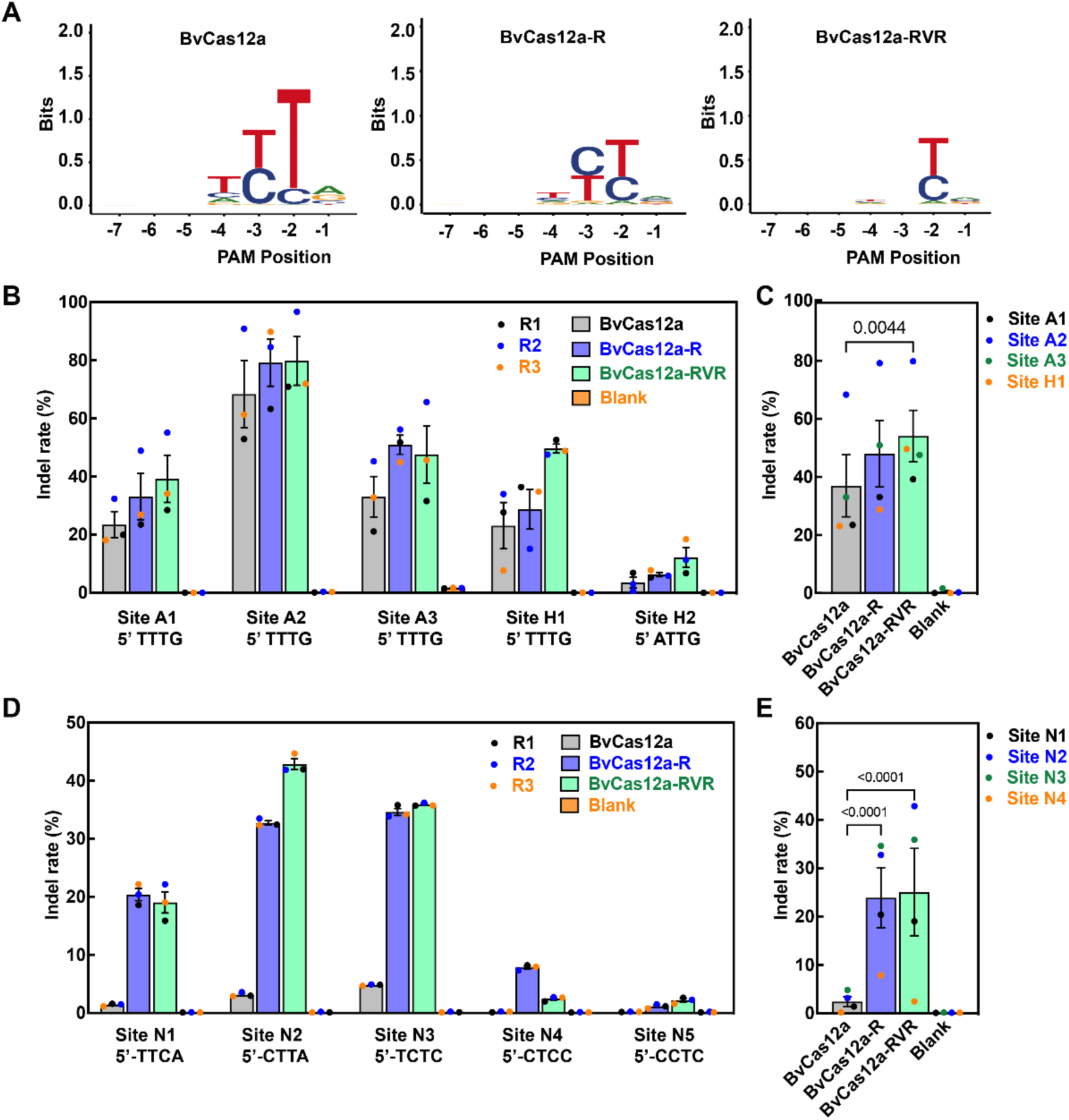
Characterization of BvCas12a Variants with Altered PAM Specificities. (A) PAM sequences for BvCas12a and its variants. The preferred PAM sequences were generated from the random 7-nt PAM library detected by DocMF platform. (B and C) Comparison of the editing activity of BvCas12a-WT/R/RVR at target sites of 5’-T rich PAMs in HEK293T cells. The editing activity of BvCas12a-WT/R/RVR in five different target sites were obtained by three independent experiments (8) and BvCas12a variants showed higher editing efficiency with 5’-TTTN PAM (C).The data obtained from various experimental repeats (B) and different sites (C) are presented with distinct colors. The indel rates of BvCa12a and its mutants in site H2 shown no signficant difference with control. A two-way ANOVA multiple comparisons with Tukey’s multiple comparison test was performed to evaluate statistical significance. (D and E) Comparison of the editing activity of BvCas12a-WT/R/RVR at target sites of 5’-C rich PAMs in HEK293T cells. The editing activity of BvCas12a-WT/R/RVR in five different target sites were obtained by three independent samples (D) and BvCas12a variants showed higher editing efficiency with 5’-YYYN PAM (E). The data acquired from various samples (D) and different sites (E) are presented with distinct colors. The indel rates of BvCa12a and its mutants in site NS shown no signficant difference with control. A two-way ANOVA multiple comparisons with Tukey’s multiple comparison test was performed to evaluate statistical significance.

To further evaluate the genome editing specificity of BvCas12a variants, we performed targeted amplicon sequencing on three targets within the AAVS1 gene and their associated off-target sites, which were predicted by Cas-OFFinder, as in previous studies (Figure S4B, S4D, and S4E). The sequencing of the NGS amplicon library revealed that the indel frequencies for BvCas12a, BvCas12a-R, and BvCas12a-RVR were 7.4%, 17.7%, and 19.6% at the A1 target site, 38.2%, 39.9%, and 30.7% at the A2 target site, and 84.9%, 94.8%, and 96.0% at the A3 target site, respectively. These indel rates were determined without accounting for the transfection rate, and they are consistent with the detection of off-target activity. Concerning off-target activity, BvCas12a and its variants, BvCas12a-R and BvCas12a-RVR, showed indel rates below 1% at all identified off-target sites, except for A1-OFF8, A1-OFF16, A2-OFF1, and A3-OFF10. Furthermore, the off-target indel frequencies detected across all sites for BvCas12a and its variants did not exhibit statistically significant differences compared to control samples (Figure S6A-S6C). Based on these results, we predict that the editing specificity of the BvCas12a variants, BvCas12a-R and BvCas12a-RVR, in eukaryotic cells is on par with that of the wild-type.

**Figure S5.**
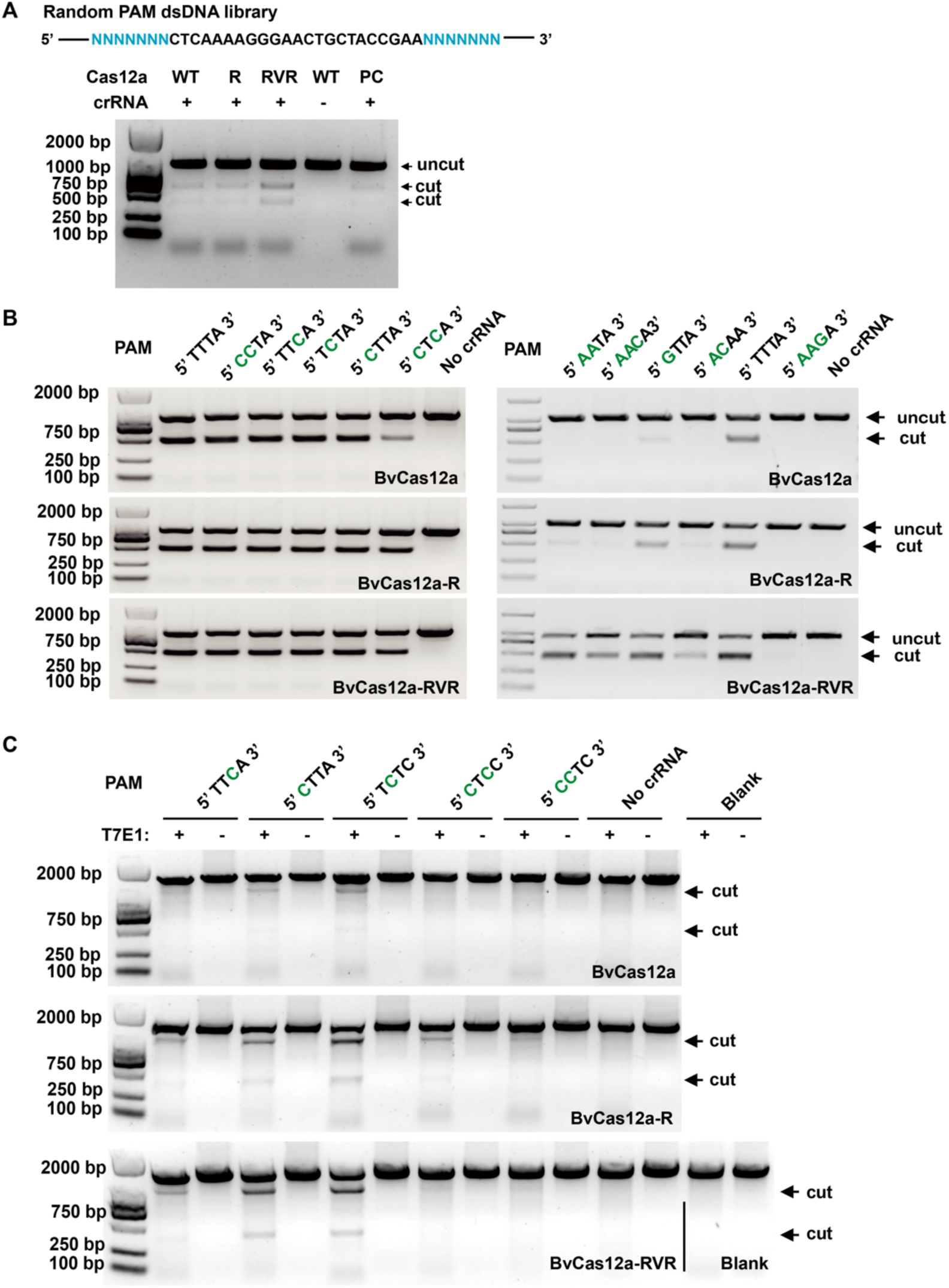
*In vitro* and *in vivo* Cleavage Activities of BvCas12a and Its Variants. (A) *In vitro* cleavage activity assay of BvCas12a and its variants using a random PAM dsDNA library. The linear dsDNA PAM library, containing a target site flanked by seven random nucleotides at both the 3’ and 5’ ends, was incubated with the BvCas12a-WT/R/RVR effector complex to assess the dsDNA cleavage activity *in vitro*. The LbCas12a used as positive control (PC). (B) *In vitro* cleavage activity assay of BvCas12a and its variants targeting the same sequence with various PAMs. The DNA fragments with varied PAMs and consensus target sequence (5’-GGCCTGTGTAGGTGTTGAGGTAG-3’) were obtained by PCR. (C) T7E1 assay to detect the gene editing activity of BvCas12a and its variants at different genomic target sites with diverse PAM sequences.

**Figure S6.**
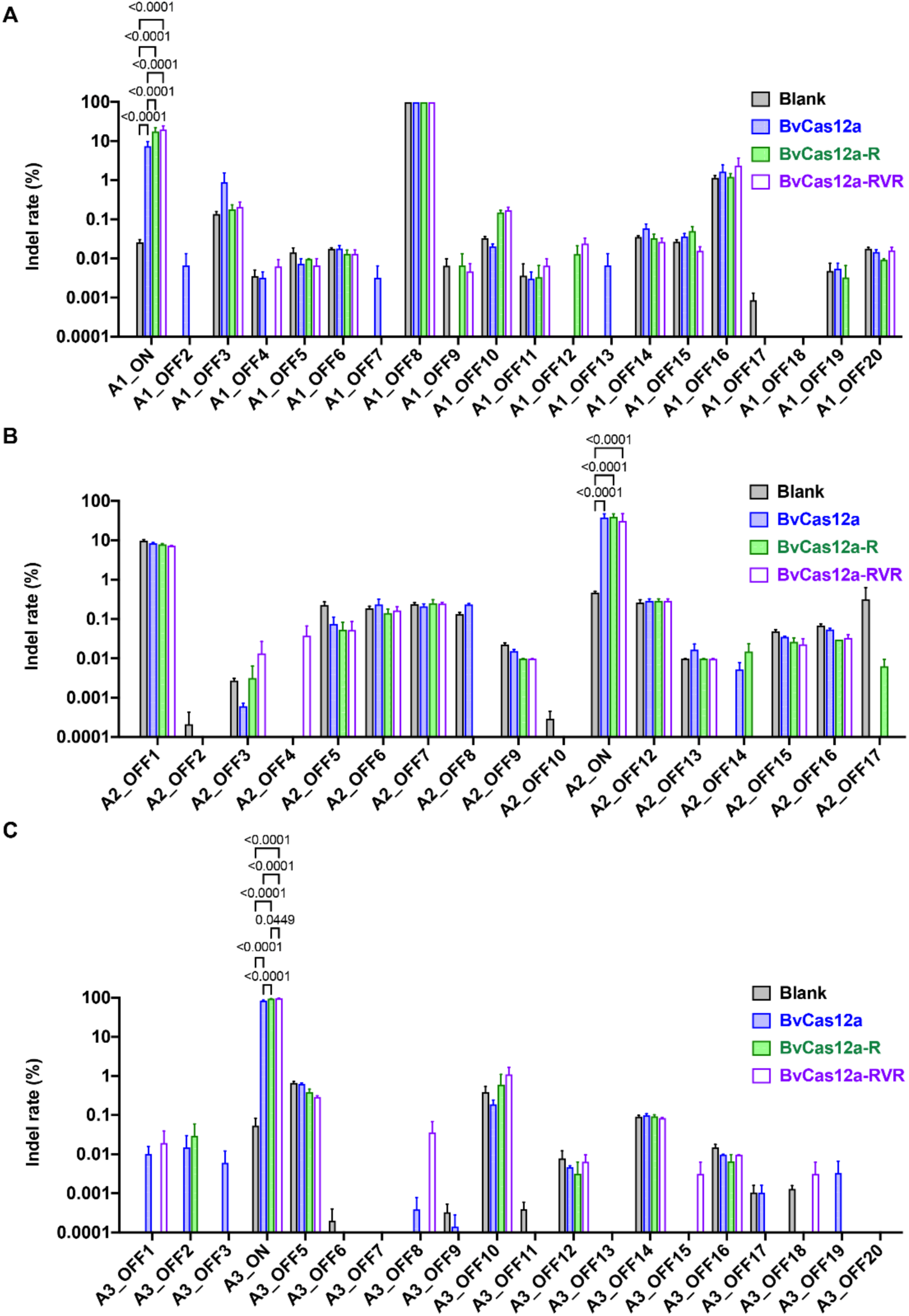
Comparison of Off-Target Mutation Induced by BvCas12a and Its Variants. (A, B, and C) The on-target (ON) and off-target (OFF) indel rates (%) for BvCas12a and its variants at three AAVS1 target sites, namely A1 site (A), A2 site (B), and A3 site (C), are depicted using logarithmic scale values. The off-target sites predicted for the three AAVS1 target sequences are consistent with previous data, as presented in Figure S4. Indel frequencies at the predicted off-target sites were assessed via targeted amplicon sequencing. The graph shows the mean values with standard error of the mean (SEM) based on three biological replicates. Statistical significance was determined using a two-way ANOVA followed by Tukey's multiple comparison test.

## Discussion

CRISPR-Cas12a is one of the most widely used gene editing systems, but its scope of use is limited by its PAM recognition sequence, editing efficiency, and specificity. Therefore, it is crucial to identify new CRISPR-Cas12a systems or engineer Cas12a proteins to overcome its limitations and expand the range of applications. Using the intestinal bacteria metagenomic sequencing data, we identified a new CRISPR-Cas12a system, derived from the anaerobic gram-negative bacterium *Butyricimonas virosa*. The CRISPR-Cas12a system effector protein BvCas12a shares the closest similarity with LbCas12a, but its amino acid sequence identity with LbCas12a is only 45.7%. The CRISPR-BvCas12a system recognizes the 5’ PAM TYTN and exhibits genome editing activity in eukaryotic cells. The scientific community has viewed incomplete CRISPR-Cas systems as evolutionary remnants, potentially less adept at precise and efficient genome editing. This perspective was based on the assumption that the absence of key components, such as Cas1 or Cas2, essential elements of the CRISPR-Cas system’s adaptation module,^34^ might render these systems functionally deficient. Nonetheless, the observation that the native locus of BvCas12a lacks both Cas1 and Cas2 challenges this view. It suggests incomplete CRISPR-Cas systems can still possess functional activity and could represent untapped sources of novel genome editing tools. This revelation underscores the importance of re-evaluating incomplete systems as potentially valuable resources in the ongoing quest to expand the CRISPR toolkit. It also highlights the need for comprehensive functional assays to fully understand the capabilities and limitations of these unconventional CRISPR-Cas variants.

BvCas12a showed slightly lower editing activity, and comparable, if not superior, editing specificity than AsCas12a. To enhance the editing capabilities, we conducted protein engineering with the rational design based on the complex structures of AsCas12a, LbCas12a previously reported, and BvCas12a predicted by AlphaFold. For the LbCas12a-crRNA complex, the looped-out helical domain (LHD) contains many positively charged amino acids on the side facing the central channel and located at an equivalent position to the wedge domain of SaCas9.^35^ The wedge domain in SaCas9 interacts with the PAM duplex of dsDNA, and the LHD in LbCas12a was also proved vital for dsDNA binding.^35,36^ The structure is also conserved in AsCas12a, and the P599 of AsCas12a inside LHD is proved to form van der Waals interactions with the nucleobase and deoxyribose moieties of dA(-2).^24^ In BvCas12a, T606 sits at the same spatial structure position with the P599 site of AsCas12a (Figure S7). Therefore, we constructed the T606P mutation and found that this mutation can significantly enhance the editing activity of BvCas12a (data not shown).

Further, to overcome the PAM restrictions of BvCas12a, we conduct protein engineering according to the structures and engineered variants of AsCas12a and LbCas12a with altered PAM preferences. For AsCas12a, two variants AsCas12a-RR (S542R/K607R) and AsCas12a-RVR (S542R/K548V/N552R) recognize the TYCV and TATV PAMs beside the canonical TTTV PAM.^22^ Similarly, two variants of LbCas12a, LbCas12a-RR (G532R/K595R) and LbCas12a-RVR (G532R/K538V/Y542R) recognize altered PAM as AsCas12a variants.^22,33^ We compared the amino acid of BvCas12a with other Cas12a orthologs and introduced N549R, K555V, and C559R mutants to BvCas12a in correspondence to the mutation in RVR mutants of AsCas12a and LbCas12a (Figure S7A). Finally, combined with sites associated with activity and PAM, we obtained two variants BvCas12a-R (N549R, T606P) and BvCas12a-RVR (N549R, K555V, C559R, T606P). As expected, BvCas12a-R and BvCas12a-RVR recognize 5’YYN and 5’YN PAM *in vitro*, respectively. Moreover, the variants exhibited enhanced activity in eukaryotic cells across all tested 5’ Y-rich PAMs, with a notable increase for non-T-rich PAMs. To discuss the mechanism by which BvCas12a variants broaden the recognition of the protospacer adjacent motif (PAM), we predicted the structure of BvCas12a and its variants complex with AlphaFold3 (Figure S7B). The predicted complex results showed that the K555 in wild-type BvCas12a can form a hydrogen bond with the adenine at position -3 (dA) in the target strand. In contrast, the RVR variant of BvCas12a lacks this K555 residue, which devastates its ability to interact with the -3 (dA). Moreover, BvCas12a variants with C559R and N549R substitutions facilitate an alternative interaction with the adenine at position -2 (dA). Together, these results demonstrated that BvCas12a variants expanded their PAM preferences. In the future, more research is needed to explore their PAM preference *in vivo*, as previous reports proved that PAM recognition *in vivo* may be stricter than *in vitro*. Usually, the broader PAM compatibility and higher gene editing activity may imply a higher potential for off-target interactions. Therefore, it is necessary to perform additional research to evaluate the off-target effects of these variants and potentially incorporate additional modifications to improve their accuracy. To our excitement, the two variants BvCas12a-R and BvCas12a-RVR have enhanced editing activity and expanded PAM preference without sacrificing specificity. So, the engineered variants BvCas12a-R and BvCas12a-RVR increase the targeting range in genome editing and provide useful tools in genome editing in the future.

**Figure S7.**
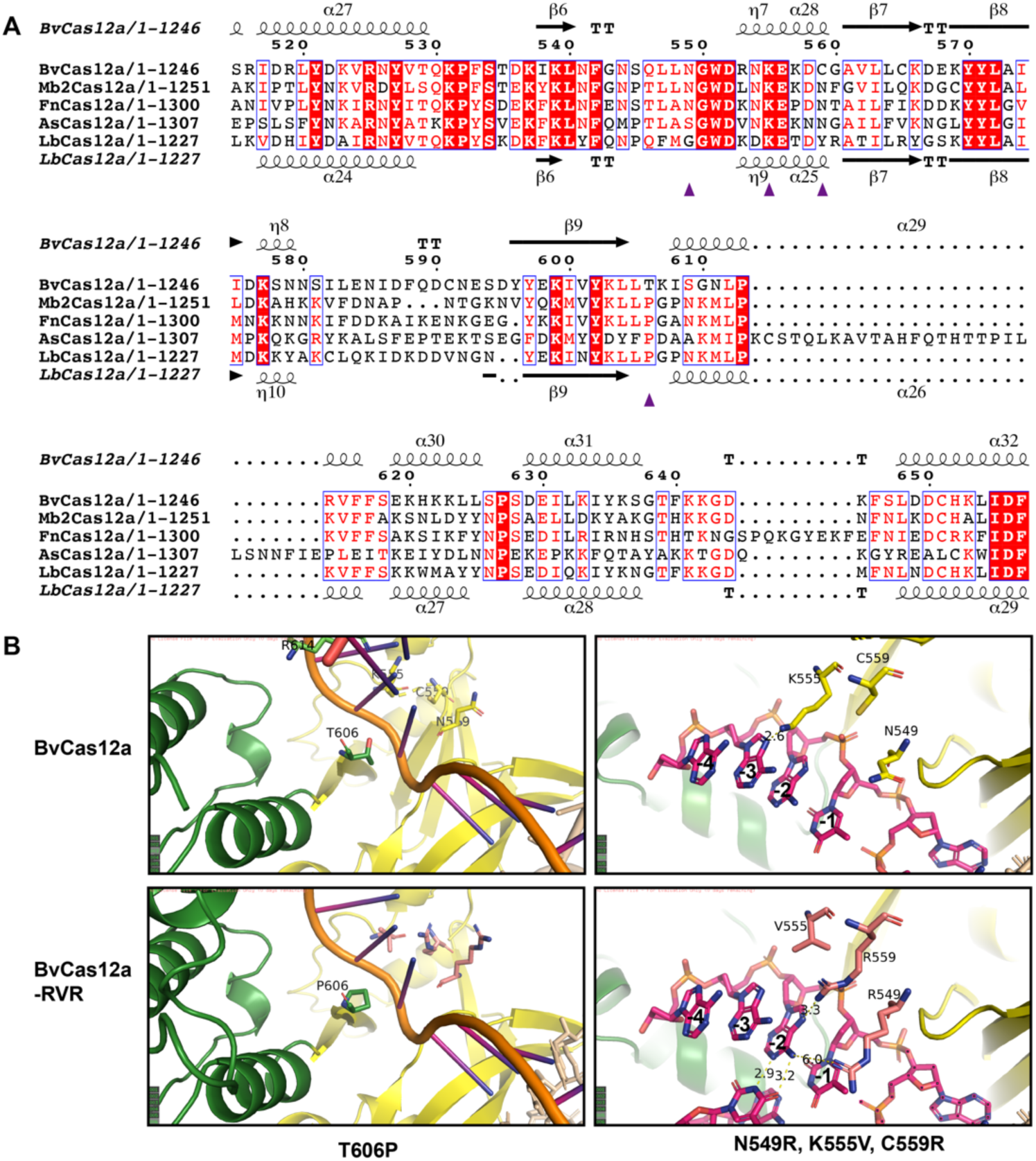
Sequence alignment and structure analysis of BvCas12a homologous proteins and BvCasI2a variants. (A) Protein sequence alignment of BvCasI 2a with its homologous proteins. The amino acids mutated in the BvCasI2a variant are marked with purple triangles. (B) Structural comparison highlighting differences between BvCas12a and its variant BvCas12a-RVR in PAM recognition. The structures of BvCasI 2a or BvCas12a-RVR in complex with target DNAwere predicted using AlphaFold3. The interactions between key amino acids and the crRNA-DNA duplex are shown as dashed lines.

Besides its application in genome editing, Cas12a has also been used in nucleoside detection with its collateral activity. The human papillomavirus (HPV) is a non-enveloped, double-stranded, circular DNA virus that causes various epithelial lesions and cancers. HPV is widespread, and early detection and accurate identification are crucial for preventing and treating cervical cancer, particularly for high-risk strains like HPV16 and HPV18.^37^ In this research, we confirmed that BvCas12a possesses collateral activity and detects HPV16 and HPV18 accurately within specific crRNA. The nucleic acid detection sensitivity of BvCas12a combined with recombinase polymerase amplification (RPA) achieved an attomolar level, that is comparable to that of LbCas12a. In the nucleic acid detection sensitivity assays, we observed a slight increase in fluorescence background when using mismatched crRNAs, which might be attributed to residual DNA templates following DNase treatment. This suggests that crRNA can activate the *trans*-cleavage activity of ssDNA by Cas12a even after *cis*-cleavage of the template ssDNA.^31^ Furthermore, BvCas12a was successfully employed for detecting HPV16-positive human genomic standard samples. All these results indicated that BvCas12a has the potential to be developed for molecular diagnostics with high sensitivity and specificity.

## MATERIALS AND METHODS

### Bioinformatical predictions of BvCas12a functions and features

BvCas12a was identified through computational analysis of bacterial genomes obtained from CNGB.^23^ CRISPR arrays were analyzed using PILER-CR (http://drive5.com/pilercr/) ^38^ with default parameters, and DNA sequences flanking 10 kb of the CRISPR arrays were extracted for further analysis. Open reading frames (ORFs) were predicted using MetaGeneMark (https://genemark.bme.gatech.edu/meta_gmhmmp.cgi)^39^ and then subjected to BLASTp to find Cas effector candidates. All structures presented were conducted using AlphaFold2 (multimer version) with the ColabFold notebook (https://github.com/sokrypton/ColabFold) or AlphaFold3 (https://alphafoldserver.com).^40^ The mature crRNAs were predicted based on the strong conservation of the direct repeat sequences for Cpf1-family proteins. The secondary structure of the sgRNA utilized in this study was predicted by RNAfold.^41^

### Plasmid construction

To obtain prokaryotic expression plasmids for protein purification, the coding sequences for BvCas12a, BvCas12a-R, and BvCas12a-RVR were synthesized by BGI Write and subsequently cloned into the pET28a plasmid featuring an N-terminal His tag.

For mammalian cell expression, the human codon-optimized sequences of BvCas12a, BvCas12a-R, and BvCas12a-RVR, each featuring dual nuclear localization signals (NLS) appended to both the N-and C-termini, were synthesized by BGI Write and cloned into a modified PX458 plasmid to create negative control (NC) plasmids. The NC plasmids were digested with BbsI (NEB) and purified via gel extraction (MACHEREY-NAGEL) to enable targeted insertion. Oligonucleotide strands with target sequences and their complementary strands were synthesized, denatured, and annealed to create double-stranded DNA (dsDNA). This dsDNA was inserted into the linearized NC plasmids utilizing T4 DNA ligase (NEB).

The NC plasmids sequences are listed in Supplementary Table 4, and all relevant targets, target primers, and sequences are listed in Supplementary Table 1 and Supplementary Table 3.

### Protein expression and purification

The prokaryotic expression plasmids coding for BvCas12a and its variants were transformed into the BL21(DE3) bacterial strain (Takara) and spread on LB agar plates supplemented with kanamycin. After transformation, positive colonies were selected and cultivated in a 50 mL LB medium containing 50 mg/L kanamycin. Then the cultured cells were transferred into 2 L LB medium with 50 mg/L kanamycin, and incubated at 37℃ in a shaking incubator until the OD600 reached 0.6. Subsequently, the cultures were cooled to 16℃, and protein expression was induced overnight by adding 0.5 mM isopropyl β-D-1-thiogalactopyranoside (IPTG) at the same temperature. The bacterial cells were harvested by centrifugation at 4℃, resuspended in lysis buffer A (20 mM Tris-HCl pH 7.5, 1 M NaCl, 20 mM imidazole, 1 mM PMSF), disrupted by sonication, and centrifuged to remove cellular debris. The clear supernatant was then loaded onto a Ni-NTA Sepharose 6 Fast Flow (Cytiva) column, and the column was agitated gently at 4℃ for one hour before eluting the proteins with buffer B (20 mM Tris-HCl pH 7.5, 1 M NaCl, 300 mM imidazole). Subsequent steps involved buffer exchange and desalting using an ultrafiltration tube to transfer the proteins into buffer C (20 mM Tris-HCl pH 7.5, 50 mM NaCl, 1 mM DTT). The AKTA chromatography system was then employed for further purification, with the sample being applied to a HiTrap Heparin HP column (Cytiva) followed by a linear gradient elution using both buffer C and buffer D (20 mM Tris-HCl, pH 7.5, 1 M NaCl, 1 mM DTT), progressively increasing the NaCl concentration. The protein was concentrated to 0.5 mL by ultrafiltration (MILLIPORE), after which the sample was subjected to size-exclusion chromatography on a Superdex 200 Increase 10/300 GL column (Cytiva). The proteins were eluted with buffer E (50 mM Tris-HCl pH 7.5, 500 mM NaCl, 1 mM DTT), and the principal peak fractions were pooled. The protein was analyzed by SDS-PAGE and concentrated to 100 μL by by ultrafiltration using a 50 kDa Amicon Ultra-0.5 ultrafiltration tube (Merck Millipore). Finally, the obtained proteins were mixed 1:1 with 70% glycerol, flash-frozen in liquid nitrogen, and stored at -80℃.

All relevant plasmid sequences are listed in Supplementary Table 4.

### *In vitro* cutting with purified protein

The crRNA was synthesized by transcribing a double-stranded DNA (dsDNA) template that contained the target-specific and scaffold sequences using the MEGAshortscript™ kit (Thermo Fisher Scientific) following the provided manual. It was then purified with the Guide-it IVT RNA Clean-Up Kit (Clontech). For *in vitro* cleavage assays, a reaction mixture containing 100 nM Cas protein, 200 nM crRNA, and 1×NEBuffer 2.1 was prepared. This mixture was first incubated at room temperature for 10 minutes. Subsequently, 200 ng of plasmids or PCR products were added, and the incubation continued at 37℃ for 30 minutes. The reaction was terminated with a 10× loading buffer, and the samples were analyzed by agarose gel electrophoresis.

All relevant DNA template sequences are listed in Supplementary Table 2.

### PAM identification

The ssDNA oligo library (Table 3) was synthesized by Gene Synthesis Platform (BGI) and used to prepare single-stranded circles (ssCir). For ssCir and DNA nanoballs (DNB) preparation, 50 fmol of ssDNA was utilized according to the instructions provided by the BGISEQ-500RS High-throughput Sequencing Kit (MGI Tech Co., Ltd.). PAM identification was performed based on the method described previously^25^ with a slight modification: 100 μL of DNBs and 33 μL of BGISEQ500 DNB loader Ⅱ (MGI Tech Co. Ltd.) were loaded onto the BGISEQ500 V3.1 chip. Single-end sequencing of 43 bases was performed following the instructions of the BGISEQ-500RS High-throughput Sequencing Set (SE100). For the DNB cleavage reaction, 0.3 μM Cas12a, 3 μM of the appropriate guide RNA, and 1 μL RNase inhibitor (Thermo Fisher Scientific) were combined in 1× NEBuffer™ r2.1 to a total volume of 300 μL, loaded onto the chip, and incubated at 37℃ for 3 hours. The PAM sequence logos were generated employing the R package ggseqlogo (https://github.com/omarwagih/ggseqlogo) ^42^.

### Cell culture and transfection

HEK293T cells were cultivated with DMEM medium (Gibco, 10566016) supplemented with 10% fetal bovine serum (Gibco, 10091148) and 1% non-essential amino acids (Gibco, 11140035). The cells were seeded onto a 12-well plate at a density of 200,000 cells per well one day before transfection. Plasmid transfection was performed with Lipofectamine 3000 (Thermo Fisher Scientific, L3000075) following the manufacturer’s protocol when cell confluency reached approximately 50%. The transfection efficiency was assessed by analyzing cell fluorescence 2 to 4 days after transfection.

### T7E1 assays

Cells were harvested after measuring the fluorescence rate, and genomic DNA was extracted using the TIANGEN Genomic DNA Kit. A specific fragment was then amplified by PCR with PrimeStar GXL DNA Polymerase (TAKARA), using 100 ng of genomic DNA as the template. The primers used for PCR amplification are listed in Supplementary Table 3. The PCR product was purified using the NucleoSpin Gel and PCR Clean-up Kit (MACHEREY-NAGEL), following the manufacturer’s protocol. Subsequently, 200 ng of the purified PCR product was denatured, reannealed, and digested with 0.3 μL of T7 Endonuclease I (NEB) at 37℃ for 20 minutes. After the reaction, 10× loading buffer was added, and the sample was subjected to agarose gel electrophoresis.

### Indel frequency analysis

Indel frequencies at on-target sites and predicted off-target sites were detected by targeted amplicon sequencing. The primers employed for PCR amplification to generate amplicons are listed in Supplementary Table 3. The amplicon-seq library was constructed using the MGIEasy PCR-Free DNA Library Preparation Kit according to the manufacturer’s protocol. For DNB preparation, 0.75 pmol of ssDNA was employed. Sequencing was carried out on a MGISEQ-2000RS sequencer (MGI). Genome editing efficiency for BvCas12a and AsCas12a was calculated by CRISPResso2^43^ or CRISPR-detector^44^ software. The data represent mean values with standard error of the mean (SEM) derived from three biological replicates. Statistical significance was evaluated using a two-way ANOVA with Tukey’s multiple comparison test.

### Collateral activity

For the collateral specificity analysis of BvCas12a, PCR amplification was conducted on standard samples containing HPV16 (BGI, RM-HPV014) and HPV18 (BGI, RM-HPV017), utilizing primers specific to each type. The substrates for subsequent analysis were prepared by purifying the PCR products. A reaction mixture of 250 nM Cas12a protein, 200 nM crRNA, NEBuffer™ r2.1 (NEB), and nuclease-free water was incubated at room temperature. Subsequently, 10 ng of the HPV16 or HPV18 PCR product was added to the mixture along with 10 μM ssDNA-FQ (fluorescein-quencher) probes (Gene Synthesis Platform, BGI). The reaction was conducted at 37℃, with fluorescence measurements taken every 5 minutes.

For collateral sensitivity assessment of BvCas12a and LbCas12a, isothermal amplification was performed on plasmids containing HPV sequences using specific primers, following the manufacturer’s instructions provided with the TwistAmp Basic reagent kit. After incubation at 37℃ for 10 min, the amplified products were used as dsDNA templates. Pre-incubated RNP complexes and ssDNA-FQ probes were then added to the system for subsequent detection, along with an additional inclusion of buffer 2.1 (NEB). In the final reaction system, the ratio of Cas12a: crRNA: FAM was 100 nM: 60 nM: 5 μM. The reaction proceeded at 37°C, with fluorescence measurements taken at five-minute intervals.

To confirm the application of BvCas12a in molecular diagnostics for human diseases, we detected HPV16-positive human reference genome samples (BGI, RM-HPV014) with BvCas12a-combined with isothermal amplification. The procedure was performed identically to the previous collateral sensitivity evaluation, with fluorescence measurements taken at five-minute intervals.

The primers used for PCR amplification and the ssDNA-FQ probes used in collateral activity assay are listed in Supplementary Table 3. All the primers used in this research were synthesized by Gene Synthesis Platform (BGI).

## Data availability

The data that support the conclusions of this study are accessible in the CNGB Sequence Archive (CNSA)^45^ of the China National GeneBank DataBase (CNGBdb),^46^ under accession number CNP0005302.

## Author contributions

H.L., Y.J. and Y.Z. conveived and designed the project. Y. Z. and L. X. provided the intestinal genome data resources. D.W., B.L., C.Q. and L.H. performed bioinformatics data mining, protein structure prediction and other bioinformatics analysis. X.S., C.L. and K.C. executed the experiments. X.S. and H.L. wrote the manuscript. The manuscript was revised by Y.Z. and approved by all members.

## Acknowledgments

We would like to thank our colleagues at China National GeneBank (Shenzhen).

