## Supplementary Tables for "Discovery and Engineering of a New BvCas12a Nuclease for Mammalian Genome Editing and Nucleic Acid Detection"

**Supplementary Table 1 DNA target**

| | | |
|---|---|---|
| **Target sites for validating *in vitro* activity of Cas proteins** | | |
| PAM library | TTCGGTAGCAGTTCCCTTTTGAG | Fig S1E, Fig S5A |
| Amp-23nt | gtatggcttcattcagctccggt | Fig 1A |
| In vitro-PAM/Temp | GGCCTGTGTAGGTGTTGAGGTAG | Fig 1B, Fig 4B, Fig S5B |
| HPV16-L1 | TGAAGTAGATATGGCAGCAC | Fig 4A, Fig 4C |
| HPV18-L1 | ACAATATGTGCTTCTACACA | |
| **Target sites for validating intracellular activity of Cas proteins** | | |
| AAVS1-gRNA1 | TCTGTCCCCTCCACCCCACAGTG | Fig 2B-2D, Fig S2A, Fig S3A-S3B, Fig 5B-5C |
| AAVS1-gRNA2 | GGCAGCTCCCCTACCCCCCTTAC | Fig 2A-2D Fig S2A-S2B, Fig S3A-S3B, Fig 5B-5C |
| AAVS1-gRNA3 | CTTACGATGGAGCCAGAGAGGATC | Fig 2B-2E, Fig S2A, Fig 3A-3C, Fig S3A-S3C, Fig 5B-5C |
| HBG-gRNA1 | CATTGAGATAGTGTGGGGAAGGG | Fig 5B-5C |
| HBG-gRNA2 | GTCAAGGCAAGGCTGGCCAACCC | |
| AAVS1-PAM-gRNA-TTCA | TTTGGGCAGCTCCCCTACCCCCC | Fig 5D-5E, Fig S5C |
| AAVS1-PAM-gRNA-CTTA | CCTCTCTAGTCTGTGCTAGCTCT | |
| AAVS1-PAM-gRNA-TCTC | TAGTCTGTGCTAGCTCTTCCAGC | |
| AAVS1-PAM-gRNA-CTCC | CCTACCCCCCTTACCTCTCTAGT | |
| AAVS1-PAM-gRNA-CCTC | TCTAGTCTGTGCTAGCTCTTCCA | |

**Supplementary Table 2 DNA templates for transcription *in vitro***

| | | |
|---|---|---|
| **T7 promoter** | | |
| T7-GGG | TAATACGACTCACTATAGgg | Fig 1A-1B, Fig S1E, Fig S5A-S5B, Fig 4A-4C |
| **Transcription template for crRNA targeting the random PAM library** | | |
| PAM library | CTCAAAAGGGAACTGCTACCGAAATCTACAATAGTAGAAATTccCTATAGTGAGTCGTATTA | Fig S1E, Fig S5A |
| **Transcription templates for crRNAs of different lengths targeting the Amp resistance gene** | | |
| Amp-20nt-U4AU4 | AAAATAAAAGGAGCTGAATGAAGCCATACATCTACAATAGTAGAAATTccCTATAGTGAGTCGTATTA | Fig 1A |
| Amp-24nt | AACCGGAGCTGAATGAAGCCATACATCTACAATAGTAGAAATTccCTATAGTGAGTCGTATTA | |
| Amp- | ACCGGAGCTGAATGAAGCCATACATCTACAATAGTAGAA | |

| | | |
|---|---|---|
| 23nt | ATTccCTATAGTGAGTCGTATTA | |
| Amp-22nt | CCGGAGCTGAATGAAGCCATACATCTACAATAGTAGAAATTccCTATAGTGAGTCGTATTA | |
| Amp-20nt | GGAGCTGAATGAAGCCATACATCTACAATAGTAGAAATTccCTATAGTGAGTCGTATTA | |
| Amp-18nt | AGCTGAATGAAGCCATACATCTACAATAGTAGAAATTccCTATAGTGAGTCGTATTA | |
| Amp-17nt | GCTGAATGAAGCCATACATCTACAATAGTAGAAATTccCTATAGTGAGTCGTATTA | |
| Amp-16nt | CTGAATGAAGCCATACATCTACAATAGTAGAAATTccCTATAGTGAGTCGTATTA | |
| Amp-15nt | TGAATGAAGCCATACATCTACAATAGTAGAAATTccCTATAGTGAGTCGTATTA | |
| Amp-14nt | GAATGAAGCCATACATCTACAATAGTAGAAATTcCTATAGTGAGTCGTATTA | |
| Amp-13nt | AATGAAGCCATACATCTACAATAGTAGAAATTcCTATAGTGAGTCGTATTA | |
| Amp-12nt | ATGAAGCCATACATCTACAATAGTAGAAATTcCTATAGTGAGTCGTATTA | |
| Amp-11nt | TGAAGCCATACATCTACAATAGTAGAAATTcCTATAGTGAGTCGTATTA | |
| Amp-10nt | GAAGCCATACATCTACAATAGTAGAAATTcCTATAGTGAGTCGTATTA | |
| **Transcription template for crRNAs targeting a fixed sequence but with different PAMs or at different temperatures** | | |
| Temp-crRNA-R | CTACCTCAACACCTACACAGGCCATCTACAATAGTAGAAATTccCTATAGTGAGTCGTATTA | Fig 1B, Fig S5B |
| **Transcription templates for crRNAs of validating *trans*-cleavage activity** | | |
| HPV16-L1-gRNA | GTGCTGCCATATCTACTTCAATCTACAATAGTAGAAATTccCTATAGTGAGTCGTATTA | Fig 4A, Fig 4C |
| HPV18-L1-gRNA | TGTGTAGAAGCACATATTGTATCTACAATAGTAGAAATTccCTATAGTGAGTCGTATTA | |
| BvCas12a-RPA-gRNA | CTACCTCAACACCTACACAGGATCTACAATAGTAGAAATTccCTATAGTGAGTCGTATTA | Fig 4B |
| LbCas12a-RPA- | CTACCTCAACACCTACACAGGCCATCTACACTTAGTAGAAATTccCTATAGTGAGTCGTATTA | |

| |
|---|
| gRNA |

**Supplementary Table 3 Primers and oligonucleotides**

| | | |
|---|---|---|
| **Oligonucleotide for DocMF** | | |
| ssDNA oligo | 5'-Phosphorylation GAACGACATGGCTACGATCCGACTTNNNNNNNNNNTTCGGTAGCAGTTCCCTTTTGAGNNNNNNNNNNNAAGTCGGAGGCCAAGCGGTCTTAGGAAGACAATTGCCTAGGCCAACTCCTTGGCTCACA | Fig 5A |
| **Primers for T7E1 activity validation at the *AAVS1* site** | | |
| AAVS1-PCR-F | TTTCCGGAGCACTTCCTTCT | Fig S2A-S2B, Fig S5C |
| AAVS1-PCR-R | CAGGAACCCCTGTAGGGAAG | |
| **Insert fragments for constructing *in vivo* cleavage plasmids** | | |
| AAVS1-1-F: | AGATTCTGTCCCCTCCACCCCACAGTG | Fig 2B-2D, Fig S2A, Fig S3A-S3B, Fig 5B-5C |
| AAVS1-2-F: | AGATGGCAGCTCCCCTACCCCCCTTAC | |
| AAVS1-3-F: | AGATCTTACGATGGAGCCAGAGAGGAT | |
| AAVS1-1-R: | aaaaCACTGTGGGGTGGAGGGGACAGA | |
| AAVS1-2-R: | aaaaGTAAGGGGGGTAGGGGAGCTGCC | |
| AAVS1-3-R: | aaaaATCCTCTCTGGCTCCATCGTAAG | |
| HDV-AAVS1-1-R: | ggccCACTGTGGGGTGGAGGGGACAGA | Fig 5B-5C |
| HDV-AAVS1-2-R: | ggccGTAAGGGGGGTAGGGGAGCTGCC | Fig 2A, Fig S2B, Fig 5B-5C |
| HDV-AAVS1-3-R: | ggccATCCTCTCTGGCTCCATCGTAAG | Fig 5B-5C |
| AAVS1-2-24nt-F | AgatGGCAGCTCCCCTACCCCCCTTACC | Fig 2A, Fig S2B |
| AAVS1-2-23nt-F | agatGGCAGCTCCCCTACCCCCCTTAC | |
| AAVS1-2-22nt-F | agatGGCAGCTCCCCTACCCCCCTTA | |
| AAVS1-2-20nt-F | agatGGCAGCTCCCCTACCCCCCT | |
| AAVS1-2-18nt-F | agatGGCAGCTCCCCTACCCCC | |
| AAVS1-2-16nt-F | agatGGCAGCTCCCCTACCC | |
| HDV-AAVS1-2-24nt-R | ggccGGTAAGGGGGGTAGGGGAGCTGCC | |
| HDV-AAVS1-2-23nt-R | ggccGGTAAGGGGGGTAGGGGAGCTGC | |
| HDV-AAVS1-2- | ggccGGTAAGGGGGGTAGGGGAGCTG | |

| | | |
|---|---|---|
| 22nt-R | | |
| HDV-AAVS1-2-20nt-R | ggccGGTAAGGGGGGTAGGGGAGC | |
| HDV-AAVS1-2-18nt-R | ggccGGTAAGGGGGGTAGGGGA | |
| HDV-AAVS1-2-16nt-R: | ggccGGTAAGGGGGGTAGGG | |
| HBG-TTTN-F | agatCCTTGTCAAGGCTATTGGTCAAG | Fig 5B-5C |
| HBG-TTTN-R | ggccCTTGACCAATAGCCTTGACAAGG | |
| HBG-TTN-F | agatGTCAAGTTTGCCTTGTCAAGGCT | |
| HBG-TTN-R | ggccAGCCTTGACAAGGCAAACTTGAC | |
| AAVS1-TTCA-F | agatTTTGGGCAGCTCCCCTACCCCCC | Fig 5D-5E, Fig S5C |
| AAVS1-CTTA-F | agatCCTCTCTAGTCTGTGCTAGCTCT | |
| AAVS1-TCTC-F | agatTAGTCTGTGCTAGCTCTTCCAGC | |
| AAVS1-CTCC-F | agatCCTACCCCCCTTACCTCTCTAGT | |
| AAVS1-CCTC-F | agatTCTAGTCTGTGCTAGCTCTTCCA | |
| AAVS1-TTCA-R | ggccGGGGGGTAGGGGAGCTGCCCAAA | |
| AAVS1-CTTA-R | ggccAGAGCTAGCACAGACTAGAGAGG | |
| AAVS1-TCTC-R | ggccGCTGGAAGAGCTAGCACAGACTA | |
| AAVS1-CTCC-R | ggccACTAGAGAGGTAAGGGGGGTAGG | |
| AAVS1-CCTC-R | ggccTGGAAGAGCTAGCACAGACTAGA | |
| **Primers for amplifying plasmids to obtain substrates containing target sequences with different PAMs** | | |
| PAM library-F | GGTGTCGGGGCTGG | Fig S1E, Fig S5A |
| PAM library-R | GTGATGCTCGTCAGGGG | |
| PAM-plasmid-F | GGACAGGTATCCGGTAAGCG | Fig 1B, Fig S5B |
| PAM-plasmid-R | AGAGTGAAGCAGAACGTGGG | |
| **Insert fragments for constructing substrates containing target sequences with different PAMs** | | |
| PAM-plasmid-CCTA-F: | agatCCTAGGCCTGTGTAGGTGTTGAGGTAG | Fig S5B |
| PAM- plasmid-CCTA-R: | ggccCTACCTCAACACCTACACAGGCCTAGG | |
| PAM- plasmid-TTCA-F: | agatTTCAGGCCTGTGTAGGTGTTGAGGTAG | |

| | | |
|---|---|---|
| PAM- plasmid-TTCA-R: | ggccCTACCTCAACACCTACACAGGCCTGAA | |
| PAM- plasmid-TCTA-F: | agatTCTAGGCCTGTGTAGGTGTTGAGGTAG | |
| PAM- plasmid-TCTA-R: | ggccCTACCTCAACACCTACACAGGCCTAGA | |
| PAM- plasmid-CTTA-F: | agatCTTAGGCCTGTGTAGGTGTTGAGGTAG | |
| PAM- plasmid-CTTA-R: | ggccCTACCTCAACACCTACACAGGCCTAAG | |
| PAM- plasmid-CTCA-F: | agatCTCAGGCCTGTGTAGGTGTTGAGGTAG | |
| PAM- plasmid-CTCA-R: | ggccCTACCTCAACACCTACACAGGCCTGAG | |
| PAM/Temp-plasmid-TTTA-F: | agatTTTAGGCCTGTGTAGGTGTTGAGGTAG | Fig1B, Fig S5B |
| PAM/Temp -plasmid-TTTA-R: | ggccCTACCTCAACACCTACACAGGCCTAAA | |
| PAM- plasmid-AATA-F: | agatAATAGGCCTGTGTAGGTGTTGAGGTAG | Fig S5B |
| PAM- plasmid-AATA-R: | ggccCTACCTCAACACCTACACAGGCCTATT | |
| PAM- plasmid-AACA-F: | agatAACAGGCCTGTGTAGGTGTTGAGGTAG | |
| PAM- plasmid-AACA-R: | ggccCTACCTCAACACCTACACAGGCCTGTT | |
| PAM- plasmid-GTTA-F: | agatGTTAGGCCTGTGTAGGTGTTGAGGTAG | |
| PAM- plasmid-GTTA-R: | ggccCTACCTCAACACCTACACAGGCCTAAC | |
| PAM- plasmid-ACAA-F: | agatACAAGGCCTGTGTAGGTGTTGAGGTAG | |
| PAM- plasmid-ACAA-R: | ggccCTACCTCAACACCTACACAGGCCTTGT | |
| PAM- plasmid-AAGA-F: | agatAAGAGGCCTGTGTAGGTGTTGAGGTAG | |
| PAM- plasmid-AAGA-R: | ggccCTACCTCAACACCTACACAGGCCTCTT | |
| **Primers and oligonucleotides for validating *trans*-cleavage activity** | | |
| FAM | 5`6-FAM-AAAAAA- 3`BHQ1 | Fig 4A-4C |
| HPV16-L1-F | TTGTTGGGGTAACCAACTATTTGTTACTGTT | Fig 4A, Fig 4C-4D |

| | | |
|---|---|---|
| HPV16-L1-R | CCTCCCCATGTCGTAGGTACTCCTTAAAG | |
| HPV18-L1-F | GCATAATCAATTATTTGTTACTGTGGTAGATACCACT | |
| HPV18-L1-R | GCTATACTGCTTAAATTTGGTAGCATCATATTGC | |
| RPA-plasmid-F | cttggctttatatatcttgtggaaaggacg | Fig 4B |
| RPA-plasmid-R | cggactagccttattttaacttgctatttcta | |
| **Primers for amplicon library construction and sequencing at *AAVS1* site and *HBG* site** | | |
| AAVS1-A1-ON-F | ACTTCAGGACAGCATGTTTGC | Fig S4A-S4B, Fig S6A |
| AAVS1-A1-ON-R | GGGGGTGTGTCACCAGATAA | |
| AAVS1-A2-ON-F | GCCTGCATCATCACCGTTTTT | |
| AAVS1-A2-ON-R | CTGTCCTGAAGTGGACATAGGG | |
| AAVS1-A3-ON-F | AGGGACAGGATTGGTGACAG | |
| AAVS1-A3-ON-R | CAGGGTGGCCACTGAGAAC | |
| HBG-ON-F | ACTTCAGGACAGCATGTTTGC | |
| HBG-ON-R | GGGGGTGTGTCACCAGATAA | |
| AAVS1-A1-OFF-F1 | AGGTTCTGGCAAGGAGAGAGATG | |
| AAVS1-A1-OFF-F2 | TCTGGGAGTACTAGAAATGCAGT | |
| AAVS1-A1-OFF-F3 | CTCCTGTACTTGCCCCATCCTTG | |
| AAVS1-A1-OFF-F4 | TTGACACCCAAACCTCCAAATGC | |
| AAVS1-A1-OFF-F5 | TTGTACCTCTCACTGTTGCTGGT | |
| AAVS1-A1-OFF-F6 | Acagctttgattggacaagagactg | |
| AAVS1-A1-OFF-F7 | CAGAGGAAGTCAGCAGCCCTAG | |
| AAVS1-A1-OFF-F8 | ctgcagtgagtgagccatgattg | |
| AAVS1-A1-OFF-F9 | aaacactTCGAGGGAGGGGAAAA | |
| AAVS1-A1-OFF-F10 | TGTGGAGTTGATTTGCCTACATCT | |

| | |
|---|---|
| AAVS1-A1-OFF-F11 | tggCTCCACAATTCATATCCCACA |
| AAVS1-A1-OFF-F12 | ATCTTTCCTCCTCTCCATTGCCC |
| AAVS1-A1-OFF-F13 | GGAGTGTGTTTTCCTAGGTAGTGGT |
| AAVS1-A1-OFF-F14 | GAGATGATGGACGTGGCTGATGA |
| AAVS1-A1-OFF-F15 | CCAAAATGTCTCTCTGGTGCCAC |
| AAVS1-A1-OFF-F16 | ACAGCAAACAAATATTAAATGGGGCA |
| AAVS1-A1-OFF-F17 | AGGTCTCTCTCTATGAATCTGTGACT |
| AAVS1-A1-OFF-F18 | TCCCTGGTTGAAGCATTAATTCT |
| AAVS1-A1-OFF-F19 | CAGTGGGAATTTGGGAGTGAAGC |
| AAVS1-A1-OFF-F20 | aggaggagaggaagaaaagagaaag |
| AAVS1-A1-OFF-R1 | ATATTCCCAGGGCCGGTTAATGT |
| AAVS1-A1-OFF-R2 | tctcTGCCTCAATGTTGTCCCTT |
| AAVS1-A1-OFF-R3 | AGATCCATACCATGCATTCTTGAA |
| AAVS1-A1-OFF-R4 | ccaggctggcgtatagtggtg |
| AAVS1-A1-OFF-R5 | ACTTGCTCTGACTTTGTGTTGGC |
| AAVS1-A1-OFF-R6 | TGCATGCTGAGTAAACAAGCCTG |
| AAVS1-A1-OFF-R7 | CCAACAGTAGGGCAGTGCATTTT |
| AAVS1-A1-OFF-R8 | gggatcatcaaactgctttccaca |
| AAVS1-A1-OFF-R9 | AGAAGACCACCAACCACAGAGTG |
| AAVS1-A1-OFF-R10 | CCTCTGAAAATAAGCAcggtggaaa |
| AAVS1-A1-OFF-R11 | CTCCCCAGGCCTGAGTCCATAG |
| AAVS1-A1-OFF-R12 | GGCAAGGTGGTTAGGaagagctt |

| | | |
|---|---|---|
| AAVS1-A1-OFF-R13 | ACAGGATATGCTTGCTTCGagaa | |
| AAVS1-A1-OFF-R14 | GATACAGGGAACGAAGAGGACCT | |
| AAVS1-A1-OFF-R15 | GGCAACAGAACAACTAAACCTAGGG | |
| AAVS1-A1-OFF-R16 | aattgctgagctTGGTGGTGTG | |
| AAVS1-A1-OFF-R17 | tctgtttatcctTGTGCCAGACA | |
| AAVS1-A1-OFF-R18 | ccaccctgcctgaccgtaaaa | |
| AAVS1-A1-OFF-R19 | CCTTCCAAAGCACAACCACCAAG | |
| AAVS1-A1-OFF-R20 | CTGACTCTCCCTGACCCACATTC | |
| AAVS1-A2-OFF-F1 | CAGACTTGCAGGAGGAGATGAGG | Fig S4C-S4D, Fig S6B |
| AAVS1-A2-OFF-F2 | TGCTATTAGTCATCACATCCCTATTT | |
| AAVS1-A2-OFF-F3 | GCTGTCAGATGAAGGCTTGCTTC | |
| AAVS1-A2-OFF-F4 | GGGGCCTGATGAGATGATAATCA | |
| AAVS1-A2-OFF-F5 | GCGTTCCAGACAAAAGCTCGTTT | |
| AAVS1-A2-OFF-F6 | AGGGTCactaaaagagaatgaaatca | |
| AAVS1-A2-OFF-F7 | agagttctGCCGACTTTCCTCTT | |
| AAVS1-A2-OFF-F8 | CGTCCTCAGTGTACAGAGTTGCT | |
| AAVS1-A2-OFF-F9 | gctgaactTGGCAACCTAATTCCT | |
| AAVS1-A2-OFF-F10 | tgcagtgaACAGGGGAGTACTAC | |
| AAVS1-A2-OFF-F11 | GCCCTGGGAATATAAGGTGGTCC | |
| AAVS1-A2-OFF-F12 | TTAGTAATGCCCTCCTTTGCCCA | |
| AAVS1-A2-OFF-F13 | CTCCCCTTCCAGCCCATTTAGTT | |
| AAVS1-A2-OFF-F14 | CTCACTTTGGCTCTCAGGTTCCT | |

| | | |
|---|---|---|
| AAVS1-A2-OFF-F15 | CAGAACACCAAGGCAGGGAGT | |
| AAVS1-A2-OFF-F16 | cctcagtggttctcagccttgg | |
| AAVS1-A2-OFF-F17 | AGTGTAGAACATGGGGAAGGAGC | |
| AAVS1-A2-OFF-R1 | CGTAGAAAAGGTTAATGATTGCGTGT | |
| AAVS1-A2-OFF-R2 | aagggaattgcttgaatccagg | |
| AAVS1-A2-OFF-R3 | CCGACAATTATCCACAGCCAAGG | |
| AAVS1-A2-OFF-R4 | ccccattctgtaggatgtcaCCC | |
| AAVS1-A2-OFF-R5 | ACCCTCTCCTTCCTAACCTCCAT | |
| AAVS1-A2-OFF-R6 | TTGAGCTTCTTCCTtccatctaac | |
| AAVS1-A2-OFF-R7 | TGCATGCAGAGAGGACAGAAAGA | |
| AAVS1-A2-OFF-R8 | AAATGAAAGTCACatcctggggt | |
| AAVS1-A2-OFF-R9 | AGATGATCTGGAAAAGTCTACACAAA | |
| AAVS1-A2-OFF-R10 | AGGATTAGTCTGGTGAAAGGGCA | |
| AAVS1-A2-OFF-R11 | CCTCTTGCTTTCTTTGCCTGGAC | |
| AAVS1-A2-OFF-R12 | TCAGGCATTGCTACTCTGGTCTC | |
| AAVS1-A2-OFF-R13 | ATTTCCTGGCTTGGCTGTGAATG | |
| AAVS1-A2-OFF-R14 | GCCAACCTCGAAATTCATGAAGGG | |
| AAVS1-A2-OFF-R15 | CCATGGGTCGAGTCCTCATTCTA | |
| AAVS1-A2-OFF-R16 | CCTGCATTTTAAAGAGCTCCTGGG | |
| AAVS1-A2-OFF-R17 | TAGGGAATGCCTGGAACTGTCG | |
| AAVS1-A3-OFF-F1 | tcttgtttcaatttGGGCTGGTGA | Fig S4E-S4F, Fig S6C |
| AAVS1-A3-OFF-F2 | CCACTTCCATCCCCAGTTCCTTG | |

| | |
|---|---|
| AAVS1-A3-OFF-F3 | AGGCGGCTCAAGATCCAAGTATT |
| AAVS1-A3-OFF-F4 | CAGGTTCTGGGAGAGGGTAGC |
| AAVS1-A3-OFF-F5 | TGAGCCTTTGGATAAGTTAacccc |
| AAVS1-A3-OFF-F6 | AGCATGCAACTTCTCCTTTTCTct |
| AAVS1-A3-OFF-F7 | GTGTTCAGTCATGTGGCCATTGT |
| AAVS1-A3-OFF-F8 | TCCAGTAGCCTTCAAGCCACAAG |
| AAVS1-A3-OFF-F9 | tCTATCTCTCTGTAATGGCCCCA |
| AAVS1-A3-OFF-F10 | GATCTGTCcttgagcctctgg |
| AAVS1-A3-OFF-F11 | acctggccaagtgaGTTAATTATGC |
| AAVS1-A3-OFF-F12 | CCATTCAGGGCAATAATCACAGA |
| AAVS1-A3-OFF-F13 | GCAGTGTAAGCCAAGAGATTGCA |
| AAVS1-A3-OFF-F14 | AAACCTTCAACTGCAGCACACAT |
| AAVS1-A3-OFF-F15 | GAGGGATGGGTCAGGTTCAGg |
| AAVS1-A3-OFF-F16 | CATCTGTCGTTGCCTCACCAG |
| AAVS1-A3-OFF-F17 | TGAGTTTGTTCTTTGATGCAAGAGT |
| AAVS1-A3-OFF-F18 | TAGTGTGGGGAGGACATTGCTAG |
| AAVS1-A3-OFF-F19 | GAGGTTAGTTACATTCCAGCTCAA |
| AAVS1-A3-OFF-F20 | CCTGGTAGGTAGGCATGGATTAGG |
| AAVS1-A3-OFF-F21 | tggaagagggTTACATAATGACAAGT |
| AAVS1-A3-OFF-R1 | tggagaaagagaagggagatgtGC |
| AAVS1-A3-OFF-R2 | GGGAACACTGGCCTTTCTGCA |
| AAVS1-A3-OFF-R3 | TTCCACAAAGTCCTACCAGCCTT |

| | |
|---|---|
| AAVS1-A3-OFF-R4 | CCTCCTCCTTCCTAGTCTCCTGA |
| AAVS1-A3-OFF-R5 | GCATCTGTCATGTTGTAGGCAGT |
| AAVS1-A3-OFF-R6 | TCTCAGTTTGATCTCCCTTTGGCA |
| AAVS1-A3-OFF-R7 | TGGCACTGTACTTTTGGGAAACAC |
| AAVS1-A3-OFF-R8 | AACCTTTGCCCTCGTTATTGGTT |
| AAVS1-A3-OFF-R9 | GATGAGGAGGATGATGAGGAGGC |
| AAVS1-A3-OFF-R10 | gactccCACATGCTAAACTCCCT |
| AAVS1-A3-OFF-R11 | cattggttacCTGGTGTACGTGC |
| AAVS1-A3-OFF-R12 | TTGGAGCCACAAAGAAATGCAGA |
| AAVS1-A3-OFF-R13 | CAAGTCACACATCCCTGGAGTCA |
| AAVS1-A3-OFF-R14 | ATTTTAGGGTTTGGGGCCACAAC |
| AAVS1-A3-OFF-R15 | CTGTGGCTGCCTGTTGTCTAAGT |
| AAVS1-A3-OFF-R16 | TTTAAGGGATGGGCAGCGTTTTC |
| AAVS1-A3-OFF-R17 | acacattctcttCAAGTACACATAGA |
| AAVS1-A3-OFF-R18 | GCCTGCCTACCTCTTAGCCC |
| AAVS1-A3-OFF-R19 | GCAAGGGAGCTAAGGGGAAG |
| AAVS1-A3-OFF-R20 | TGCCAGAGAAACCATAAGCCAGA |
| AAVS1-A3-OFF-R21 | TTTACCTTCTTTGGAGTGGGGCA |

**Supplementary Table 4 Plasmids**

**Prokaryotic and eukaryotic expression plasmid sequences containing Cas proteins**

| **Plasmid of the random PAM library** | | |
|---|---|---|
| Plsmid-PAM-library | TCGCGCGTTTCGGTGATGACGGTGAAAACCTCTGACACATGCAGCTCCCGGAGACGGTCACAGCTTGTCTGTAAGCGGATGCCGGGAGCAGACAAGCCCGTCAGGGCGCGTCAGCGGGTGTTGGCG | Fig S1E, Fig S5A |

| | |
|---|---|
| | GGTGTCGGGGCTGGCTTAACTATGCGGCATCAGAGCAGATTGT<br>ACTGAGAGTGCACCATATGCGGTGTGAAATACCGCACAGATG<br>CGTAAGGAGAAAATACCGCATCAGGCGCCATTCGCCATTCAG<br>GCTGCGCAACTGTTGGGAAGGGCGATCGGTGCGGGCCTCTTC<br>GCTATTACGCCAGCTGGCGAAAGGGGGATGTGCTGCAAGGCG<br>ATTAAGTTGGGTAACGCCAGGGTTTTCCCAGTCACGACGTTGT<br>AAAACGACGGCCAGTGAATTAGAACTCGGTACGCGCGGATCT<br>TCCAGAGATtNNNNNNNNTTCGGTAGCAGTTCCCTTTTGAGCTA<br>CTACaATCGTCGAACGGCAGGCGTGCAAACTTGGCGTAATCAT<br>GGTCATAGCTGTTTCCTGTGTGAAATTGTTATCCGCTCACAATT<br>CCACACAACATACGAGCCGGAAGCATAAAGTGTAAAGCCTGG<br>GGTGCCTAATGAGTGAGCTAACTCACATTAATTGCGTTGCGCT<br>CACTGCCCGCTTTCCAGTCGGGAAACCTGTCGTGCCAGCTGCA<br>TTAATGAATCGGCCAACGCGCGGGGAGAGGCGGTTTGCGTAT<br>TGGGCGCTCTTCCGCTTCCTCGCTCACTGACTCGCTGCGCTCG<br>GTCGTTCGGCTGCGGCGAGCGGTATCAGCTCACTCAAAGGCG<br>GTAATACGGTTATCCACAGAATCAGGGGATAACGCAGGAAAG<br>AACATGTGAGCAAAAGGCCAGCAAAAGGCCAGGAACCGTAA<br>AAAGGCCGCGTTGCTGGCGTTTTTCCATAGGCTCCGCCCCCCT<br>GACGAGCATCACAAAAATCGACGCTCAAGTCAGAGGTGGCGA<br>AACCCGACAGGACTATAAAGATACCAGGCGTTTCCCCCTGGA<br>AGCTCCCTCGTGCGCTCTCCTGTTCCGACCCTGCCGCTTACCG<br>GATACCTGTCCGCCTTTCTCCCTTCGGGAAGCGTGGCGCTTTCT<br>CATAGCTCACGCTGTAGGTATCTCAGTTCGGTGTAGGTCGTTC<br>GCTCCAAGCTGGGCTGTGTGCACGAACCCCCCGTTCAGCCCG<br>ACCGCTGCGCCTTATCCGGTAACTATCGTCTTGAGTCCAACCC<br>GGTAAGACACGACTTATCGCCACTGGCAGCAGCCACTGGTAA<br>CAGGATTAGCAGAGCGAGGTATGTAGGCGGTGCTACAGAGTT<br>CTTGAAGTGGTGGCCTAACTACGGCTACACTAGAAGAACAGTA<br>TTTGGTATCTGCGCTCTGCTGAAGCCAGTTACCTTCGGAAAAA<br>GAGTTGGTAGCTCTTGATCCGGCAAACAAACCACCGCTGGTAG<br>CGGTGGTTTTTTTGTTTGCAAGCAGCAGATTACGCGCAGAAAA<br>AAAGGATCTCAAGAAGATCCTTTGATCTTTTCTACGGGGTCTGA<br>CGCTCAGTGGAACGAAAACTCACGTTAAGGGATTTTGGTCATG<br>AGATTATCAAAAAGGATCTTCACCTAGATCCTTTTAAATTAAAA<br>ATGAAGTTTTAAATCAATCTAAAGTATATATGAGTAAACTTGGT<br>CTGACAGTTACCAATGCTTAATCAGTGAGGCACCTATCTCAGC<br>GATCTGTCTATTTCGTTCATCCATAGTTGCCTGACTCCCCGTCG<br>TGTAGATAACTACGATACGGGAGGGCTTACCATCTGGCCCCAG<br>TGCTGCAATGATACCGCGAGACCCACGCTCACCGGCTCCAGA<br>TTTATCAGCAATAAACCAGCCAGCCGGAAGGGCCGAGCGCAG<br>AAGTGGTCCTGCAACTTTATCCGCCTCCATCCAGTCTATTAATT<br>GTTGCCGGGAAGCTAGAGTAAGTAGTTCGCCAGTTAATAGTTT<br>GCGCAACGTTGTTGCCATTGCTACAGGCATCGTGGTGTCACGC |

| | | |
|---|---|---|
| | TCGTCGTTTGGTATGGCTTCATTCAGCTCCGGTTCCCAACGATC<br>AAGGCGAGTTACATGATCCCCCATGTTGTGCAAAAAAGCGGTT<br>AGCTCCTTCGGTCCTCCGATCGTTGTCAGAAGTAAGTTGGCCG<br>CAGTGTTATCACTCATGGTTATGGCAGCACTGCATAATTCTCTT<br>ACTGTCATGCCATCCGTAAGATGCTTTTCTGTGACTGGTGAGTA<br>CTCAACCAAGTCATTCTGAGAATAGTGTATGCGGCGACCGAGT<br>TGCTCTTGCCCGGCGTCAATACGGGATAATACCGCGCCACATA<br>GCAGAACTTTAAAAGTGCTCATCATTGGAAAACGTTCTTCGGG<br>GCGAAAACTCTCAAGGATCTTACCGCTGTTGAGATCCAGTTCG<br>ATGTAACCCACTCGTGCACCCAACTGATCTTCAGCATCTTTTAC<br>TTTCACCAGCGTTTCTGGGTGAGCAAAAACAGGAAGGCAAAAT<br>GCCGCAAAAAAGGGAATAAGGGCGACACGGAAATGTTGAATA<br>CTCATACTCTTCCTTTTTCAATATTATTGAAGCATTTATCAGGGT<br>TATTGTCTCATGAGCGGATACATATTTGAATGTATTTAGAAAAA<br>TAAACAAATGGGGGTTCCGCGCACATTTCCCCGAAAAGTGCCA<br>CCTGACGTCTAAGAAACCATTATTATCATGACATTAACCTATAA<br>AAATAGGCGTATCACGAGGCCCTTTCGTC | |
| **Plasmid substrate with Amp resistance** | | |
| Plasmid-Amp | tcgcgcgtttcggtgatgacggtgaaaacctctgacacatgcagctcccggagacg<br>gtcacagcttgtctgtaagcggatgccgggagcagacaagcccgtcagggcgcgt<br>cagcgggtgttggcgggtgtcggggctggcttaactatgcggcatcagagcagat<br>tgtactgagagtgcaccatatgcggtgtgaaataccgcacagatgcgtaaggaga<br>aaataccgcatcaggcgccattcgccattcaggctgcgcaactgttgggaagggc<br>gatcggtgcgggcctcttcgctattacgccagctggcgaaagggggatgtgctgc<br>aaggcgattaagttgggtaacgccagggttttcccagtcacgacgttgtaaaacga<br>cggccagtgaattcgagctcggtacccggggatcctctagagattatatttatttc<br>atatatcagaccatgagtggcacagtgaaggaatttgtcgttgaaacagattatcaa<br>gactccttgggaaaagatttagatatggaatatttaacaaaaattcttgctgttcaaga<br>atcgtcgacctgcaggcatgcaagcttggcgtaatcatggtcatagctgtttcctgtg<br>tgaaattgttatccgctcacaattccacacaacatacgagccggaagcataaagtgt<br>aaagcctggggtgcctaatgagtgagctaactcacattaattgcgttgcgctcactg<br>cccgctttccagtcgggaaacctgtcgtgccagctgcattaatgaatcggccaacgc<br>gcggggagaggcggtttgcgtattgggcgctcttccgcttcctcgctcactgactcg<br>ctgcgctcggtcgttcggctgcggcgagcggtatcagctcactcaaaggcggtaat<br>acggttatccacagaatcaggggataacgcaggaaagaacatgtgagcaaaagg<br>ccagcaaaaggccaggaaccgtaaaaaggccgcgttgctggcgtttttccatagg<br>ctccgcccccctgacgagcatcacaaaaatcgacgctcaagtcagaggtggcgaa<br>acccgacaggactataaagataccaggcgtttccccctggaagctccctcgtgcgct<br>ctcctgttccgaccctgccgcttaccggatacctgtccgcctttctcccttcgggaagc<br>gtggcgctttctcatagctcacgctgtaggtatctcagttcggtgtaggtcgttcgctc<br>caagctgggctgtgtgcacgaaccccccgttcagcccgaccgctgcgccttatccg<br>gtaactatcgtcttgagtccaacccggtaagacacgacttatcgccactggcagcag<br>ccactggtaacaggattagcagagcgaggtatgtaggcggtgctacagagttcttg<br>aagtggtggcctaactacggctacactagaagaacagtatttggtatctgcgctctg | Fig 1A |

| | |
|---|---|
| | ctgaagccagttaccttcggaaaaagagttggtagctcttgatccggcaaacaaacc<br>accgctggtagcggtggtttttttgtttgcaagcagcagattacgcgcagaaaaaaa<br>ggatctcaagaagatcctttgatcttttctacggggtctgacgctcagtggaacgaaa<br>actcacgttaagggattttggtcatgagattatcaaaaaggatcttcacctagatcctt<br>ttaaattaaaaatgaagttttaaatcaatctaaagtatatatgagtaaacttggtctgac<br>agttaccaatgcttaatcagtgaggcacctatctcagcgatctgtctatttcgttcatcc<br>atagttgcctgactccccgtcgtgtagataactacgatacgggagggcttaccatct<br>ggccccagtgctgcaatgataccgcgagacccacgctcaccggctccagatttatc<br>agcaataaaccagccagccggaagggccgagcgcagaagtggtcctgcaacttt<br>atccgcctccatccagtctattaattgttgccgggaagctagagtaagtagttcgcca<br>gttaatagtttgcgcaacgttgttgccattgctacaggcatcgtggtgtcacgctcgtc<br>gtttggtatggcttcattcagctccggttcccaacgatcaaggcgagttacatgatcc<br>cccatgttgtgcaaaaaagcggttagctccttcggtcctccgatcgttgtcagaagta<br>agttggccgcagtgttatcactcatggttatggcagcactgcataattctcttactgtc<br>atgccatccgtaagatgcttttctgtgactggtgagtactcaaccaagtcattctgaga<br>atagtgtatgcggcgaccgagttgctcttgcccggcgtcaatacgggataataccg<br>cgccacatagcagaactttaaaagtgctcatcattggaaaacgttcttcggggcgaa<br>aactctcaaggatcttaccgctgttgagatccagttcgatgtaacccactcgtgcacc<br>caactgatcttcagcatcttttactttcaccagcgtttctgggtgagcaaaaacaggaa<br>ggcaaaatgccgcaaaaaagggaataagggcgacacggaaatgttgaatactca<br>tactcttcctttttcaatattattgaagcatttatcagggttattgtctcatgagcggatac<br>atatttgaatgtatttagaaaaataaacaaataggggttccgcgcacatttccccgaa<br>aagtgccacctgacgtctaagaaaccattattatcatgacattaacctataaaaatag<br>gcgtatcacgaggccctttcgtc |
| **Prokaryotic Expression Vector of BvCas12a** | |
| Plasmid-prokaryotic -BvCas12a | tggcgaatgggacgcgccctgtagcggcgcattaagcgcggcgggtgtggtggt<br>tacgcgcagcgtgaccgctacacttgccagcgccctagcgcccgctcctttcgctttc<br>ttcccttcctttctcgccacgttcgccggctttccccgtcaagctctaaatcgggggctc<br>cctttagggttccgatttagtgctttacggcacctcgaccccaaaaaacttgattagg<br>gtgatggttcacgtagtgggccatcgccctgatagacggtttttcgccctttgacgtt<br>ggagtccacgttctttaatagtggactcttgttccaaactggaacaacactcaacccta<br>tctcggtctattcttttgatttataagggattttgccgatttcggcctattggttaaaaaat<br>gagctgatttaacaaaaatttaacgcgaattttaacaaaatattaacgtttacaatttca<br>ggtggcacttttcggggaaatgtgcgcggaacccctatttgtttatttttctaaatacat<br>tcaaatatgtatccgctcatgaattaattcttagaaaaactcatcgagcatcaaatgaa<br>actgcaatttattcatatcaggattatcaataccatatttttgaaaaagccgtttctgtaa<br>tgaaggagaaaactcaccgaggcagttccataggatggcaagatcctggtatcgg<br>tctgcgattccgactcgtccaacatcaatacaacctattaatttcccctcgtcaaaaata<br>aggttatcaagtgagaaatcaccatgagtgacgactgaatccggtgagaatggca<br>aaagtttatgcatttctttccagacttgttcaacaggccagccattacgctcgtcatcaa<br>aatcactcgcatcaaccaaaccgttattcattcgtgattgcgcctgagcgagacgaa<br>atacgcgatcgctgttaaaaggacaattacaaacaggaatcgaatgcaaccggcg<br>caggaacactgccagcgcatcaacaatattttcacctgaatcaggatattcttctaata<br>cctggaatgctgttttcccggggatcgcagtggtgagtaaccatgcatcatcagga |

| | |
|---|---|
| | gtacggataaaatgcttgatggtcggaagaggcataaattccgtcagccagtttagt<br>ctgaccatctcatctgtaacatcattggcaacgctacctttgccatgtttcagaaacaa<br>ctctggcgcatcgggcttcccatacaatcgatagattgtcgcacctgattgcccgaca<br>ttatcgcgagcccatttatacccatataaatcagcatccatgttggaatttaatcgcgg<br>cctagagcaagacgtttcccgttgaatatggctcataacaccccttgtattactgtttat<br>gtaagcagacagttttattgttcatgaccaaaatcccttaacgtgagttttcgttccact<br>gagcgtcagaccccgtagaaaagatcaaaggatcttcttgagatcctttttttctgcg<br>cgtaatctgctgcttgcaaacaaaaaaaccaccgctaccagcggtggtttgtttgcc<br>ggatcaagagctaccaactctttttccgaaggtaactggcttcagcagagcgcagat<br>accaaatactgtccttctagtgtagccgtagttaggccaccacttcaagaactctgta<br>gcaccgcctacatacctcgctctgctaatcctgttaccagtggctgctgccagtggcg<br>ataagtcgtgtcttaccgggttggactcaagacgatagttaccggataaggcgcag<br>cggtcgggctgaacggggggttcgtgcacacagcccagcttggagcgaacgacc<br>tacaccgaactgagatacctacagcgtgagctatgagaaagcgccacgcttcccga<br>agggagaaaggcggacaggtatccggtaagcggcagggtcggaacaggaga<br>gcgcacgagggagcttccagggggaaacgcctggtatctttatagtcctgtcgggt<br>ttcgccacctctgacttgagcgtcgatttttgtgatgctcgtcaggggggcggagcct<br>atggaaaaacgccagcaacgcggcctttttacggttcctggccttttgctggccttttg<br>ctcacatgttctttcctgcgttatcccctgattctgtggataaccgtattaccgcctttga<br>gtgagctgataccgctcgccgcagccgaacgaccgagcgcagcgagtcagtgag<br>cgaggaagcggaagagcgcctgatgcggtattttctccttacgcatctgtgcggtat<br>ttcacaccgcaatggtgcactctcagtacaatctgctctgatgccgcatagttaagcc<br>agtatacactccgctatcgctacgtgactgggtcatggctgcgccccgacacccgcc<br>aacacccgctgacgcgccctgacgggcttgtctgctcccggcatccgcttacagaca<br>agctgtgaccgtctccgggagctgcatgtgtcagaggttttcaccgtcatcaccgaa<br>acgcgcgaggcagctgcggtaaagctcatcagcgtggtcgtgaagcgattcacag<br>atgtctgcctgttcatccgcgtccagctcgttgagtttctccagaagcgttaatgtctg<br>gcttctgataaagcgggccatgttaagggcggttttttcctgtttggtcactgatgcct<br>ccgtgtaagggggatttctgttcatgggggtaatgataccgatgaaacgagagag<br>gatgctcacgatacgggttactgatgatgaacatgcccggttactggaacgttgtga<br>gggtaaacaactggcggtatggatgcggcgggaccagagaaaaatcactcagg<br>gtcaatgccagcgcttcgttaatacagatgtaggtgttccacagggtagccagcag<br>catcctgcgatgcagatccggaacataatggtgcagggcgctgacttccgcgtttcc<br>agactttacgaaacacggaaaccgaagaccattcatgttgttgctcaggtcgcagac<br>gttttgcagcagcagtcgcttcacgttcgctcgcgtatcggtgattcattctgctaacc<br>agtaaggcaaccccgccagcctagccgggtcctcaacgacaggagcacgatcatg<br>cgcacccgtggggccgccatgccggcgataatggcctgcttctcgccgaaacgttt<br>ggtggcgggaccagtgacgaaggcttgagcgagggcgtgcaagattccgaata<br>ccgcaagcgacaggccgatcatcgtcgcgctccagcgaaagcggtcctcgccgaa<br>aatgacccagagcgctgccggcacctgtcctacgagttgcatgataaagaagaca<br>gtcataagtgcggcgacgatagtcatgccccgcgcccaccggaaggagctgactg<br>ggttgaaggctctcaagggcatcggtcgagatcccggtgcctaatgagtgagcta<br>acttacattaattgcgttgcgctcactgcccgctttccagtcgggaaacctgtcgtgcc<br>agctgcattaatgaatcggccaacgcgcggggagaggcggtttgcgtattgggcg |

| | |
|---|---|
| | ccagggtggtttttcttttcaccagtgagacgggcaacagctgattgcccttcaccgc<br>ctggccctgagagagttgcagcaagcggtccacgctggtttgccccagcaggcga<br>aaatcctgtttgatggtggttaacggcgggatataacatgagctgtcttcggtatcgt<br>cgtatcccactaccgagatatccgcaccaacgcgcagcccggactcggtaatggcg<br>cgcattgcgcccagcgccatctgatcgttggcaaccagcatcgcagtgggaacgat<br>gccctcattcagcatttgcatggtttgttgaaaaccggacatggcactccagtcgcctt<br>cccgttccgctatcggctgaatttgattgcgagtgagatatttatgccagccagccag<br>acgcagacgcgccgagacagaacttaatgggcccgctaacagcgcgatttgctgg<br>tgacccaatgcgaccagatgctccacgcccagtcgcgtaccgtcttcatgggagaa<br>aataatactgttgatgggtgtctggtcagagacatcaagaaataacgccggaacatt<br>agtgcaggcagcttccacagcaatggcatcctggtcatccagcggatagttaatgat<br>cagcccactgacgcgttgcgcgagaagattgtgcaccgccgctttacaggcttcga<br>cgccgcttcgttctaccatcgacaccaccacgctggcacccagttgatcggcgcgag<br>atttaatcgccgcgacaatttgcgacggcgcgtgcagggccagactggaggtggc<br>aacgccaatcagcaacgactgtttgcccgccagttgttgtgccacgcggttgggaat<br>gtaattcagctccgccatcgccgcttccactttttcccgcgttttcgcagaaacgtggct<br>ggcctggttcaccacgcgggaaacggtctgataagagacaccggcatactctgcg<br>acatcgtataacgttactggtttcacattcaccaccctgaattgactctcttccgggcgc<br>tatcatgccataccgcgaaaggttttgcgccattcgatggtgtccgggatctcgacg<br>ctctcccttatgcgactcctgcattaggaagcagcccagtagtaggttgaggccgtt<br>gagcaccgccgccgcaaggaatggtgcatgcaaggagatggcgcccaacagtcc<br>cccggccacggggcctgccaccatacccacgccgaaacaagcgctcatgagcccg<br>aagtggcgagcccgatcttccccatcggtgatgtcggcgatataggcgccagcaac<br>cgcacctgtggcgccggtgatgccggccacgatgcgtccggcgtagaggatcga<br>gatctcgatcccgcgaaattaatacgactcactataggggaattgtgagcggataac<br>aattcccctctagaaataattttgtttaactttaagaaggagatataccatgggcagca<br>gccatcatcatcatcatcacagcagcggcctggtgccgcgcggcagccatatgccg<br>aagaagaaacgcaaagtgggcattcatggcgttcctgcggcgCAAGAGCGT<br>AAAAAAATATCGCATCTTACACACAGAAATTCAGTTAAAAAAA<br>CAATTAGGATGCAACTGAATCCTGTAGGTAAAACAATGGATTA<br>TTTTCAAGCAAAGCAAATTCTTGAGAATGATGAAAAGCTTAAA<br>GAGGACTATCAGAAAATCAAGGAAATAGCAGACAGGTTTTAC<br>AGAAATTTAAATGAGGATGTACTTTCAAAAACCGGGTTAGATA<br>AATTAAAAGATTATGCTGAAATTTACTATCACTGTAATACGGAT<br>GCAGACCGAAAAAGACTTAATGAATGTGCATCGGAATTAAGG<br>AAAGAAATCGTCAAAAATTTTAAGAATAGAGATGAGTATAACA<br>AACTATTCAATAAAAAGATGATTGAGATAGTTCTTCCCAAGCAT<br>CTTAAAAATGAGGACGAAAAGGAAGTTGTAGCCTCATTTAAAA<br>ATTTCACAACATACTTTACAGGTTTCTTCACTAACAGAAAAAAT<br>ATGTATTCGGACGGAGAAGAATCCACGGCAATCGCATATAGAT<br>GCATTAACGAAAATTTGCCTAAGCATCTTGACAATGTCAAGGT<br>CTTTGAAAAAGCAATTTCTAAACTATCAAAAAACGCAATTGAT<br>GATTTAGATGCCACTTATTCCGGCTTATGCGGTACAAATTTGTA<br>TGATGTTTTTACAGTTGATTATTTTAACTTTTTGCTTCCACAATCC |

| | |
|---|---|
| | GGAATTACCGAATATAACAAAATCATCGGCGGTTACACAACAA<br>GCGACGGTACAAAAGTTAAGGGTATTAACGAATATATAAATTT<br>GTACAATCAACAAGTATCCAAACGGGATAAAATTCCTAATCTT<br>AAAATTTTGTATAAACAAATTTTAAGTGAGAGTGAAAAGGTATC<br>ATTCATACCGCCAAAGTTTGAAGATGACAACGAACTTTTATCG<br>GCTGTTTCAGAGTTTTACGCAAACGACGAAACCTTTGACGGGA<br>TGCCATTAAAAAAAGCAATTGATGAAACAAAGCTATTATTCGG<br>CAATTTAGATAATTCCTCTCTTAATGGAATTTACATTCAAAATG<br>ACCGATCCGTGACAAATCTGTCAAACAGTATGTTCGGTTCTTG<br>GTCAGTAATAGAAGATTTATGGAACAAAAATTATGACTCCGTT<br>AATTCAAACAGCAGAATCAAAGATATTCAAAAGCGTGAAGAC<br>AAAAGAAAAAAAGCATACAAAGCAGAAAAGAAACTTTCACTTT<br>CATTTTTACAGGTTTTGATTTCCAATTCTGAAAATGATGAAATCA<br>GAAAAAAGTCTATCGTAGATTACTACAAGACTTCTTTAATGCAA<br>CTTACCGACAATTTATCAGACAAATACAAAGAAGCAGCACCTC<br>TGTTCAGTGAAAATTACGATAATGAAAAAGGTTTGAAAAATGA<br>CGATAAATCTATTTCATTAATTAAAAATTTTCTTGATGCCATAAA<br>AGAAATTGAAAAATTCATCAAGCCTTTGTCCGAAACTAATATTA<br>CAGGTGAGAAAAATGATTTGTTTTACAGTCAGTTCACACCATTA<br>CTTGATAATATCAGCAGAATAGACAGATTATATGATAAGGTCA<br>GAAACTATGTTACACAAAAACCGTTTTCAACAGATAAAATCAA<br>GCTTAACTTTGGCAATTCACAGCTATTAAACGGCTGGGATCGA<br>AACAAAGAAAAAGACTGCGGTGCCGTTTTGCTTTGTAAAGATG<br>AAAAGTATTATCTTGCAATCATAGATAAAAGTAATAATAGTATT<br>TTGGAAAATATTGATTTCCAAGACTGCAATGAAAGCGATTATTA<br>TGAAAAGATAGTTTATAAACTTCTGACTAAAATAAGTGGAAAT<br>CTCCCGCGTGTTTTCTTTTCAGAAAAGCACAAGAAACTTTTGTC<br>ACCGTCAGACGAAATACTTAAAATTTATAAAAGCGGCACTTTC<br>AAAAAAGGTGATAAGTTCAGCCTTGATGATTGCCATAAGTTAA<br>TTGATTTCTACAAAGAATCATTCAAAAAGTACCCAAAATGGTTA<br>ATTTATAACTTTAAATTTAAAAACACAAACGAATATAATGATAT<br>CAGCGAATTTTATAATGATGTTGCTTCACAGGGATATAATATTT<br>CAAAAATGAAAATCCCGACATCATTTATTGACAAACTTGTAGAT<br>GAAGGAAAAAATCTATCTTTTCCAACTCTACAACAAAGACTTTTC<br>ACCGCATAGCAAGGGTACTCCTAATCTGCATACACTTTATTTTA<br>AAATGTTGTTTGATGAAAGAAATCTTGAAGATGTGGTGTACAA<br>GCTTAACGGTGAGGCAGAAATGTTTTATCGTCCTGCAAGTATA<br>AAATATGACAAACCAACTCATCCCAAAAACACACCGATAAAA<br>AATAAAAATACACTCAATGATAAAAAAGCAAGCACTTTTCCTT<br>ATGACTTAATTAAAGATAAACGCTACACTAAATGGCAGTTTTCA<br>CTTCACTTCCCTATTACCATGAATTTTAAAGCTCCGGATAGGGC<br>AATGATCAATGATGATGTCAGAAATCTGCTGAAATCCTGCAAC<br>AACAATTTCATCATAGGAATTGACAGAGGCGAAAGAAACTTGC<br>TTTATGTCAGCGTAATTGACAGCAACGGTGCTATAATATATCAG |

| | |
|---|---|
| | CACTCACTCAATATTATCGGAAACAAGTTTAAAGGAAAAACAT<br>ACGAAACTAACTACCGGGAAAAACTTGCAACAAGAGAAAAAG<br>AGCGTACGGAACAGCGCCGTAACTGGAAAGCAATTGAGAGTA<br>TAAAAGAACTCAAAGAGGGCTATATCAGTCAGACTGTGCATGT<br>TATATGTCAGCTTGTTGTCAAGTACGATGCAATCATCGTTATGG<br>AAAAGCTGACTGACGGATTCAAACGAGGCAGAACAAAGTTTG<br>AAAAACAGGTTTATCAGAAATTTGAAAAAATGCTGATTGACAA<br>ACTTAATTACTATGTTGACAAAAAGCTTGATCCCAACGAAGAA<br>GGCGGTTTACTTCATGCCTACCAGCTTACGAACAAGCTTGATA<br>GCTTTGATAAACTTGGCATGCAAAGCGGTTTTATTTTCTATGTTC<br>GTCCTGATTTTACAAGCAAAATTGATCCCGTTACCGGCTTTGTA<br>AATTTGTTGTATCCTCGATATGAAAACATTGACAAAGCCAAAG<br>ATATGATTTCAAGATTTGACGATATAAGATACAATGCCGGCGA<br>GGACTTTTTTGAATTTGACATTGATTACGATAAGTTTCCAAAGA<br>CTGCGTCTGACTATCGCAAGAAGTGGACAATCTGTACTAACGG<br>CGAAAGGATTGAAGCTTTCAGAAATCCCGCAAACAATAACGA<br>ATGGAGTTATCGTACAATAATTCTTGCAGAAAAATTCAAAGAA<br>TTATTTGATAACAATTCTATAAATTATCGTGATTCTGACGATTTG<br>AAAGCTGAAATTCTTTCACAGACAAAGGGCAAATTTTTTGAGG<br>ATTTCTTCAAATTATTAAGACTTACCCTACAGATGCGAAACAGT<br>AACCCTGAAACAGGCGAGGACCGTATCCTTTCTCCCGTCAAGG<br>ACAAAAACGGCAATTTTTACGACAGTTCAAAATATGATGAAAA<br>GAGCAAGCTTCCGTGTGATGCCGATGCAAACGGTGCGTACAA<br>CATTGCCCGCAAAGGTTTGTGGATTGTTGAACAATTCAAAAAA<br>GCCGATAATGTTTCAGCTGTCGAACCGGTAATCCACAATGACA<br>AATGGCTGAAATTTGTTCAGGAGAATGATATGGCGAATAATaaa<br>cgcccggcggcgactaaaaaagcaggccaggcgaaaaaaaagaagtaagaatt<br>cgagctccgtcgacaagcttgcggccgcactcgagcaccaccaccaccaccactga<br>gatccggctgctaacaaagcccgaaaggaagctgagttggctgctgccaccgctg<br>agcaataactagcataaccccttggggcctctaaacgggtcttgaggggttttttgct<br>gaaaggaggaactatatccggat |
| Plasmid-prokaryotic -BvCas12a-R | tggcgaatgggacgcgccctgtagcggcgcattaagcgcggcgggtgtggtggt<br>tacgcgcagcgtgaccgctacacttgccagcgccctagcgcccgctcctttcgctttc<br>ttcccttcctttctcgccacgttcgccggctttccccgtcaagctctaaatcgggggctc<br>cctttagggttccgatttagtgctttacggcacctcgaccccaaaaaacttgattagg<br>gtgatggttcacgtagtgggccatcgccctgatagacggtttttcgccctttgacgtt<br>ggagtccacgttctttaatagtggactcttgttccaaactggaacaacactcaaccccta<br>tctcggtctattcttttgatttataagggattttgccgatttcggcctattggttaaaaaat<br>gagctgatttaacaaaaatttaacgcgaattttaacaaaatattaacgtttacaatttca<br>ggtggcacttttcggggaaatgtgcgcggaacccctatttgtttatttttctaaatacat<br>tcaaatatgtatccgctcatgaattaattcttagaaaaactcatcgagcatcaaatgaa<br>actgcaatttattcatatcaggattatcaataccatatttttgaaaaagccgtttctgtaa<br>tgaaggagaaaactcaccgaggcagttccataggatggcaagatcctggtatcgg<br>tctgcgattccgactcgtccaacatcaatacaacctattaatttcccctcgtcaaaaata |

| | |
|---|---|
| | aggttatcaagtgagaaatcaccatgagtgacgactgaatccggtgagaatggca<br>aaagtttatgcatttctttccagacttgttcaacaggccagccattacgctcgtcatcaa<br>aatcactcgcatcaaccaaaccgttattcattcgtgattgcgcctgagcgagacgaa<br>atacgcgatcgctgttaaaaggacaattacaaacaggaatcgaatgcaaccggcg<br>caggaacactgccagcgcatcaacaatattttcacctgaatcaggatattcttctaata<br>cctggaatgctgttttcccggggatcgcagtggtgagtaaccatgcatcatcagga<br>gtacggataaaatgcttgatggtcggaagaggcataaattccgtcagccagtttagt<br>ctgaccatctcatctgtaacatcattggcaacgctacctttgccatgtttcagaaacaa<br>ctctggcgcatcgggcttcccatacaatcgatagattgtcgcacctgattgcccgaca<br>ttatcgcgagcccatttatacccatataaatcagcatccatgttggaatttaatcgcgg<br>cctagagcaagacgtttcccgttgaatatggctcataacaccccttgtattactgtttat<br>gtaagcagacagttttattgttcatgaccaaaatcccttaacgtgagttttcgttccact<br>gagcgtcagaccccgtagaaaagatcaaaggatcttcttgagatcctttttttctgcg<br>cgtaatctgctgcttgcaaacaaaaaaaccaccgctaccagcggtggtttgtttgcc<br>ggatcaagagctaccaactctttttccgaaggtaactggcttcagcagagcgcagat<br>accaaatactgtccttctagtgtagccgtagttaggccaccacttcaagaactctgta<br>gcaccgcctacatacctcgctctgctaatcctgttaccagtggctgctgccagtggcg<br>ataagtcgtgtcttaccgggttggactcaagacgatagttaccggataaggcgcag<br>cggtcgggctgaacggggggttcgtgcacacagcccagcttggagcgaacgacc<br>tacaccgaactgagatacctacagcgtgagctatgagaaagcgccacgcttcccga<br>agggagaaaggcggacaggtatccggtaagcggcagggtcggaacaggaga<br>gcgcacgagggagcttccagggggaaacgcctggtatctttatagtcctgtcgggt<br>ttcgccacctctgacttgagcgtcgatttttgtgatgctcgtcaggggggcggagcct<br>atggaaaaacgccagcaacgcggcctttttacggttcctggccttttgctggccttttg<br>ctcacatgttctttcctgcgttatcccctgattctgtggataaccgtattaccgcctttga<br>gtgagctgataccgctcgccgcagccgaacgaccgagcgcagcgagtcagtgag<br>cgaggaagcggaagagcgcctgatgcggtattttctccttacgcatctgtgcggtat<br>ttcacaccgcaatggtgcactctcagtacaatctgctctgatgccgcatagttaagcc<br>agtatacactccgctatcgctacgtgactgggtcatggctgcgccccgacacccgcc<br>aacacccgctgacgcgccctgacgggcttgtctgctcccggcatccgcttacagaca<br>agctgtgaccgtctccgggagctgcatgtgtcagaggttttcaccgtcatcaccgaa<br>acgcgcgaggcagctgcggtaaagctcatcagcgtggtcgtgaagcgattcacag<br>atgtctgcctgttcatccgcgtccagctcgttgagtttctccagaagcgttaatgtctg<br>gcttctgataaagcgggccatgttaagggcggttttttcctgtttggtcactgatgcct<br>ccgtgtaagggggatttctgttcatgggggtaatgataccgatgaaacgagagag<br>gatgctcacgatacgggttactgatgatgaacatgcccggttactggaacgttgtga<br>gggtaaacaactggcggtatggatgcggcgggaccagagaaaaatcactcagg<br>gtcaatgccagcgcttcgttaatacagatgtaggtgttccacagggtagccagcag<br>catcctgcgatgcagatccggaacataatggtgcagggcgctgacttccgcgtttcc<br>agactttacgaaacacggaaaccgaagaccattcatgttgttgctcaggtcgcagac<br>gttttgcagcagcagtcgcttcacgttcgctcgcgtatcggtgattcattctgctaacc<br>agtaaggcaaccccgccagcctagccgggtcctcaacgacaggagcacgatcatg<br>cgcacccgtggggccgccatgccggcgataatggcctgcttctcgccgaaacgttt<br>ggtggcgggaccagtgacgaaggcttgagcgagggcgtgcaagattccgaata |

| | |
|---|---|
| | ccgcaagcgacaggccgatcatcgtcgcgctccagcgaaagcggtcctcgccgaa<br>aatgacccagagcgctgccggcacctgtcctacgagttgcatgataaagaagaca<br>gtcataagtgcggcgacgatagtcatgccccgcgcccaccggaaggagctgactg<br>ggttgaaggctctcaagggcatcggtcgagatcccggtgcctaatgagtgagcta<br>acttacattaattgcgttgcgctcactgcccgctttccagtcgggaaacctgtcgtgcc<br>agctgcattaatgaatcggccaacgcgcggggagaggcggtttgcgtattgggcg<br>ccagggtggtttttcttttcaccagtgagacgggcaacagctgattgcccttcaccgc<br>ctggccctgagagagttgcagcaagcggtccacgctggtttgccccagcaggcga<br>aaatcctgtttgatggtggttaacggcgggatataacatgagctgtcttcggtatcgt<br>cgtatcccactaccgagatatccgcaccaacgcgcagcccggactcggtaatggcg<br>cgcattgcgcccagcgccatctgatcgttggcaaccagcatcgcagtgggaacgat<br>gccctcattcagcatttgcatggtttgttgaaaaccggacatggcactccagtcgcctt<br>cccgttccgctatcggctgaatttgattgcgagtgagatatttatgccagccagccag<br>acgcagacgcgccgagacagaacttaatgggcccgctaacagcgcgatttgctgg<br>tgacccaatgcgaccagatgctccacgcccagtcgcgtaccgtcttcatgggagaa<br>aataatactgttgatgggtgtctggtcagagacatcaagaaataacgccggaacatt<br>agtgcaggcagcttccacagcaatggcatcctggtcatccagcggatagttaatgat<br>cagcccactgacgcgttgcgcgagaagattgtgcaccgccgctttacaggcttcga<br>cgccgcttcgttctaccatcgacaccaccacgctggcacccagttgatcggcgcgag<br>atttaatcgccgcgacaatttgcgacggcgcgtgcagggccagactggaggtggc<br>aacgccaatcagcaacgactgtttgcccgccagttgttgtgccacgcggttgggaat<br>gtaattcagctccgccatcgccgcttccactttttcccgcgttttcgcagaaacgtggct<br>ggcctggttcaccacgcgggaaacggtctgataagagacaccggcatactctgcg<br>acatcgtataacgttactggtttcacattcaccaccctgaattgactctcttccgggcgc<br>tatcatgccataccgcgaaaggttttgcgccattcgatggtgtccgggatctcgacg<br>ctctcccttatgcgactcctgcattaggaagcagcccagtagtaggttgaggccgtt<br>gagcaccgccgccgcaaggaatggtgcatgcaaggagatggcgcccaacagtcc<br>cccggccacggggcctgccaccatacccacgccgaaacaagcgctcatgagcccg<br>aagtggcgagcccgatcttccccatcggtgatgtcggcgatataggcgccagcaac<br>cgcacctgtggcgccggtgatgccggccacgatgcgtccggcgtagaggatcga<br>gatctcgatcccgcgaaattaatacgactcactataggggaattgtgagcggataac<br>aattcccctctagaaataattttgtttaactttaagaaggagatataccatgggcagca<br>gccatcatcatcatcatcacagcagcggcctggtgccgcgcggcagccatatgccg<br>aagaagaaacgcaaagtgggcattcatggcgttcctgcggcgcaagagcgtaaa<br>aaaatatcgcatcttacacacagaaattcagttaaaaaaacaattaggatgcaactg<br>aatcctgtaggtaaaacaatggattattttcaagcaaagcaaattcttgagaatgatg<br>aaaagcttaaagaggactatcagaaaatcaaggaaatagcagacaggttttacag<br>aaatttaaatgaggatgtactttcaaaaaccgggttagataaattaaaagattatgct<br>gaaatttactatcactgtaatacggatgcagaccgaaaaagacttaatgaatgtgca<br>tcggaattaaggaaagaaatcgtcaaaaattttaagaatagagatgagtataacaa<br>actattcaataaaaagatgattgagatagttcttcccaagcatcttaaaaatgaggac<br>gaaaaggaagttgtagcctcatttaaaaatttcacaacatactttacaggtttcttcact<br>aacagaaaaaatatgtattcggacggagaagaatccacggcaatcgcatatagat<br>gcattaacgaaaatttgcctaagcatcttgacaatgtcaaggtctttgaaaaagcaat |

| | |
|---|---|
| | ttctaaactatcaaaaaacgcaattgatgatttagatgccacttattccggcttatgcg<br>gtacaaatttgtatgatgtttttacagttgattattttaactttttgcttccacaatccgga<br>attaccgaatataacaaaatcatcggcggttacacaacaagcgacggtacaaaagt<br>taagggtattaacgaatatataaatttgtacaatcaacaagtatccaaacgggataaa<br>attcctaatcttaaaattttgtataaacaaattttaagtgagagtgaaaaggtatcattc<br>ataccgccaaagtttgaagatgacaacgaacttttatcggctgtttcagagttttacgc<br>aaacgacgaaacctttgacgggatgccattaaaaaaagcaattgatgaaacaaag<br>ctattattcggcaatttagataattcctctcttaatggaatttacattcaaaatgaccgat<br>ccgtgacaaatctgtcaaacagtatgttcggttcttggtcagtaatagaagatttatg<br>gaacaaaaattatgactccgttaattcaaacagcagaatcaaagatattcaaaagcg<br>tgaagacaaaagaaaaaaagcatacaaagcagaaaagaaactttcactttcattttt<br>acaggttttgatttccaattctgaaaatgatgaaatcagaaaaaagtctatcgtagatt<br>actacaagacttctttaatgcaacttaccgacaatttatcagacaaatacaaagaagc<br>agcacctctgttcagtgaaaattacgataatgaaaaaggtttgaaaaatgacgataa<br>atctatttcattaattaaaaattttcttgatgccataaaagaaattgaaaaattcatcaag<br>cctttgtccgaaactaatattacaggtgagaaaaatgatttgttttacagtcagttcaca<br>ccattacttgataatatcagcagaatagacagattatatgataaggtcagaaactatg<br>ttacacaaaaaccgttttcaacagataaaatcaagcttaactttggcaattcacagcta<br>ttacgcggctgggatcgaaacaaagaaaaagactgcggtgccgttttgctttgtaa<br>agatgaaaagtattatcttgcaatcatagataaaagtaataatagtattttggaaaata<br>ttgatttccaagactgcaatgaaagcgattattatgaaaagatagtttataaacttctg<br>ccgaaaataagtggaaatctcccgcgtgttttcttttcagaaaagcacaagaaacttttt<br>gtcaccgtcagacgaaatacttaaaatttataaaagcggcactttcaaaaaaggtga<br>taagttcagccttgatgattgccataagttaattgatttctacaaagaatcattcaaaaa<br>gtacccaaaatggttaatttataactttaaatttaaaaacacaaacgaatataatgatat<br>cagcgaattttataatgatgttgcttcacagggatataatatttcaaaaatgaaaatcc<br>cgacatcatttattgacaaacttgtagatgaaggaaaaatctatcttttccaactctaca<br>acaaagacttttcaccgcatagcaagggtactcctaatctgcatacactttattttaaaa<br>tgttgtttgatgaaagaaatcttgaagatgtggtgtacaagcttaacggtgaggcag<br>aaatgttttatcgtcctgcaagtataaaatatgacaaaccaactcatcccaaaaacac<br>accgataaaaaataaaaatacactcaatgataaaaaagcaagcacttttccttatgac<br>ttaattaaagataaacgctacactaaatggcagttttcacttcacttccctattaccatg<br>aattttaaagctccggatagggcaatgatcaatgatgatgtcagaaatctgctgaaa<br>tcctgcaacaacaatttcatcataggaattgacagaggcgaaagaaacttgctttatg<br>tcagcgtaattgacagcaacggtgctataatatatcagcactcactcaatattatcgg<br>aaacaagtttaaaggaaaaacatacgaaactaactaccgggaaaaacttgcaaca<br>agagaaaaagagcgtacggaacagcgccgtaactggaaagcaattgagagtata<br>aaagaactcaaagagggctatatcagtcagactgtgcatgttatatgtcagcttgttg<br>tcaagtacgatgcaatcatcgttatggaaaagctgactgacggattcaaacgaggc<br>agaacaaagtttgaaaaacaggtttatcagaaatttgaaaaaatgctgattgacaaa<br>cttaattactatgttgacaaaaagcttgatcccaacgaagaaggcggtttacttcatg<br>cctaccagcttacgaacaagcttgatagctttgataaacttggcatgcaaagcggttt<br>tattttctatgttcgtcctgattttacaagcaaaattgatcccgttaccggctttgtaaatt<br>tgttgtatcctcgatatgaaaacattgacaaagccaaagatatgatttcaagatttgac |

| | |
|---|---|
| | gatataagatacaatgccggcgaggactttttgaatttgacattgattacgataagtt<br>tccaaagactgcgtctgactatcgcaagaagtggacaatctgtactaacggcgaaa<br>ggattgaagctttcagaaatcccgcaaacaataacgaatggagttatcgtacaataa<br>ttcttgcagaaaaattcaaagaattatttgataacaattctataaattatcgtgattctga<br>cgatttgaaagctgaaattctttcacagacaaagggcaaatttttgaggatttcttca<br>aattattaagacttaccctacagatgcgaaacagtaaccctgaaacaggcgaggac<br>cgtatcctttctcccgtcaaggacaaaaacggcaatttttacgacagttcaaaatatga<br>tgaaaagagcaagcttccgtgtgatgccgatgcaaacggtgcgtacaacattgccc<br>gcaaaggtttgtggattgttgaacaattcaaaaaagccgataatgtttcagctgtcga<br>accggtaatccacaatgacaaatggctgaaatttgttcaggagaatgatatggcga<br>ataataaacgcccggcggcgactaaaaaagcaggccaggcgaaaaaaaagaag<br>taagaattcgagctccgtcgacaagcttgcggccgcactcgagcaccaccaccacc<br>accactgagatccggctgctaacaaagcccgaaaggaagctgagttggctgctgc<br>caccgctgagcaataactagcataaccccttggggcctctaaacgggtcttgaggg<br>gtttttgctgaaaggaggaactatatccggat |
| Plasmid-prokaryotic - BvCas12a-RVR | tggcgaatgggacgcgccctgtagcggcgcattaagcgcggcgggtgtggtggt<br>tacgcgcagcgtgaccgctacacttgccagcgccctagcgcccgctcctttcgctttc<br>ttcccttcctttctcgccacgttcgccggctttccccgtcaagctctaaatcgggggctc<br>cctttagggttccgatttagtgctttacggcacctcgaccccaaaaaacttgattagg<br>gtgatggttcacgtagtgggccatcgccctgatagacggtttttcgccctttgacgtt<br>ggagtccacgttctttaatagtggactcttgttccaaactggaacaacactcaaccta<br>tctcggtctattcttttgatttataagggattttgccgatttcggcctattggttaaaaaat<br>gagctgatttaacaaaaatttaacgcgaattttaacaaaatattaacgtttacaatttca<br>ggtggcacttttcggggaaatgtgcgcggaacccctatttgtttatttttctaaatacat<br>tcaaatatgtatccgctcatgaattaattcttagaaaaactcatcgagcatcaaatgaa<br>actgcaatttattcatatcaggattatcaataccatatttttgaaaaagccgtttctgtaa<br>tgaaggagaaaactcaccgaggcagttccataggatggcaagatcctggtatcgg<br>tctgcgattccgactcgtccaacatcaatacaacctattaatttcccctcgtcaaaaata<br>aggttatcaagtgagaaatcaccatgagtgacgactgaatccggtgagaatggca<br>aaagtttatgcatttctttccagacttgttcaacaggccagccattacgctcgtcatcaa<br>aatcactcgcatcaaccaaaccgttattcattcgtgattgcgcctgagcgagacgaa<br>atacgcgatcgctgttaaaaggacaattacaaacaggaatcgaatgcaaccggcg<br>caggaacactgccagcgcatcaacaatattttcacctgaatcaggatattcttctaata<br>cctggaatgctgttttcccggggatcgcagtggtgagtaaccatgcatcatcagga<br>gtacggataaaatgcttgatggtcggaagaggcataaattccgtcagccagtttagt<br>ctgaccatctcatctgtaacatcattggcaacgctacctttgccatgtttcagaaacaa<br>ctctggcgcatcgggcttcccatacaatcgatagattgtcgcacctgattgcccgaca<br>ttatcgcgagcccatttatacccatataaatcagcatccatgttggaatttaatcgcgg<br>cctagagcaagacgtttcccgttgaatatggctcataacaccccttgtattactgtttat<br>gtaagcagacagttttattgttcatgaccaaaatcccttaacgtgagttttcgttccact<br>gagcgtcagaccccgtagaaaagatcaaaggatcttcttgagatcctttttttctgcg<br>cgtaatctgctgcttgcaaacaaaaaaaccaccgctaccagcggtggtttgtttgcc<br>ggatcaagagctaccaactctttttccgaaggtaactggcttcagcagagcgcagat<br>accaaatactgtccttctagtgtagccgtagttaggccaccacttcaagaactctgta |

| | |
|---|---|
| | gcaccgcctacatacctcgctctgctaatcctgttaccagtggctgctgccagtggcg<br>ataagtcgtgtcttaccgggttggactcaagacgatagttaccggataaggcgcag<br>cggtcgggctgaacggggggttcgtgcacacagcccagcttggagcgaacgacc<br>tacaccgaactgagatacctacagcgtgagctatgagaaagcgccacgcttcccga<br>agggagaaaggcggacaggtatccggtaagcggcagggtcggaacaggaga<br>gcgcacgagggagcttccagggggaaacgcctggtatctttatagtcctgtcgggt<br>ttcgccacctctgacttgagcgtcgatttttgtgatgctcgtcaggggggcggagcct<br>atggaaaaacgccagcaacgcggcctttttacggttcctggccttttgctggccttttg<br>ctcacatgttctttcctgcgttatcccctgattctgtggataaccgtattaccgcctttga<br>gtgagctgataccgctcgccgcagccgaacgaccgagcgcagcgagtcagtgag<br>cgaggaagcggaagagcgcctgatgcggtattttctccttacgcatctgtgcggtat<br>ttcacaccgcaatggtgcactctcagtacaatctgctctgatgccgcatagttaagcc<br>agtatacactccgctatcgctacgtgactgggtcatggctgcgccccgacacccgcc<br>aacacccgctgacgcgccctgacgggcttgtctgctcccggcatccgcttacagaca<br>agctgtgaccgtctccgggagctgcatgtgtcagaggttttcaccgtcatcaccgaa<br>acgcgcgaggcagctgcggtaaagctcatcagcgtggtcgtgaagcgattcacag<br>atgtctgcctgttcatccgcgtccagctcgttgagtttctccagaagcgttaatgtctg<br>gcttctgataaagcgggccatgttaagggcggttttttcctgtttggtcactgatgcct<br>ccgtgtaagggggatttctgttcatgggggtaatgataccgatgaaacgagagag<br>gatgctcacgatacgggttactgatgatgaacatgcccggttactggaacgttgtga<br>gggtaaacaactggcggtatggatgcggcgggaccagagaaaaatcactcagg<br>gtcaatgccagcgcttcgttaatacagatgtaggtgttccacagggtagccagcag<br>catcctgcgatgcagatccggaacataatggtgcagggcgctgacttccgcgtttcc<br>agactttacgaaacacggaaaccgaagaccattcatgttgttgctcaggtcgcagac<br>gttttgcagcagcagtcgcttcacgttcgctcgcgtatcggtgattcattctgctaacc<br>agtaaggcaaccccgccagcctagccgggtcctcaacgacaggagcacgatcatg<br>cgcacccgtggggccgccatgccggcgataatggcctgcttctcgccgaaacgttt<br>ggtggcgggaccagtgacgaaggcttgagcgagggcgtgcaagattccgaata<br>ccgcaagcgacaggccgatcatcgtcgcgctccagcgaaagcggtcctcgccgaa<br>aatgacccagagcgctgccggcacctgtcctacgagttgcatgataaagaagaca<br>gtcataagtgcggcgacgatagtcatgccccgcgcccaccggaaggagctgactg<br>ggttgaaggctctcaagggcatcggtcgagatcccggtgcctaatgagtgagcta<br>acttacattaattgcgttgcgctcactgcccgctttccagtcgggaaacctgtcgtgcc<br>agctgcattaatgaatcggccaacgcgcggggagaggcggtttgcgtattgggcg<br>ccagggtggtttttcttttcaccagtgagacgggcaacagctgattgcccttcaccgc<br>ctggccctgagagagttgcagcaagcggtccacgctggtttgccccagcaggcga<br>aaatcctgtttgatggtggttaacggcgggatataacatgagctgtcttcggtatcgt<br>cgtatcccactaccgagatatccgcaccaacgcgcagcccggactcggtaatggcg<br>cgcattgcgcccagcgccatctgatcgttggcaaccagcatcgcagtgggaacgat<br>gccctcattcagcatttgcatggtttgttgaaaaccggacatggcactccagtcgcctt<br>cccgttccgctatcggctgaatttgattgcgagtgagatatttatgccagccagccag<br>acgcagacgcgccgagacagaacttaatgggcccgctaacagcgcgatttgctgg<br>tgacccaatgcgaccagatgctccacgcccagtcgcgtaccgtcttcatgggagaa<br>aataatactgttgatgggtgtctggtcagagacatcaagaaataacgccggaacatt |

| | |
|---|---|
| | agtgcaggcagcttccacagcaatggcatcctggtcatccagcggatagttaatgat<br>cagcccactgacgcgttgcgcgagaagattgtgcaccgccgctttacaggcttcga<br>cgccgcttcgttctaccatcgacaccaccacgctggcacccagttgatcggcgcgag<br>atttaatcgccgcgacaatttgcgacggcgcgtgcagggccagactggaggtggc<br>aacgccaatcagcaacgactgtttgcccgccagttgttgtgccacgcggttgggaat<br>gtaattcagctccgccatcgccgcttccactttttcccgcgttttcgcagaaacgtggct<br>ggcctggttcaccacgcgggaaacggtctgataagagacaccggcatactctgcg<br>acatcgtataacgttactggtttcacattcaccaccctgaattgactctcttccgggcgc<br>tatcatgccataccgcgaaaggttttgcgccattcgatggtgtccgggatctcgacg<br>ctctcccttatgcgactcctgcattaggaagcagcccagtagtaggttgaggccgtt<br>gagcaccgccgccgcaaggaatggtgcatgcaaggagatggcgcccaacagtcc<br>cccggccacggggcctgccaccatacccacgccgaaacaagcgctcatgagcccg<br>aagtggcgagcccgatcttccccatcggtgatgtcggcgatataggcgccagcaac<br>cgcacctgtggcgccggtgatgccggccacgatgcgtccggcgtagaggatcga<br>gatctcgatcccgcgaaattaatacgactcactataggggaattgtgagcggataac<br>aattcccctctagaaataattttgtttaactttaagaaggagatataccatgggcagca<br>gccatcatcatcatcatcacagcagcggcctggtgccgcgcggcagccatatgccg<br>aagaagaaacgcaaagtgggcattcatggcgttcctgcggcgcaagagcgtaaa<br>aaaatatcgcatcttacacacagaaattcagttaaaaaaacaattaggatgcaactg<br>aatcctgtaggtaaaacaatggattattttcaagcaaagcaaattcttgagaatgatg<br>aaaagcttaaagaggactatcagaaaatcaaggaaatagcagacaggttttacag<br>aaatttaaatgaggatgtactttcaaaaaccgggttagataaattaaaagattatgct<br>gaaatttactatcactgtaatacggatgcagaccgaaaaagacttaatgaatgtgca<br>tcggaattaaggaaagaaatcgtcaaaaattttaagaatagagatgagtataacaa<br>actattcaataaaaagatgattgagatagttcttcccaagcatcttaaaaatgaggac<br>gaaaaggaagttgtagcctcatttaaaaatttcacaacatactttacaggtttcttcact<br>aacagaaaaaatatgtattcggacggagaagaatccacggcaatcgcatatagat<br>gcattaacgaaaatttgcctaagcatcttgacaatgtcaaggtctttgaaaaagcaat<br>ttctaaactatcaaaaaacgcaattgatgatttagatgccacttattccggcttatgcg<br>gtacaaatttgtatgatgtttttacagttgattattttaactttttgcttccacaatccgga<br>attaccgaatataacaaaatcatcggcggttacacaacaagcgacggtacaaaagt<br>taagggtattaacgaatatataaatttgtacaatcaacaagtatccaaacgggataaa<br>attcctaatcttaaaattttgtataaacaaattttaagtgagagtgaaaaggtatcattc<br>ataccgccaaagtttgaagatgacaacgaacttttatcggctgtttcagagttttacgc<br>aaacgacgaaacctttgacgggatgccattaaaaaaagcaattgatgaaacaaag<br>ctattattcggcaatttagataattcctctcttaatggaatttacattcaaaatgaccgat<br>ccgtgacaaatctgtcaaacagtatgttcggttcttggtcagtaatagaagatttatg<br>gaacaaaaattatgactccgttaattcaaacagcagaatcaaagatattcaaaagcg<br>tgaagacaaaagaaaaaaagcatacaaagcagaaaagaaactttcactttcattttt<br>acaggttttgatttccaattctgaaaatgatgaaatcagaaaaaagtctatcgtagatt<br>actacaagacttctttaatgcaacttaccgacaatttatcagacaaatacaaagaagc<br>agcacctctgttcagtgaaaattacgataatgaaaaaggtttgaaaaatgacgataa<br>atctatttcattaattaaaaattttcttgatgccataaaagaaattgaaaaattcatcaag<br>cctttgtccgaaactaatattacaggtgagaaaaatgatttgttttacagtcagttcaca |

| | |
|---|---|
| | ccattacttgataatatcagcagaatagacagattatatgataaggtcagaaactatg<br>ttacacaaaaaccgttttcaacagataaaatcaagcttaactttggcaattcacagcta<br>ttacgcggctgggatcgaaacgttgaaaaagaccgcggtgccgttttgctttgtaaa<br>gatgaaaagtattatcttgcaatcatagataaaagtaataatagtattttggaaaatat<br>tgatttccaagactgcaatgaaagcgattattatgaaaagatagtttataaacttctgc<br>cgaaaataagtggaaatctcccgcgtgttttcttttcagaaaagcacaagaaactttt<br>gtcaccgtcagacgaaatacttaaaatttataaaagcggcactttcaaaaaaggtga<br>taagttcagccttgatgattgccataagttaattgatttctacaaagaatcattcaaaaa<br>gtacccaaaatggttaatttataactttaaatttaaaaacacaaacgaatataatgatat<br>cagcgaattttataatgatgttgcttcacagggatataatatttcaaaaatgaaaatcc<br>cgacatcatttattgacaaacttgtagatgaaggaaaaatctatcttttccaactctaca<br>acaaagacttttcaccgcatagcaagggtactcctaatctgcatacactttattttaaaa<br>tgttgtttgatgaaagaaatcttgaagatgtggtgtacaagcttaacggtgaggcag<br>aaatgttttatcgtcctgcaagtataaaatatgacaaaccaactcatcccaaaaacac<br>accgataaaaaataaaaatacactcaatgataaaaaagcaagcacttttccttatgac<br>ttaattaaagataaacgctacactaaatggcagttttcacttcacttccctattaccatg<br>aattttaaagctccggatagggcaatgatcaatgatgatgtcagaaatctgctgaaa<br>tcctgcaacaacaatttcatcataggaattgacagaggcgaaagaaacttgctttatg<br>tcagcgtaattgacagcaacggtgctataatatatcagcactcactcaatattatcgg<br>aaacaagtttaaaggaaaaacatacgaaactaactaccgggaaaaacttgcaaca<br>agagaaaaagagcgtacggaacagcgccgtaactggaaagcaattgagagtata<br>aaagaactcaaagagggctatatcagtcagactgtgcatgttatatgtcagcttgttg<br>tcaagtacgatgcaatcatcgttatggaaaagctgactgacggattcaaacgaggc<br>agaacaaagtttgaaaaacaggtttatcagaaatttgaaaaaatgctgattgacaaa<br>cttaattactatgttgacaaaaagcttgatcccaacgaagaaggcggtttacttcatg<br>cctaccagcttacgaacaagcttgatagctttgataaacttggcatgcaaagcggttt<br>tattttctatgttcgtcctgattttacaagcaaaattgatcccgttaccggctttgtaaatt<br>tgttgtatcctcgatatgaaaacattgacaaagccaaagatatgatttcaagatttgac<br>gatataagatacaatgccggcgaggactttttgaatttgacattgattacgataagtt<br>tccaaagactgcgtctgactatcgcaagaagtggacaatctgtactaacggcgaaa<br>ggattgaagctttcagaaatcccgcaaacaataacgaatggagttatcgtacaataa<br>ttcttgcagaaaaattcaaagaattatttgataacaattctataaattatcgtgattctga<br>cgatttgaaagctgaaattctttcacagacaaagggcaaatttttgaggatttcttca<br>aattattaagacttaccctacagatgcgaaacagtaaccctgaaacaggcgaggac<br>cgtatcctttctcccgtcaaggacaaaaacggcaatttttacgacagttcaaaatatga<br>tgaaaagagcaagcttccgtgtgatgccgatgcaaacggtgcgtacaacattgccc<br>gcaaaggtttgtggattgttgaacaattcaaaaaagccgataatgtttcagctgtcga<br>accggtaatccacaatgacaaatggctgaaatttgttcaggagaatgatatggcga<br>ataataaacgcccggcggcgactaaaaaagcaggccaggcgaaaaaaaagaag<br>taagaattcgagctccgtcgacaagcttgcggccgcactcgagcaccaccaccacc<br>accactgagatccggctgctaacaaagcccgaaaggaagctgagttggctgctgc<br>caccgctgagcaataactagcataaccccttggggcctctaaacgggtcttgaggg<br>gtttttttgctgaaaggaggaactatatccggat |
| **Eukaryote Expression Vector of BvCas12a** | |

| | |
|---|---|
| Plasmid-eukaryote BvCas12a-NC | gagggcctatttcccatgattccttcatatttgcatatacgatacaaggctgttagaga<br>gataattggaattaatttgactgtaaacacaaagatattagtacaaaatacgtgacgt<br>agaaagtaataatttcttgggtagtttgcagttttaaaattatgttttaaaatggactatc<br>atatgcttaccgtaacttgaaagtatttcgatttcttggctttatatatcttgtggaaagg<br>acgaaacaccgAATTTCTACTATTGTAGATgggtcttcgagaagaccttttt<br>ttgttttagagctagaaatagcaagttaaaataaggctagtccgtttttagcgcgtgc<br>gccaattctgcagacaaatggctctagaggtacccgttacataacttacggtaaatg<br>gcccgcctggctgaccgcccaacgacccccgcccattgacgtcaatagtaacgcca<br>atagggactttccattgacgtcaatgggtggagtatttacggtaaactgcccacttgg<br>cagtacatcaagtgtatcatatgccaagtacgccccctattgacgtcaatgacggtaa<br>atggcccgcctggcattgtgcccagtacatgaccttatgggactttcctacttggcag<br>tacatctacgtattagtcatcgctattaccatggtcgaggtgagccccacgttctgcttc<br>actctccccatctcccccccctccccacccccaattttgtatttatttattttttaattattttg<br>tgcagcgatgggggcgggggggggggggggcgcgcgccaggcggggcg<br>gggcggggcgaggggcggggcggggcgaggcggagaggtgcggcggcag<br>ccaatcagagcggcgcgctccgaaagtttccttttatggcgaggcggcggcggcg<br>gcggccctataaaaagcgaagcgcgcggcgggcgggagtcgctgcgcgctgcc<br>ttcgccccgtgccccgctccgccgccgcctcgcgccgcccgccccggctctgactga<br>ccgcgttactcccacaggtgagcgggcgggacggcccttctcctccgggctgtaat<br>tagctgagcaagaggtaagggtttaagggatggttggttggtggggtattaatgtt<br>taattacctggagcacctgcctgaaatcactttttttcaggttggaccggtgccaccat<br>ggactataaggaccacgacggagactacaaggatcatgatattgattacaaagac<br>gatgacgataagatggccccaaagaagaagcggaaggtcggtatccacggagtc<br>ccagcagccATGCAGGAGAGAAAGAAGATCAGCCACCTGACCCA<br>CAGAAACAGCGTGAAGAAAACCATCAGAATGCAGCTGAACCC<br>CGTGGGAAAGACCATGGACTACTTCCAGGCCAAGCAGATCCT<br>GGAGAACGACGAGAAGCTGAAGGAGGACTACCAGAAGATCA<br>AGGAGATCGCCGACAGATTCTACAGAAACCTGAACGAGGACG<br>TGCTGAGCAAAACCGGACTGGACAAGCTGAAGGACTACGCCG<br>AGATCTACTACCATTGCAACACCGACGCCGACAGAAAGAGAC<br>TGAACGAGTGCGCCAGCGAGCTGAGAAAGGAGATCGTGAAG<br>AACTTCAAGAACAGAGATGAGTACAACAAGCTGTTCAACAAG<br>AAGATGATCGAGATCGTGCTGCCCAAGCACCTGAAGAACGAG<br>GACGAGAAGGAAGTGGTGGCCAGCTTCAAGAACTTCACCACC<br>TACTTCACCGGCTTCTTCACCAACAGAAAGAACATGTACAGCG<br>ACGGCGAAGAGTCTACCGCTATTGCCTACAGATGCATCAACGA<br>GAACCTGCCCAAGCACCTGGACAACGTGAAGGTGTTCGAGAA<br>GGCCATCAGCAAGCTGAGCAAGAACGCCATCGACGACCTGGA<br>TGCCACATATTCTGGCCTGTGCGGCACAAATCTGTACGACGTG<br>TTCACCGTGGACTACTTCAACTTCCTGCTGCCCCAAAGCGGAA<br>TCACCGAGTACAACAAGATCATCGGCGGCTACACAACAAGCG<br>ACGGCACCAAAGTGAAGGGCATCAACGAGTACATCAACCTGT<br>ACAACCAGCAGGTGAGCAAGAGAGACAAGATCCCCAACCTGA<br>AGATCCTGTACAAGCAGATCCTGAGCGAGAGCGAGAAGGTGT |

| | |
|---|---|
| | CTTTCATCCCCCCCAAGTTCGAGGACGACAACGAACTGCTGTC TGCCGTGAGCGAGTTCTATGCCAACGACGAGACATTTGATGGC ATGCCCCTGAAGAAAGCCATCGACGAAACCAAACTGCTGTTC GGCAACCTGGACAACAGCAGCCTGAACGGCATCTACATCCAG AACGACAGAAGCGTGACCAACCTGAGCAACAGCATGTTCGGC AGCTGGAGCGTGATTGAGGACCTGTGGAACAAGAACTACGAC AGCGTGAACAGCAACAGCAGAATCAAGGACATCCAGAAGAG AGAGGACAAGAGAAAGAAGGCCTACAAGGCCGAGAAGAAGC TGAGCCTGAGCTTCCTGCAGGTGCTGATCAGCAACAGCGAGA ACGACGAGATCAGAAAGAAGAGCATCGTGGACTACTACAAGA CCAGCCTGATGCAGCTGACCGACAACCTGAGCGACAAGTACA AAGAAGCCGCCCCCCTGTTTTCTGAGAACTACGACAACGAGA AGGGCCTGAAGAACGACGACAAGAGCATCAGCCTGATCAAGA ACTTCCTGGACGCCATCAAGGAGATCGAGAAGTTCATCAAGCC CCTGAGCGAGACAAATATCACCGGCGAGAAGAACGACCTGTT CTACAGCCAGTTCACCCCCCTGCTGGACAACATCAGCAGAATC GACAGACTGTACGACAAGGTGAGAAACTACGTGACCCAGAAG CCCTTCAGCACCGACAAGATCAAGCTGAACTTCGGCAACAGC CAGCTTCTGAACGGCTGGGACAGAAACAAGGAGAAGGACTGT GGCGCTGTGCTGCTGTGTAAGGACGAGAAGTACTACCTGGCC ATCATCGACAAGAGCAACAACAGCATCCTGGAGAACATCGAC TTCCAGGACTGCAACGAGAGCGACTACTACGAGAAGATCGTG TACAAGCTGCTGACCAAGATCTCTGGCAACCTGCCCAGAGTGT TCTTCAGCGAGAAGCACAAGAAGCTGCTGAGCCCCAGCGATG AGATCCTGAAGATCTACAAGAGCGGCACCTTCAAGAAGGGCG ACAAGTTCAGCCTTGACGACTGCCACAAGCTGATCGACTTCTA CAAGGAGAGCTTCAAGAAGTACCCCAAGTGGCTGATCTACAA CTTCAAGTTCAAGAACACCAACGAGTACAACGACATCAGCGA GTTCTACAACGACGTGGCCAGCCAGGGATACAACATCAGCAA GATGAAGATCCCCACCAGCTTCATCGACAAGCTGGTGGACGA GGGCAAGATCTACCTGTTCCAGCTGTACAACAAGGACTTCAGC CCCCACAGCAAGGGAACACCTAACCTGCACACCCTGTACTTCA AGATGCTGTTCGACGAGAGAAACCTGGAGGACGTGGTGTACA AGCTGAATGGCGAGGCCGAGATGTTTTACAGACCCGCCAGCA TCAAGTATGACAAGCCCACCCACCCTAAGAACACCCCCATCAA GAACAAGAACACCCTGAACGACAAGAAGGCCAGCACCTTCCC CTACGACCTGATCAAGGACAAGAGATACACCAAGTGGCAGTT CAGCCTGCACTTCCCCATCACCATGAACTTCAAGGCCCCCGAC AGAGCCATGATCAACGACGACGTGAGAAACCTGCTGAAGAGC TGCAACAACAACTTCATCATCGGCATCGACAGAGGCGAGAGA AACCTGCTGTACGTGAGCGTGATCGATAGCAACGGCGCCATC ATCTACCAGCACAGCCTGAACATCATCGGCAACAAGTTCAAGG GCAAGACCTACGAAACCAACTACAGAGAGAAGCTGGCCACCA GAGAGAAGGAGAGAACCGAGCAGAGAAGAAACTGGAAGGCC |

| | |
|---|---|
| | ATCGAGAGCATCAAGGAGCTGAAGGAGGGCTACATCAGCCAA<br>ACCGTGCACGTGATTTGCCAGCTGGTGGTGAAGTACGACGCC<br>ATCATCGTGATGGAGAAGCTGACCGACGGCTTCAAGAGAGGC<br>AGAACCAAGTTCGAGAAGCAGGTGTACCAGAAGTTCGAGAAG<br>ATGCTGATCGACAAGCTGAACTACTACGTGGACAAGAAGCTG<br>GACCCCAATGAGGAAGGCGGACTGCTGCATGCTTATCAGCTG<br>ACCAACAAGCTGGACAGCTTCGACAAGCTGGGAATGCAGAGC<br>GGCTTCATCTTCTACGTCAGACCCGACTTCACCAGCAAAATCG<br>ACCCCGTGACCGGATTTGTGAACCTGCTGTACCCCAGATACGA<br>GAACATCGACAAGGCCAAGGACATGATCAGCAGATTCGACGA<br>CATCAGATACAACGCCGGCGAGGACTTCTTCGAGTTCGACATC<br>GACTACGACAAGTTCCCCAAGACCGCCAGCGACTACAGAAAG<br>AAGTGGACCATCTGCACCAACGGCGAGAGAATCGAGGCCTTC<br>AGAAACCCCGCCAACAACAACGAGTGGAGCTACAGAACCATC<br>ATCCTGGCCGAGAAGTTCAAGGAGCTGTTCGACAACAACAGC<br>ATCAACTACAGAGACAGCGACGACCTGAAAGCCGAGATCCTG<br>AGCCAAACCAAGGGCAAGTTCTTCGAGGACTTCTTCAAGCTGC<br>TGAGACTGACCCTGCAGATGAGAAACAGCAACCCCGAAACCG<br>GAGAGGACAGGATTCTGAGCCCCGTGAAGGACAAGAACGGC<br>AACTTCTACGACAGCAGCAAGTACGACGAGAAGAGCAAGCTG<br>CCCTGTGACGCTGATGCTAACGGCGCTTACAACATCGCCAGAA<br>AGGGCCTGTGGATCGTGGAGCAGTTCAAGAAGGCCGACAACG<br>TGTCTGCTGTGGAACCCGTGATCCACAACGACAAGTGGCTGAA<br>GTTCGTGCAGGAGAACGACATGGCCAACAACaaaaggccggcgg<br>ccacgaaaaaggccggccaggcaaaaaagaaaaaggaattcggcagtggaga<br>gggcagaggaagtctgctaacatgcggtgacgtcgaggagaatcctggcccagt<br>gagcaagggcgaggagctgttcaccggggtggtgcccatcctggtcgagctgga<br>cggcgacgtaaacggccacaagttcagcgtgtccggcgagggcgagggcgatg<br>ccacctacggcaagctgaccctgaagttcatctgcaccaccggcaagctgcccgtg<br>ccctggcccaccctcgtgaccaccctgacctacggcgtgcagtgcttcagccgctac<br>cccgaccacatgaagcagcacgacttcttcaagtccgccatgcccgaaggctacgt<br>ccaggagcgcaccatcttcttcaaggacgacggcaactacaagacccgcgccgag<br>gtgaagttcgagggcgacaccctggtgaaccgcatcgagctgaagggcatcgact<br>tcaaggaggacggcaacatcctggggcacaagctggagtacaactacaacagcc<br>acaacgtctatatcatggccgacaagcagaagaacggcatcaaggtgaacttcaa<br>gatccgccacaacatcgaggacggcagcgtgcagctcgccgaccactaccagca<br>gaacacccccatcggcgacggccccgtgctgctgcccgacaaccactacctgagc<br>acccagtccgccctgagcaaagaccccaacgagaagcgcgatcacatggtcctgc<br>tggagttcgtgaccgccgccgggatcactctcggcatggacgagctgtacaagga<br>attctaactagagctcgctgatcagcctcgactgtgccttctagttgccagccatctgtt<br>gtttgcccctcccccgtgccttccttgaccctggaaggtgccactcccactgtcctttcc<br>taataaaatgaggaaattgcatcgcattgtctgagtaggtgtcattctattctggggg<br>gtggggtggggcaggacagcaagggggaggattgggaagagaatagcaggc<br>atgctggggagcggccgcaggaacccctagtgatggagttggccactccctctctg |

| | |
|---|---|
| | cgcgctcgctcgctcactgaggccgggcgaccaaaggtcgcccgacgcccgggc<br>tttgcccgggcggcctcagtgagcgagcgagcgcgcagctgcctgcaggggcgc<br>ctgatgcggtattttctccttacgcatctgtgcggtatttcacaccgcatacgtcaaagc<br>aaccatagtacgcgccctgtagcggcgcattaagcgcggcgggtgtggtggttac<br>gcgcagcgtgaccgctacacttgccagcgccttagcgcccgctcctttcgctttcttcc<br>cttcctttctcgccacgttcgccggctttccccgtcaagctctaaatcgggggctccctt<br>tagggttccgatttagtgctttacggcacctcgaccccaaaaaacttgatttgggtga<br>tggttcacgtagtgggccatcgccctgatagacggtttttcgccctttgacgttggag<br>tccacgttctttaatagtggactcttgttccaaactggaacaacactcaactctatctcg<br>ggctattcttttgatttataagggattttgccgatttcggtctattggttaaaaaatgag<br>ctgatttaacaaaaatttaacgcgaattttaacaaaatattaacgtttacaattttatggt<br>gcactctcagtacaatctgctctgatgccgcatagttaagccagccccgacacccgc<br>caacacccgctgacgcgccctgacgggcttgtctgctcccggcatccgcttacagac<br>aagctgtgaccgtctccgggagctgcatgtgtcagaggttttcaccgtcatcaccga<br>aacgcgcgagacgaaagggcctcgtgatacgcctatttttataggttaatgtcatga<br>taataatggtttcttagacgtcaggtggcacttttcggggaaatgtgcgcggaaccc<br>ctatttgtttatttttctaaatacattcaaatatgtatccgctcatgagacaataaccctga<br>taaatgcttcaataatattgaaaaaggaagagtatgagtattcaacatttccgtgtcg<br>cccttattcccttttttgcggcattttgccttcctgtttttgctcacccagaaacgctggtg<br>aaagtaaaagatgctgaagatcagttgggtgcacgagtgggttacatcgaactgg<br>atctcaacagcggtaagatccttgagagttttcgccccgaagaacgttttccaatgat<br>gagcacttttaaagttctgctatgtggcgcggtattatcccgtattgacgccgggcaa<br>gagcaactcggtcgccgcatacactattctcagaatgacttggttgagtactcacca<br>gtcacagaaaagcatcttacggatggcatgacagtaagagaattatgcagtgctgc<br>cataaccatgagtgataacactgcggccaacttacttctgacaacgatcggaggacc<br>gaaggagctaaccgcttttttgcacaacatgggggatcatgtaactcgccttgatcgt<br>tgggaaccggagctgaatgaagccataccaaacgacgagcgtgacaccacgatg<br>cctgtagcaatggcaacaacgttgcgcaaactattaactggcgaactacttactctag<br>cttcccggcaacaattaatagactggatggaggcggataaagttgcaggaccactt<br>ctgcgctcggcccttccggctggctggtttattgctgataaatctggagccggtgag<br>cgtggaagccgcggtatcattgcagcactggggccagatggtaagccctcccgtat<br>cgtagttatctacacgacggggagtcaggcaactatggatgaacgaaatagacag<br>atcgctgagataggtgcctcactgattaagcattggtaactgtcagaccaagtttact<br>catatatactttagattgatttaaaacttcatttttaatttaaaaggatctaggtgaagat<br>cctttttgataatctcatgaccaaaatcccttaacgtgagttttcgttccactgagcgtca<br>gaccccgtagaaaagatcaaaggatcttcttgagatcctttttttctgcgcgtaatctg<br>ctgcttgcaaacaaaaaaaccaccgctaccagcggtggtttgtttgccggatcaaga<br>gctaccaactctttttccgaaggtaactggcttcagcagagcgcagataccaaatact<br>gttcttctagtgtagccgtagttaggccaccacttcaagaactctgtagcaccgcctac<br>atacctcgctctgctaatcctgttaccagtggctgctgccagtggcgataagtcgtgt<br>cttaccgggttggactcaagacgatagttaccggataaggcgcagcggtcgggct<br>gaacggggggttcgtgcacacagcccagcttggagcgaacgacctacaccgaac<br>tgagatacctacagcgtgagctatgagaaagcgccacgcttcccgaagggagaaa<br>ggcggacaggtatccggtaagcggcagggtcggaacaggagagcgcacgagg |

| | |
|---|---|
| | gagcttccagggggaaacgcctggtatctttatagtcctgtcgggtttcgccacctct<br>gacttgagcgtcgatttttgtgatgctcgtcaggggggcggagcctatggaaaaac<br>gccagcaacgcggcctttttacggttcctggccttttgctggccttttgctcacatgt |
| **Eukaryotic expression vector of BvCas12a with HDV self-cleaving ribozyme added at the 3' end of the crRNA** | |
| Plasmid-<br>eukaryotes-<br>BvCas12a-<br>HDV-NC | gagggcctatttcccatgattccttcatatttgcatatacgatacaaggctgttagaga<br>gataattggaattaatttgactgtaaacacaaagatattagtacaaaatacgtgacgt<br>agaaagtaataatttcttgggtagtttgcagttttaaaattatgttttaaaatggactatc<br>atatgcttaccgtaacttgaaagtatttcgatttcttggctttatatatcttgtggaaagg<br>acgaaacaccgAATTTCTACTATTGTAGATgggtcttcgagaagacctgg<br>ccggcatggtcccagcctcctcgctggcgccggctgggcaacatgcttcggcatgg<br>cgaatgggactttttttgttttagagctagaaatagcaagttaaaataaggctagtccg<br>tttttagcgcgtgcgccaattctgcagacaaatggctctagaggtacccgttacataa<br>cttacggtaaatggcccgcctggctgaccgcccaacgacccccgcccattgacgtc<br>aatagtaacgccaatagggactttccattgacgtcaatgggtggagtatttacggta<br>aactgcccacttggcagtacatcaagtgtatcatatgccaagtacgccccctattgac<br>gtcaatgacggtaaatggcccgcctggcattgtgcccagtacatgaccttatgggac<br>tttcctacttggcagtacatctacgtattagtcatcgctattaccatggtcgaggtgag<br>ccccacgttctgcttcactctccccatctcccccccctccccacccccaattttgtatttatt<br>tattttttaattattttgtgcagcgatgggggcgggggggggggggggcgcgcg<br>ccaggcggggcggggcggggcgaggggcggggcggggcgaggcggagag<br>gtgcggcggcagccaatcagagcggcgcgctccgaaagtttccttttatggcgag<br>gcggcggcggcggcggccctataaaaagcgaagcgcgcggcgggcgggagtc<br>gctgcgcgctgccttcgccccgtgccccgctccgccgccgcctcgcgccgcccgcc<br>ccggctctgactgaccgcgttactcccacaggtgagcgggcgggacggcccttctc<br>ctccgggctgtaattagctgagcaagaggtaagggtttaagggatggttggttggt<br>ggggtattaatgtttaattacctggagcacctgcctgaaatcactttttttcaggttgga<br>ccggtgccaccatggactataaggaccacgacggagactacaaggatcatgatatt<br>gattacaaagacgatgacgataagatggccccaaagaagaagcggaaggtcggt<br>atccacggagtcccagcagccATGCAGGAGAGAAAGAAGATCAGCC<br>ACCTGACCCACAGAAACAGCGTGAAGAAAACCATCAGAATGC<br>AGCTGAACCCCGTGGGAAAGACCATGGACTACTTCCAGGCCA<br>AGCAGATCCTGGAGAACGACGAGAAGCTGAAGGAGGACTAC<br>CAGAAGATCAAGGAGATCGCCGACAGATTCTACAGAAACCTG<br>AACGAGGACGTGCTGAGCAAAACCGGACTGGACAAGCTGAA<br>GGACTACGCCGAGATCTACTACCATTGCAACACCGACGCCGA<br>CAGAAAGAGACTGAACGAGTGCGCCAGCGAGCTGAGAAAGG<br>AGATCGTGAAGAACTTCAAGAACAGAGATGAGTACAACAAGC<br>TGTTCAACAAGAAGATGATCGAGATCGTGCTGCCCAAGCACCT<br>GAAGAACGAGGACGAGAAGGAAGTGGTGGCCAGCTTCAAGA<br>ACTTCACCACCTACTTCACCGGCTTCTTCACCAACAGAAAGAA<br>CATGTACAGCGACGGCGAAGAGTCTACCGCTATTGCCTACAG<br>ATGCATCAACGAGAACCTGCCCAAGCACCTGGACAACGTGAA<br>GGTGTTCGAGAAGGCCATCAGCAAGCTGAGCAAGAACGCCAT |

| | |
|---|---|
| | CGACGACCTGGATGCCACATATTCTGGCCTGTGCGGCACAAAT<br>CTGTACGACGTGTTCACCGTGGACTACTTCAACTTCCTGCTGCC<br>CCAAAGCGGAATCACCGAGTACAACAAGATCATCGGCGGCTA<br>CACAACAAGCGACGGCACCAAAGTGAAGGGCATCAACGAGTA<br>CATCAACCTGTACAACCAGCAGGTGAGCAAGAGAGACAAGAT<br>CCCCAACCTGAAGATCCTGTACAAGCAGATCCTGAGCGAGAG<br>CGAGAAGGTGTCTTTCATCCCCCCCAAGTTCGAGGACGACAAC<br>GAACTGCTGTCTGCCGTGAGCGAGTTCTATGCCAACGACGAGA<br>CATTTGATGGCATGCCCCTGAAGAAAGCCATCGACGAAACCA<br>AACTGCTGTTCGGCAACCTGGACAACAGCAGCCTGAACGGCA<br>TCTACATCCAGAACGACAGAAGCGTGACCAACCTGAGCAACA<br>GCATGTTCGGCAGCTGGAGCGTGATTGAGGACCTGTGGAACA<br>AGAACTACGACAGCGTGAACAGCAACAGCAGAATCAAGGACA<br>TCCAGAAGAGAGAGGACAAGAGAAAGAAGGCCTACAAGGCC<br>GAGAAGAAGCTGAGCCTGAGCTTCCTGCAGGTGCTGATCAGC<br>AACAGCGAGAACGACGAGATCAGAAAGAAGAGCATCGTGGA<br>CTACTACAAGACCAGCCTGATGCAGCTGACCGACAACCTGAG<br>CGACAAGTACAAAGAAGCCGCCCCCCTGTTTTCTGAGAACTAC<br>GACAACGAGAAGGGCCTGAAGAACGACGACAAGAGCATCAG<br>CCTGATCAAGAACTTCCTGGACGCCATCAAGGAGATCGAGAA<br>GTTCATCAAGCCCCTGAGCGAGACAAATATCACCGGCGAGAA<br>GAACGACCTGTTCTACAGCCAGTTCACCCCCCTGCTGGACAAC<br>ATCAGCAGAATCGACAGACTGTACGACAAGGTGAGAAACTAC<br>GTGACCCAGAAGCCCTTCAGCACCGACAAGATCAAGCTGAAC<br>TTCGGCAACAGCCAGCTTCTGAACGGCTGGGACAGAAACAAG<br>GAGAAGGACTGTGGCGCTGTGCTGCTGTGTAAGGACGAGAAG<br>TACTACCTGGCCATCATCGACAAGAGCAACAACAGCATCCTGG<br>AGAACATCGACTTCCAGGACTGCAACGAGAGCGACTACTACG<br>AGAAGATCGTGTACAAGCTGCTGACCAAGATCTCTGGCAACCT<br>GCCCAGAGTGTTCTTCAGCGAGAAGCACAAGAAGCTGCTGAG<br>CCCCAGCGATGAGATCCTGAAGATCTACAAGAGCGGCACCTT<br>CAAGAAGGGCGACAAGTTCAGCCTTGACGACTGCCACAAGCT<br>GATCGACTTCTACAAGGAGAGCTTCAAGAAGTACCCCAAGTG<br>GCTGATCTACAACTTCAAGTTCAAGAACACCAACGAGTACAAC<br>GACATCAGCGAGTTCTACAACGACGTGGCCAGCCAGGGATAC<br>AACATCAGCAAGATGAAGATCCCCACCAGCTTCATCGACAAG<br>CTGGTGGACGAGGGCAAGATCTACCTGTTCCAGCTGTACAACA<br>AGGACTTCAGCCCCCACAGCAAGGGAACACCTAACCTGCACA<br>CCCTGTACTTCAAGATGCTGTTCGACGAGAGAAACCTGGAGGA<br>CGTGGTGTACAAGCTGAATGGCGAGGCCGAGATGTTTTACAG<br>ACCCGCCAGCATCAAGTATGACAAGCCCACCCACCCTAAGAA<br>CACCCCCATCAAGAACAAGAACACCCTGAACGACAAGAAGGC<br>CAGCACCTTCCCCTACGACCTGATCAAGGACAAGAGATACAC<br>CAAGTGGCAGTTCAGCCTGCACTTCCCCATCACCATGAACTTC |

| | |
|---|---|
| | AAGGCCCCCGACAGAGCCATGATCAACGACGACGTGAGAAAC<br>CTGCTGAAGAGCTGCAACAACAACTTCATCATCGGCATCGACA<br>GAGGCGAGAGAAACCTGCTGTACGTGAGCGTGATCGATAGCA<br>ACGGCGCCATCATCTACCAGCACAGCCTGAACATCATCGGCA<br>ACAAGTTCAAGGGCAAGACCTACGAAACCAACTACAGAGAGA<br>AGCTGGCCACCAGAGAGAAGGAGAGAACCGAGCAGAGAAGA<br>AACTGGAAGGCCATCGAGAGCATCAAGGAGCTGAAGGAGGG<br>CTACATCAGCCAAACCGTGCACGTGATTTGCCAGCTGGTGGTG<br>AAGTACGACGCCATCATCGTGATGGAGAAGCTGACCGACGGC<br>TTCAAGAGAGGCAGAACCAAGTTCGAGAAGCAGGTGTACCAG<br>AAGTTCGAGAAGATGCTGATCGACAAGCTGAACTACTACGTGG<br>ACAAGAAGCTGGACCCCAATGAGGAAGGCGGACTGCTGCATG<br>CTTATCAGCTGACCAACAAGCTGGACAGCTTCGACAAGCTGG<br>GAATGCAGAGCGGCTTCATCTTCTACGTCAGACCCGACTTCAC<br>CAGCAAAATCGACCCCGTGACCGGATTTGTGAACCTGCTGTAC<br>CCCAGATACGAGAACATCGACAAGGCCAAGGACATGATCAGC<br>AGATTCGACGACATCAGATACAACGCCGGCGAGGACTTCTTC<br>GAGTTCGACATCGACTACGACAAGTTCCCCAAGACCGCCAGC<br>GACTACAGAAAGAAGTGGACCATCTGCACCAACGGCGAGAGA<br>ATCGAGGCCTTCAGAAACCCCGCCAACAACAACGAGTGGAGC<br>TACAGAACCATCATCCTGGCCGAGAAGTTCAAGGAGCTGTTCG<br>ACAACAACAGCATCAACTACAGAGACAGCGACGACCTGAAAG<br>CCGAGATCCTGAGCCAAACCAAGGGCAAGTTCTTCGAGGACT<br>TCTTCAAGCTGCTGAGACTGACCCTGCAGATGAGAAACAGCAA<br>CCCCGAAACCGGAGAGGACAGGATTCTGAGCCCCGTGAAGGA<br>CAAGAACGGCAACTTCTACGACAGCAGCAAGTACGACGAGAA<br>GAGCAAGCTGCCCTGTGACGCTGATGCTAACGGCGCTTACAA<br>CATCGCCAGAAAGGGCCTGTGGATCGTGGAGCAGTTCAAGAA<br>GGCCGACAACGTGTCTGCTGTGGAACCCGTGATCCACAACGA<br>CAAGTGGCTGAAGTTCGTGCAGGAGAACGACATGGCCAACAA<br>Caaaaggccggcggccacgaaaaaggccggccaggcaaaaaagaaaaagga<br>attcggcagtggagagggcagaggaagtctgctaacatgcggtgacgtcgagga<br>gaatcctggcccagtgagcaagggcgaggagctgttcaccggggtggtgcccat<br>cctggtcgagctggacggcgacgtaaacggccacaagttcagcgtgtccggcga<br>gggcgagggcgatgccacctacggcaagctgaccctgaagttcatctgcaccacc<br>ggcaagctgcccgtgccctggcccaccctcgtgaccaccctgacctacggcgtgca<br>gtgcttcagccgctaccccgaccacatgaagcagcacgacttcttcaagtccgccat<br>gcccgaaggctacgtccaggagcgcaccatcttcttcaaggacgacggcaactac<br>aagacccgcgccgaggtgaagttcgagggcgacaccctggtgaaccgcatcgag<br>ctgaagggcatcgacttcaaggaggacggcaacatcctggggcacaagctggag<br>tacaactacaacagccacaacgtctatatcatggccgacaagcagaagaacggcat<br>caaggtgaacttcaagatccgccacaacatcgaggacggcagcgtgcagctcgcc<br>gaccactaccagcagaacacccccatcggcgacggccccgtgctgctgcccgaca<br>accactacctgagcacccagtccgccctgagcaaagaccccaacgagaagcgcga |

| | |
|---|---|
| | tcacatggtcctgctggagttcgtgaccgccgccgggatcactctcggcatggacg<br>agctgtacaaggaattctaactagagctcgctgatcagcctcgactgtgccttctagt<br>tgccagccatctgttgtttgcccctcccccgtgccttccttgaccctggaaggtgccac<br>tcccactgtcctttcctaataaaatgaggaaattgcatcgcattgtctgagtaggtgtc<br>attctattctggggggtggggtggggcaggacagcaagggggaggattgggaa<br>gagaatagcaggcatgctggggagcggccgcaggaacccctagtgatggagtt<br>ggccactccctctctgcgcgctcgctcgctcactgaggccgggcgaccaaaggtcg<br>cccgacgcccgggctttgcccgggcggcctcagtgagcgagcgagcgcgcagct<br>gcctgcaggggcgcctgatgcggtattttctccttacgcatctgtgcggtatttcacac<br>cgcatacgtcaaagcaaccatagtacgcgccctgtagcggcgcattaagcgcggc<br>gggtgtggtggttacgcgcagcgtgaccgctacacttgccagcgccttagcgcccg<br>ctcctttcgctttcttcccttcctttctcgccacgttcgccggctttccccgtcaagctctaa<br>atcgggggctccctttagggttccgatttagtgctttacggcacctcgaccccaaaaa<br>acttgatttgggtgatggttcacgtagtgggccatcgccctgatagacggtttttcgc<br>cctttgacgttggagtccacgttctttaatagtggactcttgttccaaactggaacaac<br>actcaactctatctcgggctattcttttgatttataagggattttgccgatttcggtctatt<br>ggttaaaaaatgagctgatttaacaaaaatttaacgcgaattttaacaaaatattaac<br>gtttacaattttatggtgcactctcagtacaatctgctctgatgccgcatagttaagcca<br>gccccgacacccgccaacacccgctgacgcgccctgacgggcttgtctgctcccgg<br>catccgcttacagacaagctgtgaccgtctccgggagctgcatgtgtcagaggtttt<br>caccgtcatcaccgaaacgcgcgagacgaaagggcctcgtgatacgcctatttttat<br>aggttaatgtcatgataataatggtttcttagacgtcaggtggcacttttcggggaaa<br>tgtgcgcggaacccctatttgtttatttttctaaatacattcaaatatgtatccgctcatg<br>agacaataaccctgataaatgcttcaataatattgaaaaaggaagagtatgagtatt<br>caacatttccgtgtcgcccttattcccttttttgcggcattttgccttcctgtttttgctcacc<br>cagaaacgctggtgaaagtaaaagatgctgaagatcagttgggtgcacgagtgg<br>gttacatcgaactggatctcaacagcggtaagatccttgagagttttcgccccgaag<br>aacgttttccaatgatgagcacttttaaagttctgctatgtggcgcggtattatcccgta<br>ttgacgccgggcaagagcaactcggtcgccgcatacactattctcagaatgacttg<br>gttgagtactcaccagtcacagaaaagcatcttacggatggcatgacagtaagaga<br>attatgcagtgctgccataaccatgagtgataacactgcggccaacttacttctgaca<br>acgatcggaggaccgaaggagctaaccgcttttttgcacaacatgggggatcatgt<br>aactcgccttgatcgttgggaaccggagctgaatgaagccataccaaacgacgag<br>cgtgacaccacgatgcctgtagcaatggcaacaacgttgcgcaaactattaactgg<br>cgaactacttactctagcttcccggcaacaattaatagactggatggaggcggataa<br>agttgcaggaccacttctgcgctcggcccttccggctggctggtttattgctgataaa<br>tctggagccggtgagcgtggaagccgcggtatcattgcagcactggggccagat<br>ggtaagccctcccgtatcgtagttatctacacgacggggagtcaggcaactatgga<br>tgaacgaaatagacagatcgctgagataggtgcctcactgattaagcattggtaact<br>gtcagaccaagtttactcatatatactttagattgatttaaaacttcatttttaatttaaaa<br>ggatctaggtgaagatcctttttgataatctcatgaccaaaatcccttaacgtgagtttt<br>cgttccactgagcgtcagaccccgtagaaaagatcaaaggatcttcttgagatccttt<br>ttttctgcgcgtaatctgctgcttgcaaacaaaaaaaccaccgctaccagcggtggtt<br>tgtttgccggatcaagagctaccaactctttttccgaaggtaactggcttcagcagag |

| | |
|---|---|
| | cgcagataccaaatactgttcttctagtgtagccgtagttaggccaccacttcaagaa<br>ctctgtagcaccgcctacatacctcgctctgctaatcctgttaccagtggctgctgcca<br>gtggcgataagtcgtgtcttaccgggttggactcaagacgatagttaccggataag<br>gcgcagcggtcgggctgaacggggggttcgtgcacacagcccagcttggagcg<br>aacgacctacaccgaactgagatacctacagcgtgagctatgagaaagcgccacg<br>cttcccgaagggagaaaggcggacaggtatccggtaagcggcagggtcggaac<br>aggagagcgcacgagggagcttccagggggaaacgcctggtatctttatagtcct<br>gtcgggtttcgccacctctgacttgagcgtcgatttttgtgatgctcgtcaggggggc<br>ggagcctatggaaaaacgccagcaacgcggcctttttacggttcctggccttttgct<br>ggccttttgctcacatgt |
| **Eukaryote Expression Vector of AsCas12a** | |
| Plasmid-eukaryotes-AsCas12a-NC | gagggcctatttcccatgattccttcatatttgcatatacgatacaaggctgttagaga<br>gataattggaattaatttgactgtaaacacaaagatattagtacaaaatacgtgacgt<br>agaaagtaataatttcttgggtagtttgcagttttaaaattatgttttaaaatggactatc<br>atatgcttaccgtaacttgaaagtatttcgatttcttggctttatatatcttgtggaaagg<br>acgaaacaccgaatttctactcttgtagatgggtcttcgagaagaccttttttttgtttta<br>gagctagaaatagcaagttaaaataaggctagtccgtttttagcgcgtgcgccaatt<br>ctgcagacaaatggctctagaggtacccgttacataacttacggtaaatggcccgcc<br>tggctgaccgcccaacgacccccgcccattgacgtcaatagtaacgccaataggga<br>ctttccattgacgtcaatgggtggagtatttacggtaaactgcccacttggcagtaca<br>tcaagtgtatcatatgccaagtacgccccctattgacgtcaatgacggtaaatggccc<br>gcctggcattgtgcccagtacatgaccttatgggactttcctacttggcagtacatcta<br>cgtattagtcatcgctattaccatggtcgaggtgagccccacgttctgcttcactctcc<br>ccatctcccccccctccccacccccaattttgtatttatttattttttaattattttgtgcagc<br>gatgggggcggggggggggggggggcgcgcgccaggcggggcggggcgg<br>ggcgaggggcggggcggggcgaggcggagaggtgcggcggcagccaatca<br>gagcggcgcgctccgaaagtttccttttatggcgaggcggcggcggcggcggcc<br>ctataaaaagcgaagcgcgcggcgggcgggagtcgctgcgcgctgccttcgccc<br>cgtgccccgctccgccgccgcctcgcgccgcccgccccggctctgactgaccgcgt<br>tactcccacaggtgagcgggcgggacggcccttctcctccgggctgtaattagctg<br>agcaagaggtaagggtttaagggatggttggttggtggggtattaatgtttaattac<br>ctggagcacctgcctgaaatcactttttttcaggttggaccggtgccaccatggacta<br>taaggaccacgacggagactacaaggatcatgatattgattacaaagacgatgac<br>gataagatggccccaaagaagaagcggaaggtcggtatccacggagtcccagca<br>gccatgacacagttcgagggctttaccaacctgtatcaggtgagcaagacactgcg<br>gtttgagctgatcccacagggcaagaccctgaagcacatccaggagcagggcttc<br>atcgaggaggacaaggcccgcaatgatcactacaaggagctgaagcccatcatcg<br>atcggatctacaagacctatgccgaccagtgcctgcagctggtgcagctggattgg<br>gagaacctgagcgccgccatcgactcctatagaaaggagaaaaccgaggagaca<br>aggaacgccctgatcgaggagcaggccacatatcgcaatgccatccacgactactt<br>catcggccggacagacaacctgaccgatgccatcaataagagacacgccgagatc<br>tacaagggcctgttcaaggccgagctgtttaatggcaaggtgctgaagcagctgg<br>gcaccgtgaccacaaccgagcacgagaacgccctgctgcggagcttcgacaagtt<br>tacaacctacttctccggcttttatgagaacaggaagaacgtgttcagcgccgagga |

| | |
|---|---|
| | tatcagcacagccatcccacaccgcatcgtgcaggacaacttccccaagtttaagga<br>gaattgtcacatcttcacacgcctgatcaccgccgtgcccagcctgcgggagcacttt<br>gagaacgtgaagaaggccatcggcatcttcgtgagcacctccatcgaggaggtgt<br>tttccttccctttttataaccagctgctgacacagacccagatcgacctgtataaccagc<br>tgctgggaggaatctctcgggaggcaggcaccgagaagatcaagggcctgaac<br>gaggtgctgaatctggccatccagaagaatgatgagacagcccacatcatcgcctc<br>cctgccacacagattcatcccccctgtttaagcagatcctgtccgataggaacaccctg<br>tctttcatcctggaggagtttaagagcgacgaggaagtgatccagtccttctgcaag<br>tacaagacactgctgagaaacgagaacgtgctggagacagccgaggccctgttta<br>acgagctgaacagcatcgacctgacacacatcttcatcagccacaagaagctggag<br>acaatcagcagcgccctgtgcgaccactgggatacactgaggaatgccctgtatga<br>gcggagaatctccgagctgacaggcaagatcaccaagtctgccaaggagaaggt<br>gcagcgcagcctgaagcacgaggatatcaacctgcaggagatcatctctgccgca<br>ggcaaggagctgagcgaggccttcaagcagaaaaccagcgagatcctgtcccac<br>gcacacgccgccctggatcagccactgcctacaaccctgaagaagcaggaggag<br>aaggagatcctgaagtctcagctggacagcctgctgggcctgtaccacctgctgga<br>ctggtttgccgtggatgagtccaacgaggtggaccccgagttctctgcccggctga<br>ccggcatcaagctggagatggagccttctctgagcttctacaacaaggccagaaat<br>tatgccaccaagaagccctactccgtggagaagttcaagctgaactttcagatgcct<br>acactggcctctggctgggacgtgaataaggagaagaacaatggcgccatcctgt<br>ttgtgaagaacggcctgtactatctgggcatcatgccaaagcagaagggcaggtat<br>aaggccctgagcttcgagcccacagagaaaaccagcgagggctttgataagatgt<br>actatgactacttccctgatgccgccaagatgatcccaaagtgcagcacccagctga<br>aggccgtgacagcccactttcagacccacacaacccccatcctgctgtccaacaattt<br>catcgagcctctggagatcacaaaggagatctacgacctgaacaatcctgagaag<br>gagccaaagaagtttcagacagcctacgccaagaaaaccggcgaccagaaggg<br>ctacagagaggccctgtgcaagtggatcgacttcacaagggattttctgtccaagta<br>taccaagacaacctctatcgatctgtctagcctgcggccatcctctcagtataaggac<br>ctgggcgagtactatgccgagctgaatcccctgctgtaccacatcagcttccagaga<br>atcgccgagaaggagatcatggatgccgtggagacaggcaagctgtacctgttcc<br>agatctataacaaggactttgccaagggccaccacggcaagcctaatctgcacaca<br>ctgtattggaccggcctgttttctccagagaacctggccaagacaagcatcaagctg<br>aatggccaggccgagctgttctaccgccctaagtccaggatgaagaggatggcac<br>accggctgggagagaagatgctgaacaagaagctgaaggatcagaaaacccca<br>atccccgacaccctgtaccaggagctgtacgactatgtgaatcacagactgtcccac<br>gacctgtctgatgaggccagggccctgctgcccaacgtgatcaccaaggaggtgt<br>ctcacgagatcatcaaggataggcgctttaccagcgacaagttctttttccacgtgcc<br>tatcacactgaactatcaggccgccaattccccatctaagttcaaccagagggtgaat<br>gcctacctgaaggagcaccccgagacacctatcatcggcatcgatcggggcgaga<br>gaaacctgatctatatcacagtgatcgactccaccggcaagatcctggagcagcgg<br>agcctgaacaccatccagcagtttgattaccagaagaagctggacaacagggaga<br>aggagagggtggcagcaaggcaggcctggtctgtggtgggcacaatcaaggat<br>ctgaagcagggctatctgagccaggtcatccacgagatcgtggacctgatgatcca<br>ctaccaggccgtggtggtgctggagaacctgaatttcggctttaagagcaagagg |

| | |
|---|---|
| | accggcatcgccgagaaggccgtgtaccagcagttcgagaagatgctgatcgata<br>agctgaattgcctggtgctgaaggactatccagcagagaaagtgggaggcgtgct<br>gaacccataccagctgacagaccagttcacctcctttgccaagatgggcacccagtc<br>tggcttcctgttttacgtgcctgccccatatacatctaagatcgatcccctgaccggctt<br>cgtggaccccttcgtgtggaaaaccatcaagaatcacgagagccgcaagcacttcc<br>tggagggcttcgactttctgcactacgacgtgaaaaccggcgacttcatcctgcactt<br>taagatgaacagaaatctgtccttccagaggggcctgcccggctttatgcctgcatg<br>ggatatcgtgttcgagaagaacgagacacagtttgacgccaagggcaccccctttca<br>tcgccggcaagagaatcgtgccagtgatcgagaatcacagattcaccggcagata<br>ccgggacctgtatcctgccaacgagctgatcgccctgctggaggagaagggcatc<br>gtgttcagggatggctccaacatcctgccaaagctgctggagaatgacgattctcac<br>gccatcgacaccatggtggccctgatccgcagcgtgctgcagatgcggaactcca<br>atgccgccacaggcgaggactatatcaacagccccgtgcgcgatctgaatggcgt<br>gtgcttcgactcccggtttcagaacccagagtggcccatggacgccgatgccaatg<br>gcgcctaccacatcgccctgaagggccagctgctgctgaatcacctgaaggagag<br>caaggatctgaagctgcagaacggcatctccaatcaggactggctggcctacatcc<br>aggagctgcgcaacaaaaggccggcggccacgaaaaaggccggccaggcaaa<br>aaagaaaaaggaattcggcagtggagagggcagaggaagtctgctaacatgcg<br>gtgacgtcgaggagaatcctggcccagtgagcaagggcgaggagctgttcaccg<br>gggtggtgcccatcctggtcgagctggacggcgacgtaaacggccacaagttcag<br>cgtgtccggcgagggcgagggcgatgccacctacggcaagctgaccctgaagtt<br>catctgcaccaccggcaagctgcccgtgccctggcccaccctcgtgaccaccctgac<br>ctacggcgtgcagtgcttcagccgctaccccgaccacatgaagcagcacgacttctt<br>caagtccgccatgcccgaaggctacgtccaggagcgcaccatcttcttcaaggacg<br>acggcaactacaagacccgcgccgaggtgaagttcgagggcgacaccctggtga<br>accgcatcgagctgaagggcatcgacttcaaggaggacggcaacatcctggggc<br>acaagctggagtacaactacaacagccacaacgtctatatcatggccgacaagcag<br>aagaacggcatcaaggtgaacttcaagatccgccacaacatcgaggacggcagc<br>gtgcagctcgccgaccactaccagcagaacacccccatcggcgacggccccgtgc<br>tgctgcccgacaaccactacctgagcacccagtccgccctgagcaaagaccccaac<br>gagaagcgcgatcacatggtcctgctggagttcgtgaccgccgccgggatcactct<br>cggcatggacgagctgtacaaggaattctaactagagctcgctgatcagcctcgac<br>tgtgccttctagttgccagccatctgttgtttgcccctcccccgtgccttccttgaccctg<br>gaaggtgccactcccactgtcctttcctaataaaatgaggaaattgcatcgcattgtct<br>gagtaggtgtcattctattctggggggtggggtggggcaggacagcaagggggg<br>aggattgggaagagaatagcaggcatgctggggagcggccgcaggaacccta<br>gtgatggagttggccactccctctctgcgcgctcgctcgctcactgaggccgggcg<br>accaaaggtcgcccgacgcccgggctttgcccgggcggcctcagtgagcgagcg<br>agcgcgcagctgcctgcaggggcgcctgatgcggtattttctccttacgcatctgtg<br>cggtatttcacaccgcatacgtcaaagcaaccatagtacgcgccctgtagcggcgc<br>attaagcgcggcgggtgtggtggttacgcgcagcgtgaccgctacacttgccagc<br>gccttagcgcccgctcctttcgctttcttcccttcctttctcgccacgttcgccggctttcc<br>ccgtcaagctctaaatcgggggctccctttagggttccgatttagtgctttacggcac<br>ctcgaccccaaaaaacttgatttgggtgatggttcacgtagtgggccatcgccctga |

| | |
|---|---|
| | tagacggtttttcgccctttgacgttggagtccacgttctttaatagtggactcttgttcc<br>aaactggaacaacactcaactctatctcgggctattcttttgatttataagggattttgc<br>cgatttcggtctattggttaaaaaatgagctgatttaacaaaaatttaacgcgaatttta<br>acaaaatattaacgtttacaattttatggtgcactctcagtacaatctgctctgatgccg<br>catagttaagccagccccgacacccgccaacacccgctgacgcgccctgacgggc<br>ttgtctgctcccggcatccgcttacagacaagctgtgaccgtctccgggagctgcat<br>gtgtcagaggttttcaccgtcatcaccgaaacgcgcgagacgaaagggcctcgtg<br>atacgcctatttttataggttaatgtcatgataataatggtttcttagacgtcaggtggc<br>acttttcggggaaatgtgcgcggaacccctatttgtttatttttctaaatacattcaaata<br>tgtatccgctcatgagacaataaccctgataaatgcttcaataatattgaaaaaggaa<br>gagtatgagtattcaacatttccgtgtcgcccttattcccttttttgcggcattttgccttc<br>ctgtttttgctcacccagaaacgctggtgaaagtaaaagatgctgaagatcagttgg<br>gtgcacgagtgggttacatcgaactggatctcaacagcggtaagatccttgagagt<br>tttcgccccgaagaacgttttccaatgatgagcacttttaaagttctgctatgtggcgc<br>ggtattatcccgtattgacgccgggcaagagcaactcggtcgccgcatacactattc<br>tcagaatgacttggttgagtactcaccagtcacagaaaagcatcttacggatggcat<br>gacagtaagagaattatgcagtgctgccataaccatgagtgataacactgcggcca<br>acttacttctgacaacgatcggaggaccgaaggagctaaccgcttttttgcacaacat<br>gggggatcatgtaactcgccttgatcgttgggaaccggagctgaatgaagccatac<br>caaacgacgagcgtgacaccacgatgcctgtagcaatggcaacaacgttgcgcaa<br>actattaactggcgaactacttactctagcttcccggcaacaattaatagactggatg<br>gaggcggataaagttgcaggaccacttctgcgctcggcccttccggctggctggttt<br>attgctgataaatctggagccggtgagcgtggaagccgcggtatcattgcagcact<br>ggggccagatggtaagccctcccgtatcgtagttatctacacgacggggagtcag<br>gcaactatggatgaacgaaatagacagatcgctgagataggtgcctcactgattaa<br>gcattggtaactgtcagaccaagtttactcatatatactttagattgatttaaaacttcat<br>ttttaatttaaaaggatctaggtgaagatcctttttgataatctcatgaccaaaatccctt<br>aacgtgagttttcgttccactgagcgtcagaccccgtagaaaagatcaaaggatctt<br>cttgagatcctttttttctgcgcgtaatctgctgcttgcaaacaaaaaaaccaccgctac<br>cagcggtggtttgtttgccggatcaagagctaccaactctttttccgaaggtaactgg<br>cttcagcagagcgcagataccaaatactgttcttctagtgtagccgtagttaggccac<br>cacttcaagaactctgtagcaccgcctacatacctcgctctgctaatcctgttaccagt<br>ggctgctgccagtggcgataagtcgtgtcttaccgggttggactcaagacgatagtt<br>accggataaggcgcagcggtcgggctgaacggggggttcgtgcacacagccca<br>gcttggagcgaacgacctacaccgaactgagatacctacagcgtgagctatgaga<br>aagcgccacgcttcccgaagggagaaaggcggacaggtatccggtaagcggca<br>gggtcggaacaggagagcgcacgagggagcttccagggggaaacgcctggtat<br>ctttatagtcctgtcgggtttcgccacctctgacttgagcgtcgatttttgtgatgctcgt<br>caggggggcggagcctatggaaaaacgccagcaacgcggcctttttacggttcct<br>ggccttttgctggccttttgctcacatgt |
| **Eukaryote Expression Vector of BvCas12a-R** | |
| BvCas12a-<br>eukaryotes-<br>R-HDV-NC | gagggcctatttcccatgattccttcatatttgcatatacgatacaaggctgttagaga<br>gataattggaattaatttgactgtaaacacaaagatattagtacaaaatacgtgacgt<br>agaaagtaataatttcttgggtagtttgcagttttaaaattatgttttaaaatggactatc |

| | |
|---|---|
| | atatgcttaccgtaacttgaaagtatttcgatttcttggctttatatatcttgtggaaagg<br>acgaaacaccgAATTTCTACTATTGTAGATgggtcttcgagaagacctgg<br>ccggcatggtcccagcctcctcgctggcgccggctgggcaacatgcttcggcatgg<br>cgaatgggactttttttgttttagagctagaaatagcaagttaaaataaggctagtccg<br>tttttagcgcgtgcgccaattctgcagacaaatggctctagaggtacccgttacataa<br>cttacggtaaatggcccgcctggctgaccgcccaacgacccccgcccattgacgtc<br>aatagtaacgccaatagggactttccattgacgtcaatgggtggagtatttacggta<br>aactgcccacttggcagtacatcaagtgtatcatatgccaagtacgccccctattgac<br>gtcaatgacggtaaatggcccgcctggcattgtgcccagtacatgaccttatgggac<br>tttcctacttggcagtacatctacgtattagtcatcgctattaccatggtcgaggtgag<br>ccccacgttctgcttcactctccccatctcccccccctccccacccccaattttgtatttatt<br>tatttttaattattttgtgcagcgatgggggcgggggggggggggggcgcgcg<br>ccaggcggggcggggcggggcgaggggcggggcggggcgaggcggagag<br>gtgcggcggcagccaatcagagcggcgcgctccgaaagtttccttttatggcgag<br>gcggcggcggcggcggccctataaaaagcgaagcgcgcggcgggcgggagtc<br>gctgcgcgctgccttcgccccgtgccccgctccgccgccgcctcgcgccgcccgcc<br>ccggctctgactgaccgcgttactcccacaggtgagcgggcgggacggcccttctc<br>ctccgggctgtaattagctgagcaagaggtaagggtttaagggatggttggttggt<br>ggggtattaatgtttaattacctggagcacctgcctgaaatcactttttttcaggttgga<br>ccggtgccaccatggactataaggaccacgacggagactacaaggatcatgatatt<br>gattacaaagacgatgacgataagatggccccaaagaagaagcggaaggtcggt<br>atccacggagtcccagcagccATGCAGGAGAGAAAGAAGATCAGCC<br>ACCTGACCCACAGAAACAGCGTGAAGAAAACCATCAGAATGC<br>AGCTGAACCCCGTGGGAAAGACCATGGACTACTTCCAGGCCA<br>AGCAGATCCTGGAGAACGACGAGAAGCTGAAGGAGGACTAC<br>CAGAAGATCAAGGAGATCGCCGACAGATTCTACAGAAACCTG<br>AACGAGGACGTGCTGAGCAAAACCGGACTGGACAAGCTGAA<br>GGACTACGCCGAGATCTACTACCATTGCAACACCGACGCCGA<br>CAGAAAGAGACTGAACGAGTGCGCCAGCGAGCTGAGAAAGG<br>AGATCGTGAAGAACTTCAAGAACAGAGATGAGTACAACAAGC<br>TGTTCAACAAGAAGATGATCGAGATCGTGCTGCCCAAGCACCT<br>GAAGAACGAGGACGAGAAGGAAGTGGTGGCCAGCTTCAAGA<br>ACTTCACCACCTACTTCACCGGCTTCTTCACCAACAGAAAGAA<br>CATGTACAGCGACGGCGAAGAGTCTACCGCTATTGCCTACAG<br>ATGCATCAACGAGAACCTGCCCAAGCACCTGGACAACGTGAA<br>GGTGTTCGAGAAGGCCATCAGCAAGCTGAGCAAGAACGCCAT<br>CGACGACCTGGATGCCACATATTCTGGCCTGTGCGGCACAAAT<br>CTGTACGACGTGTTCACCGTGGACTACTTCAACTTCCTGCTGCC<br>CCAAAGCGGAATCACCGAGTACAACAAGATCATCGGCGGCTA<br>CACAACAAGCGACGGCACCAAAGTGAAGGGCATCAACGAGTA<br>CATCAACCTGTACAACCAGCAGGTGAGCAAGAGAGACAAGAT<br>CCCCAACCTGAAGATCCTGTACAAGCAGATCCTGAGCGAGAG<br>CGAGAAGGTGTCTTTCATCCCCCCCAAGTTCGAGGACGACAAC<br>GAACTGCTGTCTGCCGTGAGCGAGTTCTATGCCAACGACGAGA |

| | |
|---|---|
| | CATTTGATGGCATGCCCCTGAAGAAAGCCATCGACGAAACCA<br>AACTGCTGTTCGGCAACCTGGACAACAGCAGCCTGAACGGCA<br>TCTACATCCAGAACGACAGAAGCGTGACCAACCTGAGCAACA<br>GCATGTTCGGCAGCTGGAGCGTGATTGAGGACCTGTGGAACA<br>AGAACTACGACAGCGTGAACAGCAACAGCAGAATCAAGGACA<br>TCCAGAAGAGAGAGGACAAGAGAAAGAAGGCCTACAAGGCC<br>GAGAAGAAGCTGAGCCTGAGCTTCCTGCAGGTGCTGATCAGC<br>AACAGCGAGAACGACGAGATCAGAAAGAAGAGCATCGTGGA<br>CTACTACAAGACCAGCCTGATGCAGCTGACCGACAACCTGAG<br>CGACAAGTACAAAGAAGCCGCCCCCCTGTTTTCTGAGAACTAC<br>GACAACGAGAAGGGCCTGAAGAACGACGACAAGAGCATCAG<br>CCTGATCAAGAACTTCCTGGACGCCATCAAGGAGATCGAGAA<br>GTTCATCAAGCCCCTGAGCGAGACAAATATCACCGGCGAGAA<br>GAACGACCTGTTCTACAGCCAGTTCACCCCCCTGCTGGACAAC<br>ATCAGCAGAATCGACAGACTGTACGACAAGGTGAGAAACTAC<br>GTGACCCAGAAGCCCTTCAGCACCGACAAGATCAAGCTGAAC<br>TTCGGCAACAGCCAGCTTCTGagaGGCTGGGACAGAAACAAG<br>GAGAAGGACTGTGGCGCTGTGCTGCTGTGTAAGGACGAGAAG<br>TACTACCTGGCCATCATCGACAAGAGCAACAACAGCATCCTGG<br>AGAACATCGACTTCCAGGACTGCAACGAGAGCGACTACTACG<br>AGAAGATCGTGTACAAGCTGCTGcccAAGATCTCTGGCAACCT<br>GCCCAGAGTGTTCTTCAGCGAGAAGCACAAGAAGCTGCTGAG<br>CCCCAGCGATGAGATCCTGAAGATCTACAAGAGCGGCACCTT<br>CAAGAAGGGCGACAAGTTCAGCCTTGACGACTGCCACAAGCT<br>GATCGACTTCTACAAGGAGAGCTTCAAGAAGTACCCCAAGTG<br>GCTGATCTACAACTTCAAGTTCAAGAACACCAACGAGTACAAC<br>GACATCAGCGAGTTCTACAACGACGTGGCCAGCCAGGGATAC<br>AACATCAGCAAGATGAAGATCCCCACCAGCTTCATCGACAAG<br>CTGGTGGACGAGGGCAAGATCTACCTGTTCCAGCTGTACAACA<br>AGGACTTCAGCCCCCACAGCAAGGGAACACCTAACCTGCACA<br>CCCTGTACTTCAAGATGCTGTTCGACGAGAGAAACCTGGAGGA<br>CGTGGTGTACAAGCTGAATGGCGAGGCCGAGATGTTTTACAG<br>ACCCGCCAGCATCAAGTATGACAAGCCCACCCACCCTAAGAA<br>CACCCCCATCAAGAACAAGAACACCCTGAACGACAAGAAGGC<br>CAGCACCTTCCCCTACGACCTGATCAAGGACAAGAGATACAC<br>CAAGTGGCAGTTCAGCCTGCACTTCCCCATCACCATGAACTTC<br>AAGGCCCCCGACAGAGCCATGATCAACGACGACGTGAGAAAC<br>CTGCTGAAGAGCTGCAACAACAACTTCATCATCGGCATCGACA<br>GAGGCGAGAGAAACCTGCTGTACGTGAGCGTGATCGATAGCA<br>ACGGCGCCATCATCTACCAGCACAGCCTGAACATCATCGGCA<br>ACAAGTTCAAGGGCAAGACCTACGAAACCAACTACAGAGAGA<br>AGCTGGCCACCAGAGAGAAGGAGAGAACCGAGCAGAGAAGA<br>AACTGGAAGGCCATCGAGAGCATCAAGGAGCTGAAGGAGGG<br>CTACATCAGCCAAACCGTGCACGTGATTTGCCAGCTGGTGGTG |

| | |
|---|---|
| | AAGTACGACGCCATCATCGTGATGGAGAAGCTGACCGACGGC<br>TTCAAGAGAGGCAGAACCAAGTTCGAGAAGCAGGTGTACCAG<br>AAGTTCGAGAAGATGCTGATCGACAAGCTGAACTACTACGTGG<br>ACAAGAAGCTGGACCCCAATGAGGAAGGCGGACTGCTGCATG<br>CTTATCAGCTGACCAACAAGCTGGACAGCTTCGACAAGCTGG<br>GAATGCAGAGCGGCTTCATCTTCTACGTCAGACCCGACTTCAC<br>CAGCAAAATCGACCCCGTGACCGGATTTGTGAACCTGCTGTAC<br>CCCAGATACGAGAACATCGACAAGGCCAAGGACATGATCAGC<br>AGATTCGACGACATCAGATACAACGCCGGCGAGGACTTCTTC<br>GAGTTCGACATCGACTACGACAAGTTCCCCAAGACCGCCAGC<br>GACTACAGAAAGAAGTGGACCATCTGCACCAACGGCGAGAGA<br>ATCGAGGCCTTCAGAAACCCCGCCAACAACAACGAGTGGAGC<br>TACAGAACCATCATCCTGGCCGAGAAGTTCAAGGAGCTGTTCG<br>ACAACAACAGCATCAACTACAGAGACAGCGACGACCTGAAAG<br>CCGAGATCCTGAGCCAAACCAAGGGCAAGTTCTTCGAGGACT<br>TCTTCAAGCTGCTGAGACTGACCCTGCAGATGAGAAACAGCAA<br>CCCCGAAACCGGAGAGGACAGGATTCTGAGCCCCGTGAAGGA<br>CAAGAACGGCAACTTCTACGACAGCAGCAAGTACGACGAGAA<br>GAGCAAGCTGCCCTGTGACGCTGATGCTAACGGCGCTTACAA<br>CATCGCCAGAAAGGGCCTGTGGATCGTGGAGCAGTTCAAGAA<br>GGCCGACAACGTGTCTGCTGTGGAACCCGTGATCCACAACGA<br>CAAGTGGCTGAAGTTCGTGCAGGAGAACGACATGGCCAACAA<br>Caaaaggccggcggccacgaaaaaggccggccaggcaaaaaagaaaaagga<br>attcggcagtggagagggcagaggaagtctgctaacatgcggtgacgtcgagga<br>gaatcctggcccagtgagcaagggcgaggagctgttcaccggggtggtgcccat<br>cctggtcgagctggacggcgacgtaaacggccacaagttcagcgtgtccggcga<br>gggcgagggcgatgccacctacggcaagctgaccctgaagttcatctgcaccacc<br>ggcaagctgcccgtgccctggcccaccctcgtgaccaccctgacctacggcgtgca<br>gtgcttcagccgctaccccgaccacatgaagcagcacgacttcttcaagtccgccat<br>gcccgaaggctacgtccaggagcgcaccatcttcttcaaggacgacggcaactac<br>aagacccgcgccgaggtgaagttcgagggcgacaccctggtgaaccgcatcgag<br>ctgaagggcatcgacttcaaggaggacggcaacatcctggggcacaagctggag<br>tacaactacaacagccacaacgtctatatcatggccgacaagcagaagaacggcat<br>caaggtgaacttcaagatccgccacaacatcgaggacggcagcgtgcagctcgcc<br>gaccactaccagcagaacacccccatcggcgacggccccgtgctgctgcccgaca<br>accactacctgagcacccagtccgccctgagcaaagaccccaacgagaagcgcga<br>tcacatggtcctgctggagttcgtgaccgccgccgggatcactctcggcatggacg<br>agctgtacaaggaattctaactagagctcgctgatcagcctcgactgtgccttctagt<br>tgccagccatctgttgtttgcccctcccccgtgccttccttgaccctggaaggtgccac<br>tcccactgtcctttcctaataaaatgaggaaattgcatcgcattgtctgagtaggtgtc<br>attctattctggggggtggggtggggcaggacagcaagggggaggattgggaa<br>gagaatagcaggcatgctggggagcggccgcaggaacccctagtgatggagtt<br>ggccactccctctctgcgcgctcgctcgctcactgaggccgggcgaccaaaggtcg<br>cccgacgcccgggctttgcccgggcggcctcagtgagcgagcgagcgcgcagct |

| | |
|---|---|
| | gcctgcaggggcgcctgatgcggtattttctccttacgcatctgtgcggtatttcacac<br>cgcatacgtcaaagcaaccatagtacgcgccctgtagcggcgcattaagcgcggc<br>gggtgtggtggttacgcgcagcgtgaccgctacacttgccagcgccttagcgcccg<br>ctcctttcgctttcttcccttcctttctcgccacgttcgccggctttccccgtcaagctctaa<br>atcgggggctccctttagggttccgatttagtgctttacggcacctcgaccccaaaaa<br>acttgatttgggtgatggttcacgtagtgggccatcgccctgatagacggtttttcgc<br>cctttgacgttggagtccacgttctttaatagtggactcttgttccaaactggaacaac<br>actcaactctatctcgggctattcttttgatttataagggattttgccgatttcggtctatt<br>ggttaaaaaatgagctgatttaacaaaaatttaacgcgaattttaacaaaatattaac<br>gtttacaattttatggtgcactctcagtacaatctgctctgatgccgcatagttaagcca<br>gccccgacacccgccaacacccgctgacgcgccctgacgggcttgtctgctcccgg<br>catccgcttacagacaagctgtgaccgtctccgggagctgcatgtgtcagaggtttt<br>caccgtcatcaccgaaacgcgcgagacgaaagggcctcgtgatacgcctatttttat<br>aggttaatgtcatgataataatggtttcttagacgtcaggtggcacttttcggggaaa<br>tgtgcgcggaacccctatttgtttatttttctaaatacattcaaatatgtatccgctcatg<br>agacaataaccctgataaatgcttcaataatattgaaaaaggaagagtatgagtatt<br>caacatttccgtgtcgcccttattcccttttttgcggcattttgccttcctgtttttgctcacc<br>cagaaacgctggtgaaagtaaaagatgctgaagatcagttgggtgcacgagtgg<br>gttacatcgaactggatctcaacagcggtaagatccttgagagttttcgccccgaag<br>aacgttttccaatgatgagcacttttaaagttctgctatgtggcgcggtattatcccgta<br>ttgacgccgggcaagagcaactcggtcgccgcatacactattctcagaatgacttg<br>gttgagtactcaccagtcacagaaaagcatcttacggatggcatgacagtaagaga<br>attatgcagtgctgccataaccatgagtgataacactgcggccaacttacttctgaca<br>acgatcggaggaccgaaggagctaaccgcttttttgcacaacatgggggatcatgt<br>aactcgccttgatcgttgggaaccggagctgaatgaagccataccaaacgacgag<br>cgtgacaccacgatgcctgtagcaatggcaacaacgttgcgcaaactattaactgg<br>cgaactacttactctagcttcccggcaacaattaatagactggatggaggcggataa<br>agttgcaggaccacttctgcgctcggcccttccggctggctggtttattgctgataaa<br>tctggagccggtgagcgtggaagccgcggtatcattgcagcactggggccagat<br>ggtaagccctcccgtatcgtagttatctacacgacggggagtcaggcaactatgga<br>tgaacgaaatagacagatcgctgagataggtgcctcactgattaagcattggtaact<br>gtcagaccaagtttactcatatatactttagattgatttaaaacttcatttttaatttaaaa<br>ggatctaggtgaagatcctttttgataatctcatgaccaaaatcccttaacgtgagtttt<br>cgttccactgagcgtcagaccccgtagaaaagatcaaaggatcttcttgagatccttt<br>ttttctgcgcgtaatctgctgcttgcaaacaaaaaaaccaccgctaccagcggtggtt<br>tgtttgccggatcaagagctaccaactctttttccgaaggtaactggcttcagcagag<br>cgcagataccaaatactgttcttctagtgtagccgtagttaggccaccacttcaagaa<br>ctctgtagcaccgcctacatacctcgctctgctaatcctgttaccagtggctgctgcca<br>gtggcgataagtcgtgtcttaccgggttggactcaagacgatagttaccggataag<br>gcgcagcggtcgggctgaacggggggttcgtgcacacagcccagcttggagcg<br>aacgacctacaccgaactgagatacctacagcgtgagctatgagaaagcgccacg<br>cttcccgaagggagaaaggcggacaggtatccggtaagcggcagggtcggaac<br>aggagagcgcacgagggagcttccagggggaaacgcctggtatctttatagtcct<br>gtcgggtttcgccacctctgacttgagcgtcgatttttgtgatgctcgtcaggggggc |

| | |
|---|---|
| | ggagcctatggaaaaacgccagcaacgcggcctttttacggttcctggccttttgct<br>ggccttttgctcacatgt |
| **Eukaryote Expression Vector of BvCas12a-RVR** | |
| BvCas12a-<br>eukaryotes-<br>RVR-HDV-<br>NC | gagggcctatttcccatgattccttcatatttgcatatacgatacaaggctgttagaga<br>gataattggaattaatttgactgtaaacacaaagatattagtacaaaatacgtgacgt<br>agaaagtaataatttcttgggtagtttgcagttttaaaattatgttttaaaatggactatc<br>atatgcttaccgtaacttgaaagtatttcgatttcttggctttatatatcttgtggaaagg<br>acgaaacaccgAATTTCTACTATTGTAGATgggtcttcgagaagacctgg<br>ccggcatggtcccagcctcctcgctggcgccggctgggcaacatgcttcggcatgg<br>cgaatgggactttttgttttagagctagaaatagcaagttaaaataaggctagtccg<br>tttttagcgcgtgcgccaattctgcagacaaatggctctagaggtacccgttacataa<br>cttacggtaaatggcccgcctggctgaccgcccaacgacccccgcccattgacgtc<br>aatagtaacgccaatagggactttccattgacgtcaatgggtggagtatttacggta<br>aactgcccacttggcagtacatcaagtgtatcatatgccaagtacgccccctattgac<br>gtcaatgacggtaaatggcccgcctggcattgtgcccagtacatgaccttatgggac<br>tttcctacttggcagtacatctacgtattagtcatcgctattaccatggtcgaggtgag<br>ccccacgttctgcttcactctccccatctcccccccctccccacccccaattttgtatttatt<br>tattttttaattattttgtgcagcgatgggggcgggggggggggggggcgcgcg<br>ccaggcggggcggggcggggcgaggggcggggcggggcgaggcggagag<br>gtgcggcggcagccaatcagagcggcgcgctccgaaagtttccttttatggcgag<br>gcggcggcggcggcggccctataaaaagcgaagcgcgcggcgggcgggagtc<br>gctgcgcgctgccttcgccccgtgccccgctccgccgccgcctcgcgccgcccgcc<br>ccggctctgactgaccgcgttactcccacaggtgagcgggcgggacggcccttctc<br>ctccgggctgtaattagctgagcaagaggtaagggtttaagggatggttggttggt<br>ggggtattaatgtttaattacctggagcacctgcctgaaatcactttttttcaggttgga<br>ccggtgccaccatggactataaggaccacgacggagactacaaggatcatgatatt<br>gattacaaagacgatgacgataagatggccccaaagaagaagcggaaggtcggt<br>atccacggagtcccagcagccATGCAGGAGAGAAAGAAGATCAGCC<br>ACCTGACCCACAGAAACAGCGTGAAGAAAACCATCAGAATGC<br>AGCTGAACCCCGTGGGAAAGACCATGGACTACTTCCAGGCCA<br>AGCAGATCCTGGAGAACGACGAGAAGCTGAAGGAGGACTAC<br>CAGAAGATCAAGGAGATCGCCGACAGATTCTACAGAAACCTG<br>AACGAGGACGTGCTGAGCAAAACCGGACTGGACAAGCTGAA<br>GGACTACGCCGAGATCTACTACCATTGCAACACCGACGCCGA<br>CAGAAAGAGACTGAACGAGTGCGCCAGCGAGCTGAGAAAGG<br>AGATCGTGAAGAACTTCAAGAACAGAGATGAGTACAACAAGC<br>TGTTCAACAAGAAGATGATCGAGATCGTGCTGCCCAAGCACCT<br>GAAGAACGAGGACGAGAAGGAAGTGGTGGCCAGCTTCAAGA<br>ACTTCACCACCTACTTCACCGGCTTCTTCACCAACAGAAAGAA<br>CATGTACAGCGACGGCGAAGAGTCTACCGCTATTGCCTACAG<br>ATGCATCAACGAGAACCTGCCCAAGCACCTGGACAACGTGAA<br>GGTGTTCGAGAAGGCCATCAGCAAGCTGAGCAAGAACGCCAT<br>CGACGACCTGGATGCCACATATTCTGGCCTGTGCGGCACAAAT<br>CTGTACGACGTGTTCACCGTGGACTACTTCAACTTCCTGCTGCC |

| | |
|---|---|
| | CCAAAGCGGAATCACCGAGTACAACAAGATCATCGGCGGCTA<br>CACAACAAGCGACGGCACCAAAGTGAAGGGCATCAACGAGTA<br>CATCAACCTGTACAACCAGCAGGTGAGCAAGAGAGACAAGAT<br>CCCCAACCTGAAGATCCTGTACAAGCAGATCCTGAGCGAGAG<br>CGAGAAGGTGTCTTTCATCCCCCCCAAGTTCGAGGACGACAAC<br>GAACTGCTGTCTGCCGTGAGCGAGTTCTATGCCAACGACGAGA<br>CATTTGATGGCATGCCCCTGAAGAAAGCCATCGACGAAACCA<br>AACTGCTGTTCGGCAACCTGGACAACAGCAGCCTGAACGGCA<br>TCTACATCCAGAACGACAGAAGCGTGACCAACCTGAGCAACA<br>GCATGTTCGGCAGCTGGAGCGTGATTGAGGACCTGTGGAACA<br>AGAACTACGACAGCGTGAACAGCAACAGCAGAATCAAGGACA<br>TCCAGAAGAGAGAGGACAAGAGAAAGAAGGCCTACAAGGCC<br>GAGAAGAAGCTGAGCCTGAGCTTCCTGCAGGTGCTGATCAGC<br>AACAGCGAGAACGACGAGATCAGAAAGAAGAGCATCGTGGA<br>CTACTACAAGACCAGCCTGATGCAGCTGACCGACAACCTGAG<br>CGACAAGTACAAAGAAGCCGCCCCCCTGTTTTCTGAGAACTAC<br>GACAACGAGAAGGGCCTGAAGAACGACGACAAGAGCATCAG<br>CCTGATCAAGAACTTCCTGGACGCCATCAAGGAGATCGAGAA<br>GTTCATCAAGCCCCTGAGCGAGACAAATATCACCGGCGAGAA<br>GAACGACCTGTTCTACAGCCAGTTCACCCCCCTGCTGGACAAC<br>ATCAGCAGAATCGACAGACTGTACGACAAGGTGAGAAACTAC<br>GTGACCCAGAAGCCCTTCAGCACCGACAAGATCAAGCTGAAC<br>TTCGGCAACAGCCAGCTTCTGagaGGCTGGGACAGAAACgtgG<br>AGAAGGACagaGGCGCTGTGCTGCTGTGTAAGGACGAGAAGT<br>ACTACCTGGCCATCATCGACAAGAGCAACAACAGCATCCTGG<br>AGAACATCGACTTCCAGGACTGCAACGAGAGCGACTACTACG<br>AGAAGATCGTGTACAAGCTGCTGcccAAGATCTCTGGCAACCT<br>GCCCAGAGTGTTCTTCAGCGAGAAGCACAAGAAGCTGCTGAG<br>CCCCAGCGATGAGATCCTGAAGATCTACAAGAGCGGCACCTT<br>CAAGAAGGGCGACAAGTTCAGCCTTGACGACTGCCACAAGCT<br>GATCGACTTCTACAAGGAGAGCTTCAAGAAGTACCCCAAGTG<br>GCTGATCTACAACTTCAAGTTCAAGAACACCAACGAGTACAAC<br>GACATCAGCGAGTTCTACAACGACGTGGCCAGCCAGGGATAC<br>AACATCAGCAAGATGAAGATCCCCACCAGCTTCATCGACAAG<br>CTGGTGGACGAGGGCAAGATCTACCTGTTCCAGCTGTACAACA<br>AGGACTTCAGCCCCCACAGCAAGGGAACACCTAACCTGCACA<br>CCCTGTACTTCAAGATGCTGTTCGACGAGAGAAACCTGGAGGA<br>CGTGGTGTACAAGCTGAATGGCGAGGCCGAGATGTTTTACAG<br>ACCCGCCAGCATCAAGTATGACAAGCCCACCCACCCTAAGAA<br>CACCCCCATCAAGAACAAGAACACCCTGAACGACAAGAAGGC<br>CAGCACCTTCCCCTACGACCTGATCAAGGACAAGAGATACAC<br>CAAGTGGCAGTTCAGCCTGCACTTCCCCATCACCATGAACTTC<br>AAGGCCCCCGACAGAGCCATGATCAACGACGACGTGAGAAAC<br>CTGCTGAAGAGCTGCAACAACAACTTCATCATCGGCATCGACA |

| | |
|---|---|
| | GAGGCGAGAGAAACCTGCTGTACGTGAGCGTGATCGATAGCA<br>ACGGCGCCATCATCTACCAGCACAGCCTGAACATCATCGGCA<br>ACAAGTTCAAGGGCAAGACCTACGAAACCAACTACAGAGAGA<br>AGCTGGCCACCAGAGAGAAGGAGAGAACCGAGCAGAGAAGA<br>AACTGGAAGGCCATCGAGAGCATCAAGGAGCTGAAGGAGGG<br>CTACATCAGCCAAACCGTGCACGTGATTTGCCAGCTGGTGGTG<br>AAGTACGACGCCATCATCGTGATGGAGAAGCTGACCGACGGC<br>TTCAAGAGAGGCAGAACCAAGTTCGAGAAGCAGGTGTACCAG<br>AAGTTCGAGAAGATGCTGATCGACAAGCTGAACTACTACGTGG<br>ACAAGAAGCTGGACCCCAATGAGGAAGGCGGACTGCTGCATG<br>CTTATCAGCTGACCAACAAGCTGGACAGCTTCGACAAGCTGG<br>GAATGCAGAGCGGCTTCATCTTCTACGTCAGACCCGACTTCAC<br>CAGCAAAATCGACCCCGTGACCGGATTTGTGAACCTGCTGTAC<br>CCCAGATACGAGAACATCGACAAGGCCAAGGACATGATCAGC<br>AGATTCGACGACATCAGATACAACGCCGGCGAGGACTTCTTC<br>GAGTTCGACATCGACTACGACAAGTTCCCCAAGACCGCCAGC<br>GACTACAGAAAGAAGTGGACCATCTGCACCAACGGCGAGAGA<br>ATCGAGGCCTTCAGAAACCCCGCCAACAACAACGAGTGGAGC<br>TACAGAACCATCATCCTGGCCGAGAAGTTCAAGGAGCTGTTCG<br>ACAACAACAGCATCAACTACAGAGACAGCGACGACCTGAAAG<br>CCGAGATCCTGAGCCAAACCAAGGGCAAGTTCTTCGAGGACT<br>TCTTCAAGCTGCTGAGACTGACCCTGCAGATGAGAAACAGCAA<br>CCCCGAAACCGGAGAGGACAGGATTCTGAGCCCCGTGAAGGA<br>CAAGAACGGCAACTTCTACGACAGCAGCAAGTACGACGAGAA<br>GAGCAAGCTGCCCTGTGACGCTGATGCTAACGGCGCTTACAA<br>CATCGCCAGAAAGGGCCTGTGGATCGTGGAGCAGTTCAAGAA<br>GGCCGACAACGTGTCTGCTGTGGAACCCGTGATCCACAACGA<br>CAAGTGGCTGAAGTTCGTGCAGGAGAACGACATGGCCAACAA<br>Caaaaggccggcggccacgaaaaaggccggccaggcaaaaaagaaaaagga<br>attcggcagtggagagggcagaggaagtctgctaacatgcggtgacgtcgagga<br>gaatcctggcccagtgagcaagggcgaggagctgttcaccggggtggtgcccat<br>cctggtcgagctggacggcgacgtaaacggccacaagttcagcgtgtccggcga<br>gggcgagggcgatgccacctacggcaagctgaccctgaagttcatctgcaccacc<br>ggcaagctgcccgtgccctggcccaccctcgtgaccaccctgacctacggcgtgca<br>gtgcttcagccgctaccccgaccacatgaagcagcacgacttcttcaagtccgccat<br>gcccgaaggctacgtccaggagcgcaccatcttcttcaaggacgacggcaactac<br>aagacccgcgccgaggtgaagttcgagggcgacaccctggtgaaccgcatcgag<br>ctgaagggcatcgacttcaaggaggacggcaacatcctggggcacaagctggag<br>tacaactacaacagccacaacgtctatatcatggccgacaagcagaagaacggcat<br>caaggtgaacttcaagatccgccacaacatcgaggacggcagcgtgcagctcgcc<br>gaccactaccagcagaacacccccatcggcgacggccccgtgctgctgcccgaca<br>accactacctgagcacccagtccgccctgagcaaagaccccaacgagaagcgcga<br>tcacatggtcctgctggagttcgtgaccgccgccgggatcactctcggcatggacg<br>agctgtacaaggaattctaactagagctcgctgatcagcctcgactgtgccttctagt |

| | |
|---|---|
| | tgccagccatctgttgtttgcccctcccccgtgccttccttgaccctggaaggtgccac<br>tcccactgtcctttcctaataaaatgaggaaattgcatcgcattgtctgagtaggtgtc<br>attctattctggggggtggggtggggcaggacagcaagggggaggattgggaa<br>gagaatagcaggcatgctggggagcggccgcaggaacccctagtgatggagtt<br>ggccactccctctctgcgcgctcgctcgctcactgaggccgggcgaccaaaggtcg<br>cccgacgcccgggctttgcccgggcggcctcagtgagcgagcgagcgcgcagct<br>gcctgcaggggcgcctgatgcggtattttctccttacgcatctgtgcggtatttcacac<br>cgcatacgtcaaagcaaccatagtacgcgccctgtagcggcgcattaagcgcggc<br>gggtgtggtggttacgcgcagcgtgaccgctacacttgccagcgccttagcgcccg<br>ctcctttcgctttcttcccttcctttctcgccacgttcgccggctttccccgtcaagctctaa<br>atcgggggctccctttagggttccgatttagtgctttacggcacctcgaccccaaaaa<br>acttgatttgggtgatggttcacgtagtgggccatcgccctgatagacggtttttcgc<br>cctttgacgttggagtccacgttctttaatagtggactcttgttccaaactggaacaac<br>actcaactctatctcgggctattcttttgatttataagggattttgccgatttcggtctatt<br>ggttaaaaaatgagctgatttaacaaaaatttaacgcgaattttaacaaaatattaac<br>gtttacaattttatggtgcactctcagtacaatctgctctgatgccgcatagttaagcca<br>gccccgacacccgccaacacccgctgacgcgccctgacgggcttgtctgctcccgg<br>catccgcttacagacaagctgtgaccgtctccgggagctgcatgtgtcagaggtttt<br>caccgtcatcaccgaaacgcgcgagacgaaagggcctcgtgatacgcctatttttat<br>aggttaatgtcatgataataatggtttcttagacgtcaggtggcacttttcggggaaa<br>tgtgcgcggaacccctatttgtttatttttctaaatacattcaaatatgtatccgctcatg<br>agacaataaccctgataaatgcttcaataatattgaaaaaggaagagtatgagtatt<br>caacatttccgtgtcgcccttattcccttttttgcggcattttgccttcctgtttttgctcacc<br>cagaaacgctggtgaaagtaaaagatgctgaagatcagttgggtgcacgagtgg<br>gttacatcgaactggatctcaacagcggtaagatccttgagagttttcgccccgaag<br>aacgttttccaatgatgagcacttttaaagttctgctatgtggcgcggtattatcccgta<br>ttgacgccgggcaagagcaactcggtcgccgcatacactattctcagaatgacttg<br>gttgagtactcaccagtcacagaaaagcatcttacggatggcatgacagtaagaga<br>attatgcagtgctgccataaccatgagtgataacactgcggccaacttacttctgaca<br>acgatcggaggaccgaaggagctaaccgcttttttgcacaacatgggggatcatgt<br>aactcgccttgatcgttgggaaccggagctgaatgaagccataccaaacgacgag<br>cgtgacaccacgatgcctgtagcaatggcaacaacgttgcgcaaactattaactgg<br>cgaactacttactctagcttcccggcaacaattaatagactggatggaggcggataa<br>agttgcaggaccacttctgcgctcggcccttccggctggctggtttattgctgataaa<br>tctggagccggtgagcgtggaagccgcggtatcattgcagcactggggccagat<br>ggtaagccctcccgtatcgtagttatctacacgacggggagtcaggcaactatgga<br>tgaacgaaatagacagatcgctgagataggtgcctcactgattaagcattggtaact<br>gtcagaccaagtttactcatatatactttagattgatttaaaacttcatttttaatttaaaa<br>ggatctaggtgaagatcctttttgataatctcatgaccaaaatcccttaacgtgagtttt<br>cgttccactgagcgtcagaccccgtagaaaagatcaaaggatcttcttgagatccttt<br>ttttctgcgcgtaatctgctgcttgcaaacaaaaaaaccaccgctaccagcggtggtt<br>tgtttgccggatcaagagctaccaactctttttccgaaggtaactggcttcagcagag<br>cgcagataccaaatactgttcttctagtgtagccgtagttaggccaccacttcaagaa<br>ctctgtagcaccgcctacatacctcgctctgctaatcctgttaccagtggctgctgcca |

| | |
|---|---|
| | gtggcgataagtcgtgtcttaccgggttggactcaagacgatagttaccggataag<br>gcgcagcggtcgggctgaacggggggttcgtgcacacagcccagcttggagcg<br>aacgacctacaccgaactgagatacctacagcgtgagctatgagaaagcgccacg<br>cttcccgaagggagaaaggcggacaggtatccggtaagcggcagggtcggaac<br>aggagagcgcacgagggagcttccagggggaaacgcctggtatctttatagtcct<br>gtcgggtttcgccacctctgacttgagcgtcgatttttgtgatgctcgtcaggggggc<br>ggagcctatggaaaaacgccagcaacgcggcctttttacggttcctggccttttgct<br>ggccttttgctcacatgt |
